# A predictive mechanochemical modeling framework for the deformation and remodeling of the nuclear lamina

**DOI:** 10.64898/2026.02.19.706840

**Authors:** Emmet A. Francis, Aanya Sawhney, Einollah Sarikhani, Hamed Naghsh-Nilchi, Dhivya Pushpa Meganathan, Zeinab Jahed, Padmini Rangamani

## Abstract

Nuclear envelope (NE) deformation and rupture are common in cell spreading and migration in a variety of microenvironments and lead to marked changes in nucleocytoplasmic transport. Predicting responses to different mechanochemical cues that are transmitted to the nucleus remains an open problem in the field of mechanomedicine. We have developed a predictive modeling framework to examine how nuclear deformation on substrates with different nanotopographies influences nucleocytoplasmic transport and rearrangement of the nuclear lamina. Using the finite element method, we simulated nuclear compression by the perinuclear actin cap on substrates with arrays of nanopillars, modeling the NE-lamin composite as a nonlinear elastic structure and coupling deformations to a biochemical model of lamin remodeling and nucleocytoplasmic transport. Our simulations generated nuclear morphologies that agreed well with our experimental measurements and predicted that the regions of highest NE stretch are adjacent to cell-nanopillar contacts, leading to maximized NE tension on nanopillars spaced by 5-7 microns. We then considered the effects on transport of YAP/TAZ and remodeling of the nuclear lamina. In our model, increased nuclear flattening facilitated YAP/TAZ nuclear localization due to NPC dilation. We validated these predictions by measuring nuclear YAP and lamin localization in subsequent experiments. Finally, we considered the implications of our model for NE rupture. We found that substrate nanotopography and lamin abundance were both strong predictors of the force load in the nuclear lamina and the resulting rupture likelihood. We validated this prediction in subsequent experiments with lamin-depleted U2OS cells, establishing the central importance of lamin transport and microenvironment nanotopography in nuclear mechanotransduction.

**Significance:** Cell nuclei experience large strains, but the biophysical effects of such deformations on nucleocytoplasmic transport and nuclear envelope rupture remain incompletely understood. Here, we present a modeling framework that includes the mechanics of nuclear deformation and the resulting effects on molecular transport. As a well-controlled setup for comparing experiments and simulations, we consider nuclear indentations exhibited by cells on nanopillar substrates. Our simulations predicted a correlation between nuclear YAP/TAZ and nuclear flattening and predicted that low-lamin cells experience higher force loads in the nuclear lamina. We validated these predictions experimentally, showing that flattened nuclei have higher nuclear YAP and lamin-depleted cells are more likely to exhibit nuclear rupture. Overall, our framework presents opportunities to predict nuclear mechanoadaptation to different microenvironments.

## Introduction

Cells respond to external biochemical and mechanical cues via shape changes and induction of intracellular signaling events. Changes in cell shape also affect the organization of intracellular structures such as the endoplasmic reticulum or mitochondria, as evidenced by dynamic measurements of organelle localization on substrates with different nanoscale features [1, 2]. Among organelles, the nucleus exhibits an especially dramatic response, as nuclear deformation is required for cell navigation through complex microenvironments [3, 4, 5, 6, 7]. In the extreme case of extravasation, immune cells or cancer cells must squeeze their nuclei between endothelial cells when migrating from blood vessels into surrounding tissue [8, 9]. Dissecting the forces required for organelle deformation and the implications for cellular state remains a pressing topic in the emerging field of mechanomedicine [10, 11].

Nuclear deformation acts as a key decision-making point in mechanotransduction. For instance, dilation of nuclear pore complexes (NPCs) in the nuclear envelope (NE) [12, 13, 14, 15] and/or volumetric compression of the nucleus [16] have been shown to accelerate nuclear entry of YAP/TAZ and other molecules such as TWIST1. Furthermore, nuclear stretch leads to NE rupture (NER) in cells squeezing through small channels [17, 18, 3, 7] or spreading on nanopillar substrates [19, 20]. Intermediate filaments known as lamins form a meshwork that makes up a significant portion of the layer beneath the NE double lipid bilayer and control NE stretch via interactions with the cytoskeleton mediated by the Linker of the Nucleoskeleton and Cytoskeleton (LINC) complex [21, 22, 23]. A-type lamins (lamin A/C) are particularly important determinants of nuclear rigidity [24, 25, 26] and the likelihood of NER [18, 14, 27]. The importance of a robust NE is clear from the many health problems associated with laminopathies, diseases defined by mutations in the gene for lamin A/C (LMNA) [28, 29]. Accordingly, it is crucial to refine our quantitative understanding of how lamin content and organization regulate nuclear mechanotransduction.

Engineered substrates with nanoscale topographical features have offered a powerful experimental tool to probe the effects of nanotopography on the nucleus [30, 31, 19, 20, 32]. Features such as nanopillars or nanoneedles introduce well-controlled, reproducible nuclear indentations, facilitating the study of nucleoskeletal and cytoskeletal remodeling adjacent to induced NE curvature [32, 30, 20, 19] (Figure 1A-B). These experiments have shown that F-actin and lamin A/C localize adjacent to cell and nuclear indentations [33, 32, 20]. In addition to cytoskeletal activation due to curvature of the plasma membrane [33, 34], NE curvature modulates arrangement of the lamin nucleoskeleton [18, 32]. However, the mechanisms coupling NE deformation to lamin remodeling remain unclear.

Quantitative modeling of NE deformation has served as a robust complementary approach to experimental techniques. Many studies use continuum models that describe the NE and lamina as an elastic shell with finite thickness [35, 36, 37, 38, 39, 8] or as an infinitesimally thin layer, analogous to the boundary of a liquid droplet [14, 40, 41, 42]. These continuum models facilitate predictions of nuclear morphology as a function of applied stress, e.g., due to micropipette aspiration [38], nuclear squeezing during migration [8, 40], or compression by the actin cytoskeleton [39, 35]. Although the mechanical properties of the nucleus have been characterized in some detail, coupling nuclear deformations to biochemical transport via advection and diffusion remains an open problem. Approaches to simulate mechanochemical coupling in the nucleus are required to answer several questions. For instance, how does rearrangement of lamins within the NE depend on the nanoscale topography of the microenvironment? Furthermore, in what ways do lamin remodeling and NPC dilation together determine changes in nucleocytoplasmic transport? Advances in three-dimensional simulations of reaction-transport systems and moving boundary problems with high fidelity present opportunities to examine coupled mechanochemical problems [43, 44, 45, 40, 46]. Thus, we are now well-positioned to simulate the effects of deformation on nucleocytoplasmic transport and lamin distribution.

In this work, we hypothesize that nuclear compression and indentation lead to lamin A/C rearrangement within the NE, causing increased YAP/TAZ influx and increased likelihood of NER upon deformation. To test this hypothesis, we developed a coupled mechanochemical computational framework to examine the effects of nanotopography-induced nuclear deformation on nucleocytoplasmic transport and on lamin rearrangement. Throughout, we related our modeling predictions to our experiments measuring the effects of nuclear deformation in U2OS cells on nanopillar substrates to reliably characterize cell response to nanopillar substrates with varied height, radius, and spacing (Figure 1A-B). Our approach incorporates mechanical deformation of the nucleus (Figure 1C-D) and reaction-transport of YAP/TAZ, NPCs, and lamin A/C (Figure 1E). We compare several modeling outputs directly against experimental measurements, including nuclear morphology, YAP nuclear-to-cytosolic ratio, lamin localization, and NE rupture likelihood. This framework generates quantitative predictions of higher nanotopography-mediated NE rupture rates in cells with lower lamin expression, which we confirm in subsequent experiments. Overall, our model and data together offer new insights into the complex interplay between nuclear mechanics and mechanotransduction signaling.

**Figure 1:**
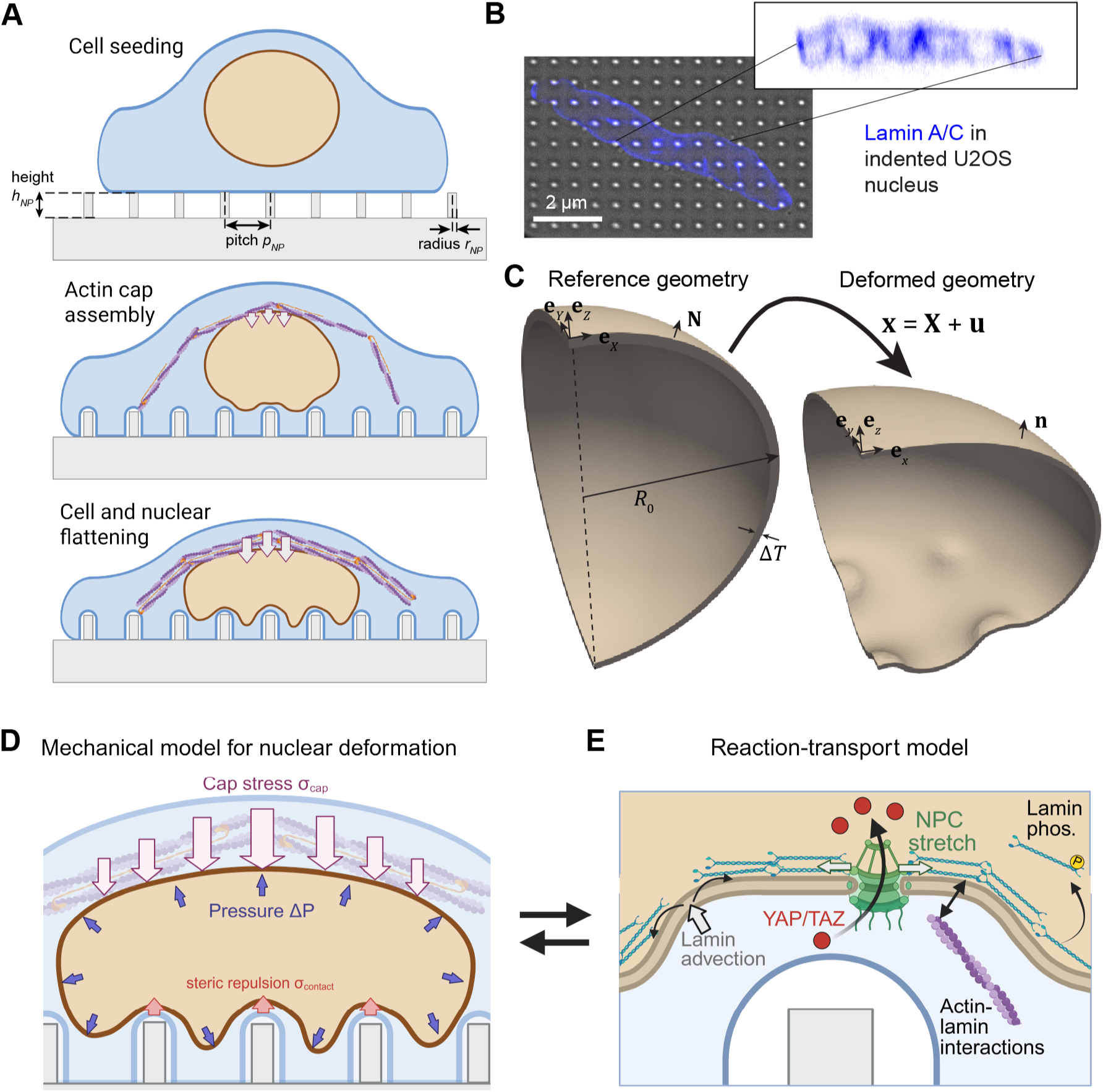
Modeling framework to study large nuclear deformations on nanopillar substrates. A) Schematic showing stages of cell adhesion and nuclear compression in cells on nanopillar substrates. Perinuclear actin cap assembly leads to cell and nuclear flattening over time. Variables describing nanopillar height (*h*_NP_), radius (*r*_NP_), and center-to-center spacing (pitch, *p*_NP_) are defined in the upper schematic. B) Deformed U2OS nucleus with stained lamin A/C. Inset shows the orthogonal view of lamin A/C intensity. C) Reference mesh and deformed geometry of NE shell utilized in our mechanical model. The reference mesh is a spherical shell with outer radius *R*_0_ and thickness Δ*T*, deformed according to the displacement field **u**. Unit vectors denote coordinate directions ([*X, Y, Z*] in reference geometry, [*x, y, z*] in deformed geometry) and **N** and **n** denote the normal vectors associated with the reference and the deformed geometry, respectively. D) Hyperelastic mechanical model of the NE with applied stresses due to steric repulsion at nanopillar contact regions (*σ*contact), osmotic pressure (Δ*P*), and assembly of the actin cap (*σ*cap). E) Mixed-dimensional reaction-transport model for lamins, NPCs, and YAP/TAZ. This model includes the effects of dilution and aggregation due to advection of species in the NE, as well as diffusion of all volumetric species. Arrows between panels D and E indicate bidirectional coupling via transport and NPC dilation (left-to-right) and via lamin-dependent changes in nuclear stiffness (right-to-left). Schematics in A, D, and E created with BioRender.com.

## Results

### Model development

Our modeling approach relied on two modules, including a mechanical model describing deformation of the NE (Figure 1C-D) and a mixed-dimensional reaction-transport model describing evolution of NPCs, lamin A/C, and YAP/TAZ over time (Figure 1E). These two modules were bidirectionally coupled. Mechanical deformations drive changes in nuclear geometry, advective transport of signaling species, and mechanically-induced dilation of NPCs. In turn, lamin A/C density dictates the local stiffness of the nuclear lamina. Below, we briefly summarize both modules. The governing equations and details on numerical implementation are provided in Supporting Information.

Our mechanical model treated the NE and the lamina together as a three-dimensional composite including two lipid bilayers and the underlying lamin meshwork (Figure 1D, Fig S1). The NE-lamina composite was approximated as an incompressible rubber-like hyperelastic material in line with previous quantitative models and experiments [39, 35, 36], allowing us to capture nonlinearities associated with large strains of elastic bodies [47, 48]. Our choice of a hyperelastic shell model rather than a Helfrich-type model or a model dominated by the bulk mechanics of the nucleoplasm (fluid droplet or elastic continuum) reflects the assumption that the mechanics of nuclear deformation are largely determined by the elasticity of the lamina [24, 26]. We assume that the mechanics of the lamina are dominated by lamin A/C rather than B-type lamins [26], an assumption utilized in our biochemical model below. While the composite is incompressible, our model allowed the volume of the enclosed nucleoplasm to decrease during compression, corresponding to water loss through the semipermeable nuclear membrane [8, 35]. We accounted for resulting changes to the osmotic pressure gradient across the NE due to volume changes, in line with previous studies [49] (Fig S2). The areas of the inner and outer surfaces of the shell also changed during nuclear deformations; biophysically, this corresponds to folding and unfolding of the NE [35, 42, 14, 50] and lateral remodeling of the lamin meshwork [18, 51]. In each simulation, we subjected an initially spherical NE shell to a resting osmotic pressure gradient [35] (Δ*P*) and then introduced a downward force due to the perinuclear actin cap (*σ*_cap_) (Figure 1D) [52, 53]. The region of the composite directly above nanopillar regions was subjected to a repulsive stress (*σ*_contact_) to capture the steric interactions with the actin layer between the NE and the plasma membrane adherent to the substrate [30] (Figure 1D). The magnitude of stress from the perinuclear actin cap (*σ*_cap_) was varied to capture multiple states during actin cap assembly (Figure 1A) and, ultimately, to reproduce previous measurements of large-scale nuclear deformations observed in cells on nanopillar substrates (Figure 1B) [30, 19]. In all cases, we assumed that the nucleus was at mechanical equilibrium but the actin cap stress and osmotic pressure evolved over time.

To capture the effects of nuclear deformation on nucleocytoplasmic transport and lamin A/C localization, we coupled the mechanical model to a biochemical reaction-transport model implemented in our recently developed software package, Spatial Modeling Algorithms for Reactions and Transport (SMART) [44]. This portion of the model was informed by our previous work [44], but only considers the nuclear surface and nucleoplasm in spatial detail, as opposed to the whole cell, fixed geometry approach previously used. We demonstrate the congruence between our updated model here and simulations with our previous model in Fig S7. In brief, this signaling model assumed curvature-dependent actin assembly around inwardly curved regions of the cell membrane in line with our previous simulations [20] and as characterized across several experimental studies [33, 32]. For the sake of simplicity, we did not explicitly consider actomyosin dynamics contributing to perinuclear cap assembly in this reaction-transport module, effectively treating actin within the perinuclear cap as a separate pool. Cytoplasmic actin engages with the nuclear lamina via the linker of cytoskeleton and nucleoskeleton (LINC) complex, thus transmitting force to the NE [54, 55, 56]. This interaction, together with the overall deformation of the NE, is assumed to facilitate dilation of nuclear pore complexes (NPCs) [15, 57], as described in further detail in a later section (Figure 1E). We tracked the transport of NPCs and lamins via advection during nuclear deformations, assuming that each species was nondiffusive in line with experimental measurements [58, 59]. We also considered the phosphorylation-regulated cycling of lamins between the nucleoplasm and NE (Figure 1E) [60, 27], wherein lamins were either retained at the NE in an actin-dependent manner or were phosphorylated and solubilized in the nucleoplasm. Given these assumptions for lamin A/C and NPC dynamics, we predicted YAP/TAZ transport in and out of the nucleus, assuming that YAP/TAZ within the cytosol remained well-mixed over time.

We solved both the mechanical and the biochemical reaction-transport equations using the finite element method, as detailed in Supporting Information. Below, we first consider the results from solving the mechanical model in isolation, and then describe the coupled mechanochemical model of nuclear deformation and reaction-transport. In both cases, we directly compare our model predictions to experimental measurements of U2OS cells on nanopillar substrates.

### Simulated nuclear envelope deformation by nanopillars agrees with experimental measurements

We first examined the equilibrium conformation of the NE for increasing applied stresses on the upper region of the nucleus due to the assembly of the perinuclear actin cap. To probe the effects of microenvironment nanotopography on NE deformation, we simulated nuclear compression on different nanopillar substrates with nanopillar radius (*r*_NP_) 200 nm and height (*h*_NP_) 2.5 µm. We note that in experiments, the nanopillar radius used for most experiments was measured between 0.3285 and 0.3745 µm (Fig S4). Our choice of nanopillar radius of 0.2 µm was used to approximate the experimental dimensions tested throughout. While we did not exactly match the measured nanopillar radius, we note that the material conclusions of our mechanical simulations are similar when the nanopillar radius is 350 nm (Fig S3). Furthermore, in our model treatment, the NE bends quite gradually around the peripheral region of nanopillars, as shown in Fig S1D-E. This is because we assume that the PM adherent to the nanopillar surface adds 50 nm to the effective nanopillar radius and the resulting boundary of the PM is not sharp, but is prescribed by a smooth function (Fig S1D, Eq S48). We examined the range of nuclear indentation and NE tension as a function of nanopillar spacing (pitch, *p*_NP_), including deformation on a flat substrate as the control condition.

**Figure 2:**
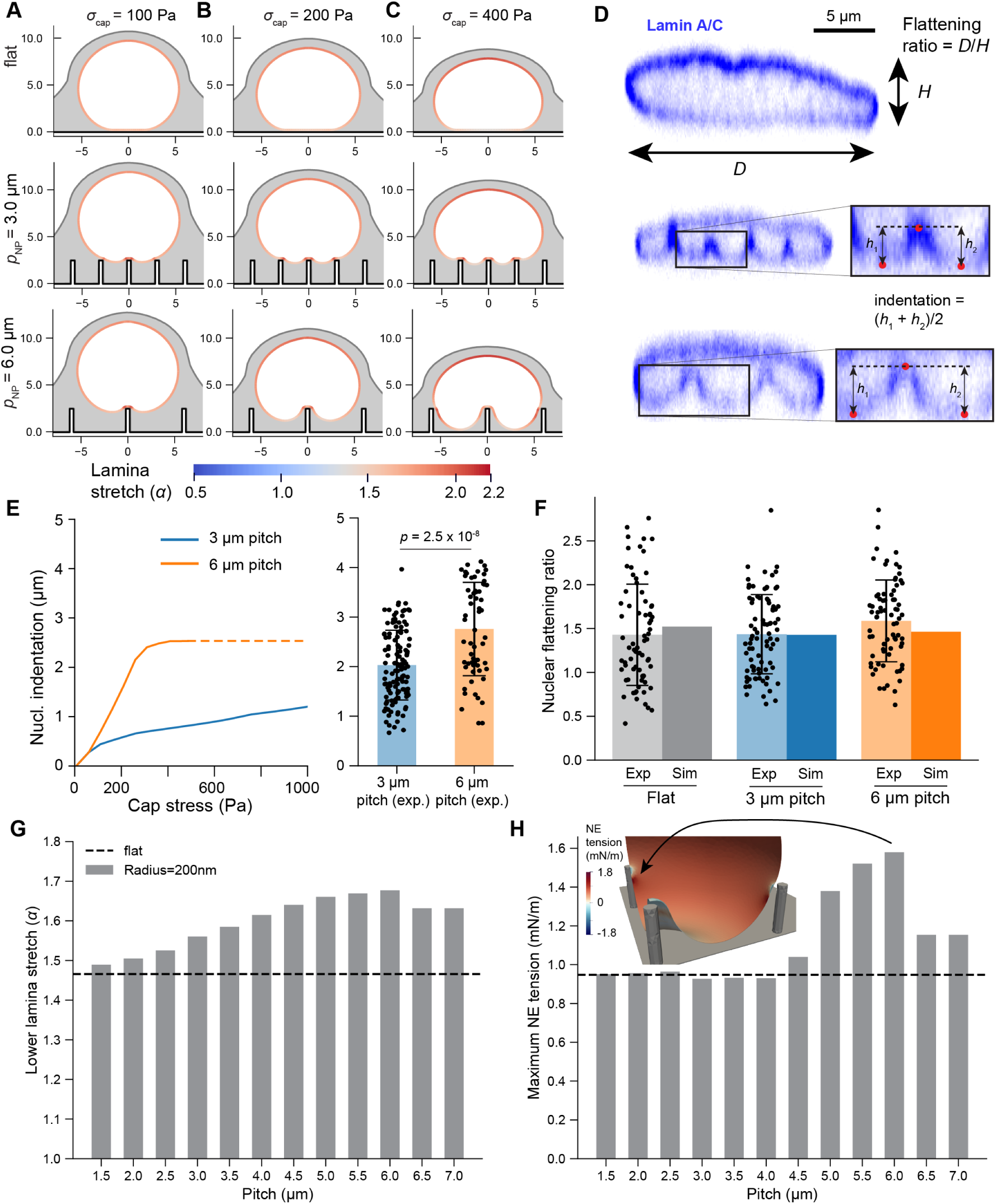
Simulated nuclear morphologies agree with experimental measurements. A-C) Cross sections of equilibrium conformations of deformed nuclei on a flat substrate or substrates with *r*_NP_ =200 nm, *h*_NP_ =2.5 µm, and *p*_NP_ = 3.0 µm or 6 µm for cap stress (*σ*cap) of 100 Pa (A), 200 Pa (B), or 400 Pa (C). The colormap indicates the magnitude of nuclear lamina stretch at the inner surface of the composite. D) Experimentally measured nuclear morphologies on flat, 3 µm, and 6 µm pitch substrates. Lamin A/C immunofluorescence images are shown for each case, with insets depicting the definition of NE indentation used in image analysis. E) Comparison of nuclear indentation in simulations versus experiments on 3 micron pitch (*N* = 24 cells, 124 indentations), and 6 micron pitch (*N* = 24 cells, 60 indentations) substrates, collected 8 hours after seeding. Simulation values in panel E correspond to *z*-displacement of the NE at the central nanopillar, and the dashed line begins where simulations no longer reliably converge for taller nanopillars. F) Comparison of nuclear flattening index in simulations versus experiments on flat (*N* = 72 cells), 3 micron pitch (*N* = 84 cells), and 6 micron pitch (*N* = 75 cells) substrates. Note that these indices were calculated from the DAPI channel in the YAP/Ku-80/DAPI imaging experiments. Simulation values are shown for a cap stress of 400 Pa. G) Average inner lamina stretch as a function of nanopillar pitch for 400 Pa cap stress. Average stretch was computed over the elements of the inner lamina surface with *z* values less than *z*center + 0.05, where *z*center is the *z* value associated with the point of the inner NE directly above the central nanopillar (Fig S1B). H) Maximum NE tension as a function of nanopillar pitch for 400 Pa cap stress. Inset shows the region of peak tension directly adjacent to the nanopillar for a 6 µm pitch.

We first assessed the effect of nanopillar spacing on compression and indentation of the NE (Figure 2,Movie 1). Nuclear indentation was minimal on low-pitch substrates; the nucleus rested on top of the pillar array even for high cap stresses, resulting in similar morphologies to a nucleus on a flat substrate (Figure 2A-C). Increasing the nanopillar pitch led to a monotonic increase in nuclear indentation, up to saturation at the height of the nanopillars themselves (Figure 2E). We also assessed nuclear compression as quantified by the flattening index, the nuclear width divided by the nuclear height. This quantity changed by less than 10% as a function of nanopillar pitch (Figure 2F), suggesting that global nuclear compression by the perinuclear actin cap is consistent regardless of the underlying nanotopography.

We then tested these predictions in experiments with U2OS cells on nanopillar substrates. U2OS cells were seeded on nanopillars with diameter 0.703 *±* 0.046 µm, height 2.98 *±* 0.16 µm, and pitches (center-to-center spacing) of either 3.09 *±* 0.05 µm or 6.05 *±* 0.13 µm. We fixed cells 8 hours after seeding and stained them for lamin A/C, Ku-80 and DAPI to assess nuclear morphology in intact cells. We imaged via confocal microscopy, observing that nuclear flattening on nanopillar substrates was associated with large indentations of the nuclear lamina (Figure 2D). In agreement with our model predictions and with previous work [30], nuclear indentation was higher on the higher-pitch substrate (Figure 2E). As in previous work [19], the magnitude of indentation was shown to increase over time on the lower pitch substrate (Fig S6). We note that the overall indentation predicted by simulations were generally on the lower end of those measured in experiments for 3 µm pitch substrates; this may be attributed to factors not currently considered in the model including viscosity of the NE-lamin composite (see Discussion).

We then conducted an additional set of experiments in which we assessed the correlation between bulk nuclear morphology and changes in the nuclear localization of YAP and Ku-80 (YAP discussed in the following section). For the purpose of these measurements, we used the Ku-80 channel to identify non-ruptured nuclei and the DAPI channel to quantify the flattening index. The resulting flattening indices in non-ruptured cells did not change significantly between substrates, and matched well with the computed values from our model for moderate cap stress (*σ*_cap_ = 400 Pa) (Figure 2F).

Having established that our model predicts nuclear shapes in good agreement with experiments, we considered the implications of these simulated deformations for changes in nuclear lamina stretch and NE tension (Figure 2G-H), as both are proposed predictors of accelerated nucleocytoplasmic transport [12, 13, 16] and susceptibility to NER [18, 61]. We first computed the stretch ratio *α*, defined as the local ratio of the deformed surface area to the reference surface area (see Eq S26). Increased indentation on moderate to high pitch substrates was accompanied by higher values of stretch adjacent to nanopillar contact regions (Fig S5A, Movie 1). Somewhat surprisingly, the magnitude of the stretch ratio averaged over the lower region of the NE increased up to some critical pitch value, beyond which this stretch declined (Figure 2G). This effect is a consequence of cells on high-pitch nanopillar substrates only contacting a single central nanopillar in our model, leading to a lower average stretch over the lower region. We then approximated the tension experienced by the NE by computing the average in-plane tension over the outer 50 nm layer of the NE-lamina composite, representing the thickness of both bilayers plus the perinuclear space [62] (Eqs S12-S14, Fig S5B). Overall, the maximum NE tension (highest value at any point within the geometry at a fixed cap stress of 400 Pa) followed a similar trend to stretch – nuclei exhibited the highest maximum tensions for pitches between 5 to 6 microns (Figure 2H). The peak tension was observed directly adjacent to nanopillar contacts formed on moderate-pitch nanopillar substrates (Figure 2H, inset, Movie 2). Altogether, our model predicts that NE tension and strain are both maximized at moderate-to-high pitch, a consequence of the tradeoff between contact with more nanopillars at lower pitches and the increasing local stretch facilitated by wider gaps between nanopillars.

**Figure 3:**
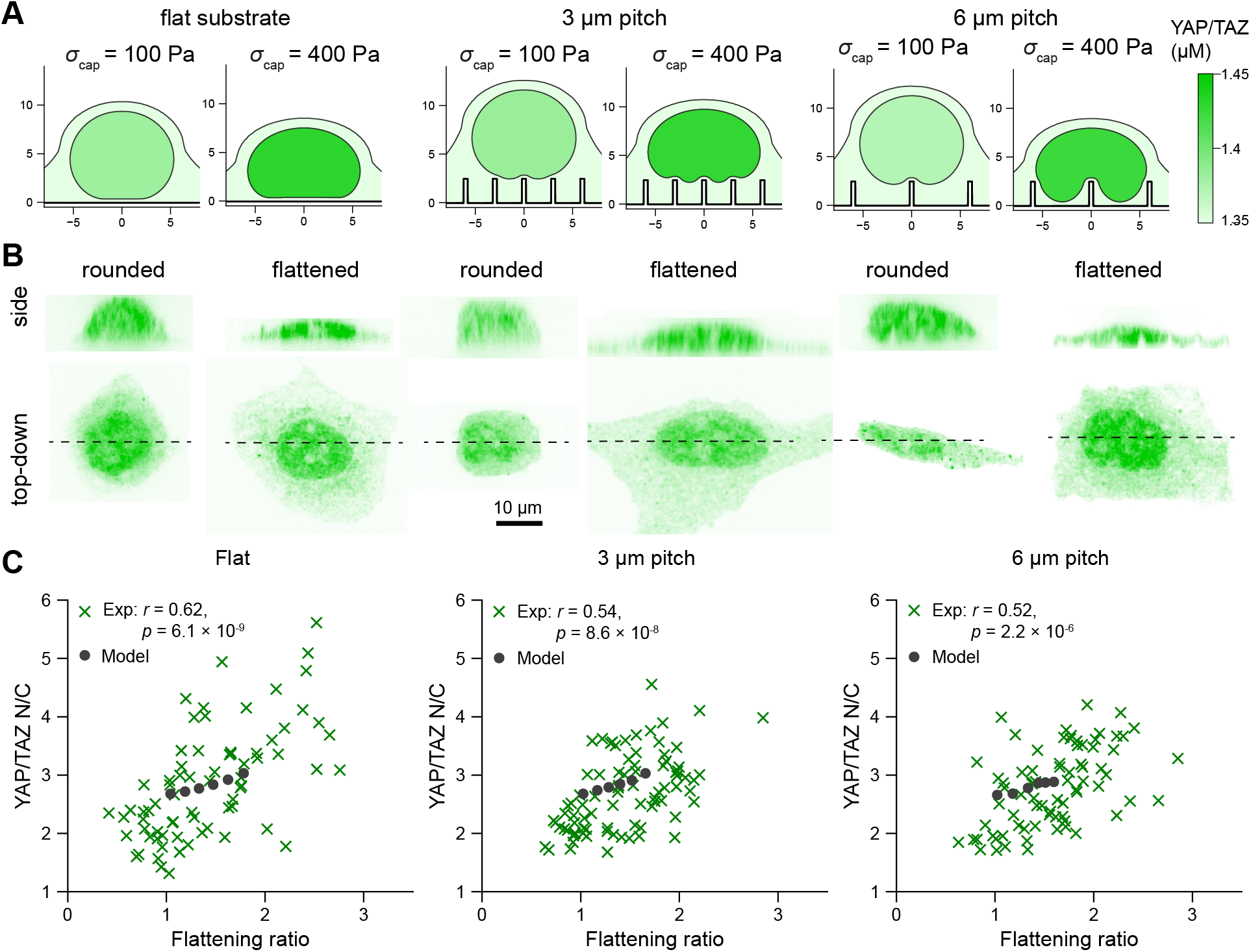
YAP/TAZ nuclear localization correlates with compression on flat or nanopillar substrates. A) Model predictions of YAP/TAZ localization in the nucleus and the cytoplasm at low cap stress (100 Pa) and high cap stress (400 Pa) on each substrate. Reported concentrations correspond to the average concentration in the respective domain at the final time point of the simulation (*t* = 10000 s). B) Representative images of YAP localization in experiments in cells on a flat substrate (*N* = 72 cells) or on 3 (*N* = 84 cells) or 6 µm pitch (*N* = 75 cells) substrates, with an example of a rounded nucleus and a flattened nucleus shown in each case. For each cell, the side-view (top) is shown above the maximum intensity projection of the top-down view (bottom). C) Comparison of experimentally measured correlations between YAP N/C and the nuclear flattening index and model predictions for nuclei experiencing different cap stresses (from left to right on each plot – 0, 100, 200, 300, 400, 500 Pa). Reported statistics denote Pearson’s correlation coefficient (*r*) and the *p*-value associated with the null hypothesis *r* = 0.

### Nuclear compression leads to increased YAP/TAZ nuclear translocation

Previous experimental work suggests that the rate of molecular import through NPCs is sensitive to the stretch of the NE [12, 13, 16]. Accordingly, regions of high tension and stretch in the NE identified above are likely sites of NPC dilation. To investigate this further, we coupled our mechanical model to a reaction-transport model of NPC dilation, YAP/TAZ transport, and lamin localization (Figure 1E). In these simulations, we assumed that the cap stress evolves over time:

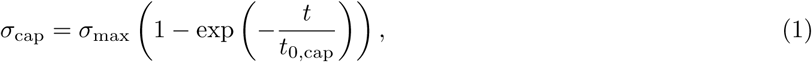

where *σ*_max_ sets the maximum cap stress after the cell has fully spread and *t*_0,cap_ sets the characteristic timescale of actin cap assembly. We assumed that NPC dilation was mediated by overall nuclear deformation and by coupling between the cytoskeleton and nucleoskeleton through the LINC complex, in line with previous experiments [12].

The total import rate of YAP/TAZ through NPCs is given by

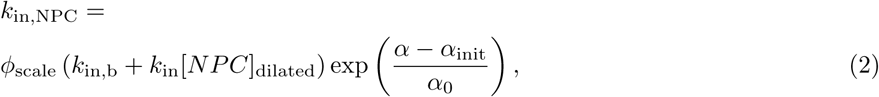

where [*NPC*]_dilated_ is the density of dilated NPCs due to stress transduction via LINC complexes (see Eqns S37-S38), *α*_init_ is the average NE stretch at *t* = 0 (after increasing the nuclear pressure), and *k*_in,b_ and *k*_in_ are the baseline and NPC-dilation-dependent import rates. We treated two parameters as unknowns – *α*_0_, which determines the NPC sensitivity to NE stretch, and *ϕ*_scale_, which is a cell-type dependent flux scaling factor. *ϕ*_scale_ was fixed to ensure that the YAP/TAZ N/C in nondeformed nuclei agreed with experiments. We assume that the export rates through NPCs remain similar regardless of mechanical cues, in line with previous measurements [13].

To inform the cell geometries and constrain free parameters associated with this model, we utilized the U2OS cell experiments described above, staining for YAP, Ku-80 (to monitor for NER), and DAPI. This allowed us to simultaneously quantify YAP nuclear-to-cytosolic ratio (N/C) and nuclear flattening for intact nuclei. Across all cases, we observed a clear correlation between nuclear flattening and YAP N/C (Figure 3B-C). In contrast to previous measurements [20, 32, 31], we found that the overall YAP N/C was not significantly higher on flat substrates. In Fig S7, we use our previous whole-cell model [44] to show that this discrepancy is accounted for by similar contact areas measured across all conditions in our experiments. In most previous studies, cells on flat substrates spread much more, contributing to the elevated levels of nuclear YAP.

We then conducted mechanochemically coupled simulations, matching the overall nuclear-to-cytoplasmic volumes measured in experiments and using F-actin and activated myosin densities informed by runs of our previous model (Supporting Information, Fig S7). We tested a range of different values for NPC sensitivity to NE stretch (*α*_0_), finding that most cases resulted in a positive correlation between nuclear flattening and YAP/TAZ N/C, with the overall slope of this correlation determined by *α*_0_ (Fig S8). *α*_0_ = 0.5 resulted in the best match to our experimental data (Fig S6). Examining the model predictions from this set of parameters, we found that nuclear flattening was accompanied by a transition of NPCs to the dilated state and global stretch of the NE, accelerating YAP nuclear entry (Fig S9, Movie 3). NPC dilation was mediated by aggregation of F-actin around nanopillars, leading to stretch of the NE via LINC complex engagement. This structural assembly resulted in distinct patterns of lamin localization around nanopillar indentations. Overall, the model and data together indicate that nuclear flattening controls the extent of YAP/TAZ nuclear localization across different substrates, leading to changes in YAP/TAZ N/C despite similar levels of stimulus from the substrate.

### Substrate nanotopography modulates lamin distribution and loading

In our biochemical model, the lamin content in the NE was tightly regulated by the rates of lamin phosphorylation and dephosphorylation and by advective transport. To examine the time-dependent effects of nanotopography on lamin reorganization and laminar stresses, we quantified the local concentration of lamin within the laminar meshwork and the force experienced per lamin dimer (Eq S44). Given the established role of lamin A/C in nuclear integrity [18, 60, 24], we used this force per lamin dimer as a predictor for NER. We note that even in the absence of applied force, the force experienced per lamin dimer can change over time due to the kinetics of lamin phosphorylation controlling its localization to the NE vs the nucleoplasm.

Our model predicted an initial stage of lamin depletion in regions of high strain, followed by lamin enrichment around nanopillars at later time points (Figure 4A-B, Movie 4). The short term depletion was due to advective effects, whereas the long term enrichment was mediated by local actin assembly near regions of nanopillar indentation (Fig S10). The duration of the depletion stage was sensitive to the prescribed actin density and lamin phosphorylation rate (Fig S10), suggesting possible mechanisms for the variability of lamin A/C response to indentation in previous experiments [51]. The initial depletion resulted in a peak in the local force per lamin dimer surrounding the central nanopillar at intermediate time points (*t <* 1000 s), suggesting a period in which NER may be most likely (Figure 4C-D, Movie 5). The force per lamin dimer was higher on 6 micron pitch substrates, in line with our purely mechanical simulations. Regardless of nanopillar spacing, we observed consistent enrichment of lamin A/C surrounding pillars at later time points (*t >* 1000 s, Figure 4E).

We then compared our predictions to measurements of lamin localization in our experiments. On both 3 and 6 micron pitch substrates, lamin A/C was highest in nanopillar indented regions (Figure 4F-G). Examining these profiles across many cells on each substrate, we found that this effect was consistent, although there was some variability in terms of the lamin density directly above the pillar versus at the periphery of the nanopillar indentations (Figure 4F). In the model, this effect was mediated by lower levels of assumed actin polymerization adjacent to the flat membrane directly above the nanopillar (Eq S90). The resulting average lamin intensity profile around pillars showed good agreement between simulations and experiments (Figure 4G).

**Figure 4:**
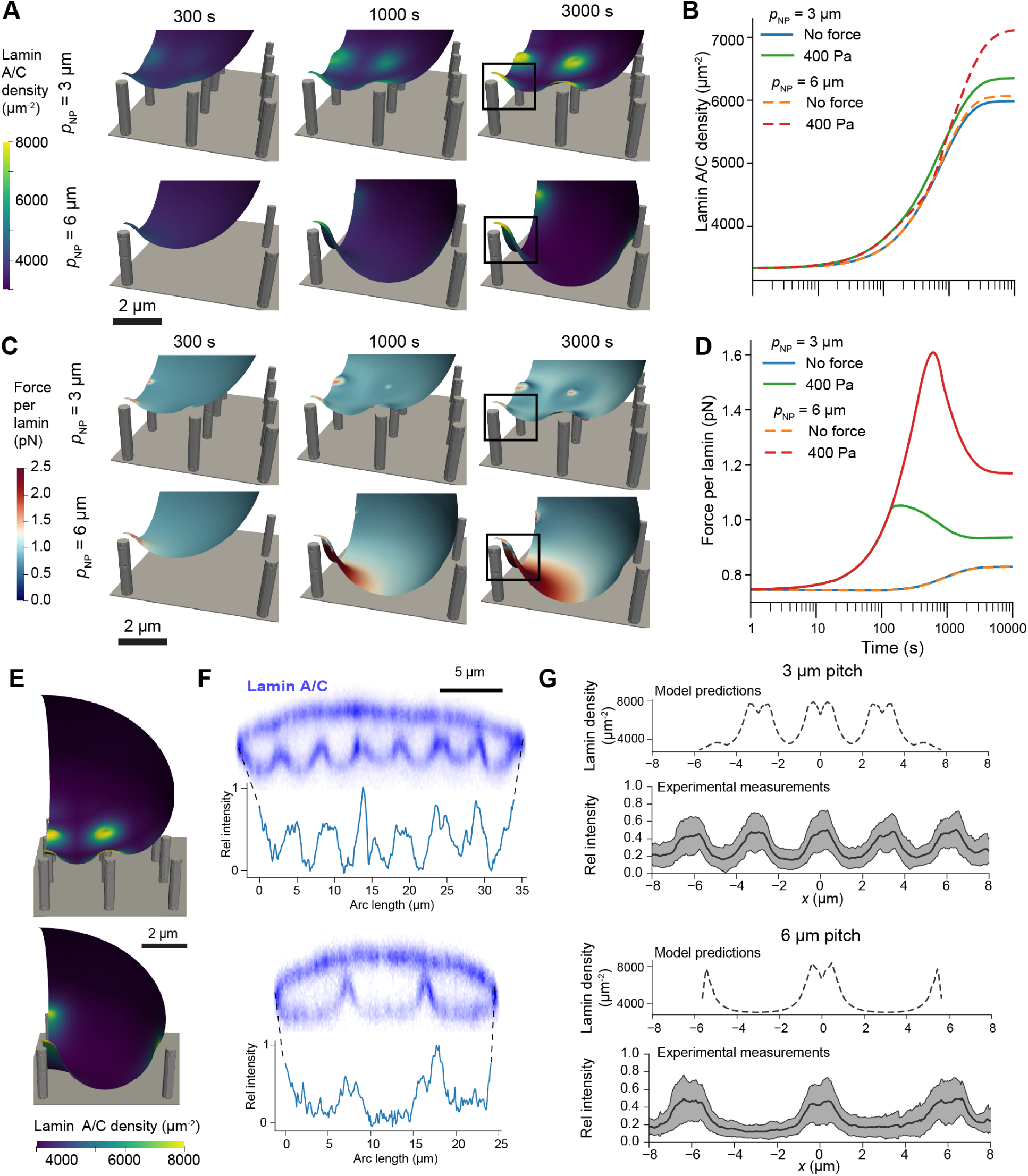
Lamin localization and load are modulated by NE indentation. A) Lamin A/C distribution within the nuclear lamina for an applied cap stress of 400 Pa. Lamin A/C density increases at later time points due to local actin assembly on 3 µm and 6 µm pitch nanopillar substrates. B) Average lamin A/C density within the identified region of interest in panel A shows that force application leads to increased lamin density surrounding nanopillars. C) The resulting force per lamin dimer within the nuclear lamina for the cases in panel A shows regions of high force at intermediate time points. D) Average force per lamin within the boxed regions in panel C shows an intermediate peak followed by force reduction due to lamin accumulation at later time points. E) Lamin distribution at final time points (*t* = 10000 s for 3 and 6 micron pitch nanopillar substrates. F) Representative images of lamin A/C distributions in cells on nanopillar substrates, with associated intensity traces along the nuclear boundary. G) Lamin density traces predicted by the model on each nanopillar substrate, versus those measured experimentally pooled across 2, 5, and 8 hr timepoints (*N* = 75 cells each substrate). Model traces show a consistent dip in lamin density directly above the nanopillar due to lower levels of actin assembly at this locally flat region. Experimental measurements were averaged after first scaling lamin intensities on a cell-by-cell basis to a range from 0 to 1. Data is shown as the mean *±* standard deviation.

### Model predictions agree with measurements of nuclear rupture in low-lamin cells

Having established that our model can accurately predict nuclear morphologies and lamin distributions, we asked whether our model could be used to predict the consequences of these morphology changes, including NER. Given that many previous studies have found low-lamin cells are more susceptible to NER [18, 27, 63], we tested the predictions of our model for cells with reduced lamin content. These model cells were assumed to be less rigid, as measured in previous studies [64, 27]. From these simulations, we used the computed force per lamin dimer to quantitatively predict the likelihood of NER, as fully described in Supporting Information. Treating individual ruptures as independent events governed by Bell’s law [65], we computed the total likelihood of a rupture occurring anywhere within the laminar network over the simulation (see Supporting Information, Eqs S90-S93). This introduced two new parameters associated with the baseline rupture rate and the molecular-scale displacement, which we constrained by comparing predictions with our previous measurements of NER (Supporting Information, Fig S11). Our simulations revealed consistently higher forces experienced per lamin dimer in lamin-deficient cells, resulting in a much higher rupture likelihood in these regions (Figure 5A, Movie 6). Considering a test case of larger radius (450 nm) nanopillars, we found that cells with normal (100%) lamin expression had an overall rupture likelihood of about 3.7% at some point during the simulated period, but cells expressing 70% of lamin approached 100% likelihood of rupture in our calculations (Figure 5A, Movie 6). We then considered a population of cells with a wider range of relative lamin expression (between 50% to 150%) and calculated the resulting cumulative rupture likelihoods. Sampling from these likelihoods at the 1 hr time point, we found that lamin expression was much lower on average in ruptured nuclei versus non-ruptured nuclei (Figure 5C).

To validate these modeling predictions, we cultured U2OS cells on engineered quartz nanopillar substrates (height 3.24 *±* 0.25 µm, diameter 0.90*±*0.016 µm, and pitch 3.49*±*0.06 µm). We reduced lamin A/C expression using small interfering RNA (siRNA) targeting the LMNA gene. The treated population showed heterogeneous lamin A/C expression, with many cells exhibiting a significant reduction in lamin content compared to controls (Figure 5A). To examine nuclear mechanical integrity across a wide range of lamin expression levels, we pooled single-cell data from both the siLMNA treated and non-silenced control populations (Figure 5B). Cells were immunostained for Ku-80 to detect NER via protein mislocalization from the nucleus to the cytoplasm. Using a data-driven Gaussian Mixture Model (GMM) on this combined population, we determined a Ku-80 cytoplasmic-to-nuclear ratio (C/N) of 0.16 as the threshold distinguishing ruptured from non-ruptured nuclei. This analysis revealed that low-lamin cells were much more abundant in the population with ruptured nuclei, suggesting that reduced lamin content increases rupture likelihood (Figure 5D-E) and providing strong agreement with our model predictions (Figure 5C).

## Discussion

Cell motility in physiological environments is strongly influenced by nuclear size and deformability. In contrast to many classic motility assays, in which cells freely migrate over flat 2D surfaces, migration through pericellular spaces in vivo requires cells to squeeze through small passages and adapt to nanoscale curvature [66, 67, 68]. In addition to the well-appreciated role of PM curvature in modulating protein aggregation and cellular signaling [66, 69], recent studies indicate that changes in nuclear curvature influence biochemical events at the NE [18, 70]. For instance, rapid induction of Gaussian curvature leads to dilution of lamin A/C and lamin B within the NE, increasing the likelihood of rupture [18, 51]. Independent of curvature, nuclear compression leads to nuclear volume loss [35, 16, 71] and increased likelihood of rupture [17]. Our study integrates these recent discoveries within a single modeling framework, elucidating the effects of compression and imposed curvature on stresses experienced by the nucleus and on remodeling of the nuclear lamina.

Our model of the NE relies on a hyperelastic formulation that supports large deformations, as utilized in several previous models [8, 35, 36, 39]. This approach incorporates more detailed mechanics of the nuclear surface than other recent models of the nucleus as a droplet with surface tension [14]. Our mechanical model alone does not explicitly include viscous terms, but we consider time-dependent changes in NE stiffness due to lamin reaction and transport, which effectively captures some degree of viscous remodeling. A major innovation in our approach lies in the coupling between nuclear mechanics and biochemical reaction-transport. Using our recently developed software framework [44] and state-of-the-art finite element analysis in FEniCS [72], we were able to recapitulate several key features of nuclear mechanotransduction in silico.

Our simulations make clear predictions about changes in YAP/TAZ nucleocytoplasmic transport, a well-established player in mechanotransduction signaling [73, 74]. In particular, we show that nuclear compression by the perinuclear actin cap leads to global stretch of the nuclear envelope, potentially facilitating dilation of NPCs [15]. We also assume that NPC dilation is in part facilitated by interactions between the nucleoskeleton and cytoskeleton via the LINC complex, as supported by studies that suggest disruption of the LINC complex leads to slower NPC-mediated import [16, 13]. The overall relationship between NE deformation and NPC opening likely involves a complex interplay between NE curvature, NE stretch, and nucleoskeleton-cytoskeleton interactions. The predicted effects of nanotopography on nucleocytoplasmic transport in this work may also extend to other known mechanosensitive transcription regulatory factors such as TWIST1 [75, 13].

**Figure 5:**
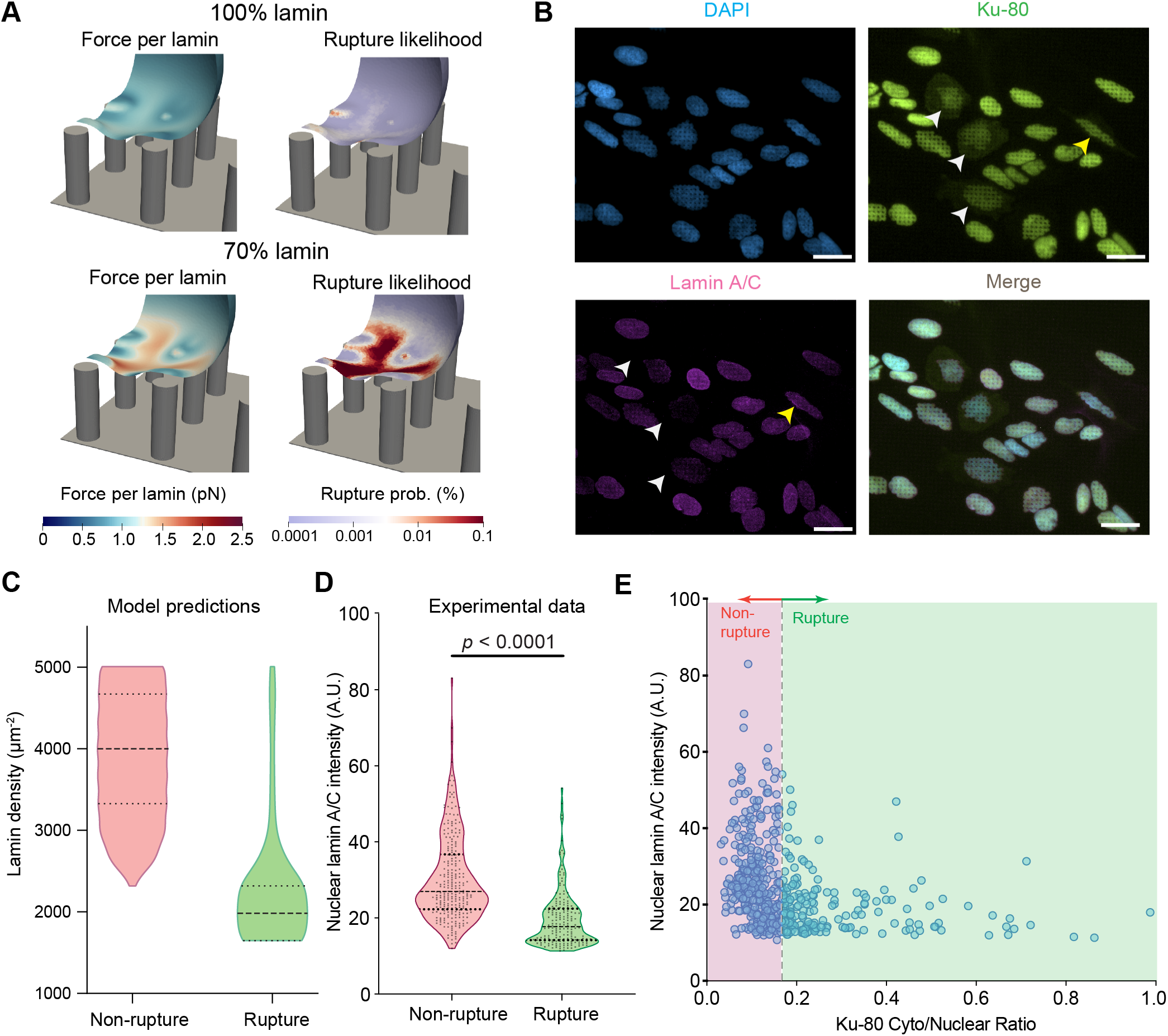
Low-lamin cells are more likely to exhibit NER on nanopillar substrates. A) Model predictions of force per lamin dimer and cumulative rupture likelihood in nuclei with 100% lamin expression versus 70% lamin expression on substrates with *r*_NP_ = 0.45 µm, *p*_NP_ = 3.5 µm, *h*_NP_ = 3.25 µm. B) Representative immunofluorescence images of U2OS cells treated with siLMNA on nanopillars. Cells were stained for DAPI (blue), Ku-80 (green), and lamin A/C (magenta). White arrows indicate cells with low lamin A/C expression that show NER evidenced by nucleus-to-cytoplasm mislocalization of Ku-80. The yellow arrow indicates a cell with normal lamin A/C expression that shows NER, consistent with stochastic rupture events on high-curvature nanotopography. Scale bar: 20 µm. C) Lamin densities in cells with ruptured nuclei versus non-ruptured nuclei. Simulations were run with 50%, 60%, 70%, 80%, 90%, 100%, 110%, 120%, 130%, 140%, and 150% lamin. D) Quantification of lamin A/C intensity for ruptured versus non-ruptured subpopulations classified in (B). **** denotes p *≤* 0.0001 according to a Student’s *t*-test and dashed horizontal lines in panel C indicate medians and upper and lower quartiles. E) Scatter plot characterizing the relationship between nuclear lamin A/C intensity and Ku-80 cytoplasmic-to-nuclear ratio (C/N) across the heterogeneous cell population. The vertical dashed line indicates the Ku-80 ratio threshold used to classify ruptured versus non-ruptured cells.

Our current model considers many key features of nuclear mechanics established in the literature; for instance, lamin redistribution was assumed to change the local NE stiffness [25] and also determined the force experienced by the NE, dictating the likelihood of NER [18, 14, 27]. However, many other molecular constituents not explicitly considered here also play important roles in nuclear mechanics. Although the LINC complex is implicitly represented by interactions between actin and lamin in our model (Figure 1E), we do not evaluate the direct contribution of LINC-mediated connections to the NE tension. This may help explain why the tension approximated by our model (Figure 2, up to 1 mN/m) is higher than previously estimated [76, 77]. Furthermore, our current estimates of tension neglect the detailed wrinkling of the NE observed in some experiments and models [35, 14, 42]. The surface area of the outer NE in our model reflects the apparent surface area, whereas the total membrane surface area is likely much larger, suggesting that our calculation of the tension may be an overestimate. Furthermore, the role for B-type lamins should be explored in more detail in future models; while we assume the laminar shell mechanics is dominated by lamin A/C here [26], B-type lamins can also contribute to nuclear deformability, particularly in other cell types [78]. Finally, our model does not explicitly consider the role of chromatin in nuclear mechanics. Although previous theoretical work has found that chromatin confinement stresses are negligible compared to the pressure gradient across the NE [49], changes in chromatin condensation [79] and interactions between chromatin and lamins [80] may still play a modulatory role in nuclear deformations. Furthermore, motors such as BRG1 that facilitate chromatin reorganization have recently been shown to strongly alter nuclear stiffness [81, 82], but further data is required to constrain models that include such proteins in descriptions of nuclear mechanics. Our model predicts that intermediate-to-high nanopillar pitch (5-6 µm) maximizes NE stress and stretch. These mechanically stressed states are expected to increase transient nuclear permeability through stretch-mediated NPC opening and, in some cases, NE rupture. This provides a geometry-guided design parameter for nuclear delivery and can facilitate nuclear access to cargoes that are otherwise transport-limited, such as large protein complexes (CRISPR RNPs), donor DNA templates, and transcriptional modulators (e.g., TEAD inhibitors) [83]. More broadly, these predictions reinforce the importance of nanotopography in shaping cellular responses in fibrous matrices where cells experience curvature-driven stress focusing at ECM contact points.

Our work has natural implications for the class of diseases known as laminopathies, which are associated with mutations in lamin A/C [28, 84]. Most laminopathies are associated with changes in nuclear morphology [85, 84, 26], suggesting changes in mechanical properties of the nuclear lamina. This may be attributed to changes in protein structure or in lamin phosphorylation state, leading to altered abundance of lamins in the NE [27, 86]. As shown by our simulations with low-lamin cells and experiments with lamin-depleted cells, reduced lamin content can render cells more susceptible to NER. In fact, several studies have measured increased prevalence of NER in lamin mutant cells [87, 88, 89]. Outside of the context of disease, lamin expression levels vary considerably across different tissues [27]. Our modeling framework could therefore be used to help elucidate changes in nuclear mechanics and mechanotransduction observed in cell types across different organs and tissues in the human body.

## Materials and Methods

### Numerical Methods

The coupled system was solved using the finite element method via FEniCS [72]. For this purpose, we leveraged a new version of our software, Spatial Modeling Algorithms for Reactions and Transport (SMART) [90], equipped to include advective transport. The hyperelastic problem was solved using the FEniCS Python API, allowing us to easily pass solutions to and from SMART, which also uses FEniCS. Our code is readily available on Github [91]. The full set of equations and numerical methods are in Supporting Information.

### Experimental Methods

Experimental methods can be found in Supporting Information.

## Use of AI

Google Gemini was used for literature searches; any papers identified from these searches were carefully reviewed before including as a reference. Claude (Sonnet 5) was used to conduct a mathematical consistency check across all equations in this work.

## Data, Materials, and Software Availability

All code used for the simulations shown in this study is available via the smart-mechanotransduction repository [91] (https://doi.org/10.5281/zenodo.18690945), which depends on a new release of our SMART package [90]. All code used to analyze experimental results and the compiled data are available via the Zenodo repository at https://doi.org/10.5281/zenodo.22905724 [92].

## Acknowledgments

The authors would like to thank Alessandro Contri for his helpful feedback on this manuscript, as well as Dr. Jørgen Dokken and Dr. Henrik Finsberg at Simula Research Laboratory for providing input on solving hyperelastic equations with contact boundary conditions in FEniCS. Simulation results presented in this paper benefited from the Triton Shared Compute Cluster at the San Diego Supercomputer Center [93]. This work was supported by the National Science Foundation under National Institutes of Health, grant number 1K99GM162992-01A1 (to E.A.F.), grant EEC-2127509 to the American Society for Engineering Education (to E.A.F.) and under MODULUS Grant MCB 2327243 (to P.R.). This work was performed in part at the San Diego Nanotechnology Infrastructure (SDNI) of UCSD, a member of the National Nanotechnology Coordinated Infrastructure, which was supported by the National Science Foundation (Grant ECCS-2025752). The authors would also like to thank Peng Guo and the Nikon Imaging Center at UCSD for their support with microscopy experiments. A.S. received funding from the Powell-Focht Fellowship from the Charles Lee Powell Foundation. This work was additionally supported by the Air Force Office of Scientific Research YIP award (AFOSR FA9550-23-1-0090) to Z.J. and by the National Institutes of Health (NIH) Institutional Training Grant T32 EB 009380 to H.N.N. P.R. is a consultant for Simula Research Laboratories in Oslo, Norway and receives income. The terms of this arrangement have been reviewed and approved by the University of California, San Diego in accordance with its conflict-of-interest policies. Z.J. and E.S. are inventors on a patent application related to this work (Patent Pending No. 009062-8542. WO00).

## Author Contributions

E.A.F. designed research, performed research, analyzed data, wrote the paper. A.S. performed research, analyzed data, wrote the paper. E.S. performed research and analyzed data. H.N.N. analyzed data and wrote the paper. D.P.M. contributed new reagents/analytic tools. Z.J. and P.R. designed research, supervised the project, and wrote the paper.

## References

[1] Yang Yang, Luis A Valencia, Chih-Hao Lu, Melissa L Nakamoto, Ching-Ting Tsai, Chun Liu, Huaxiao Yang, Wei Zhang, Zeinab Jahed, Wan-Ru Lee, Francesca Santoro, Jen Liou, Joseph C Wu, and Bianxiao Cui. Plasma membrane curvature regulates the formation of contacts with the endoplasmic reticulum. Nat. Cell Biol., 26(11):1878–1891, 17 November 2024.

[2] Yan Liu, Bingcheng Yi, Liangliang Yang, Yanyan Yang, Tianxiang Li, Xiaolu Li, Jae Youl Cho, Dengshen Zhang, Qihui Zhou, and Tao Yu. Anisotropic micro/nanotopography regulating mitochondrial dynamics in cardiomyocytes. Research (Wash. D.C.), 2025(0891):0891, 16 September 2025.

[3] A J Lomakin, C J Cattin, D Cuvelier, Z Alraies, M Molina, G P F Nader, N Srivastava, P J Sáez, J M Garcia-Arcos, I Y Zhitnyak, A Bhargava, M K Driscoll, E S Welf, R Fiolka, R J Petrie, N S De Silva, J M González-Granado, N Manel, A M Lennon-Duménil, D J Müller, and M Piel. The nucleus acts as a ruler tailoring cell responses to spatial constraints. Science, 370(6514):eaba2894, 16 October 2020.

[4] Patricia M Davidson, Celine Denais, Maya C Bakshi, and Jan Lammerding. Nuclear deformability constitutes a rate-limiting step during cell migration in 3-D environments. Cell. Mol. Bioeng., 7(3):293–306, 1 September 2014.

[5] Yuntao Xia, Charlotte R Pfeifer, and Dennis E Discher. Nuclear mechanics during and after constricted migration. Acta Mech. Sin., 35(2):299–308, 22 April 2019.

[6] Hong-Pyo Lee, Farid Alisafaei, Kolade Adebawale, Julie Chang, Vivek B Shenoy, and Ovijit Chaudhuri. The nuclear piston activates mechanosensitive ion channels to generate cell migration paths in confining microen-vironments. Sci. Adv., 7(2):eabd4058, January 2021.

[7] Michelle A Baird, Cayla E Jewett, Daniela A Malide, Lisa Kratz, Alexander X Cartagena-Rivera, Robert S Fischer, and Clare M Waterman. Lamin B receptor upregulation in metastatic melanoma causes nuclear envelope fragility in confined migration during cancer invasion. Proc. Natl. Acad. Sci. U. S. A., 123(8):e2513031123, 24 February 2026.

[8] Xuan Cao, Emad Moeendarbary, Philipp Isermann, Patricia M Davidson, Xiao Wang, Michelle B Chen, Anya K Burkart, Jan Lammerding, Roger D Kamm, and Vivek B Shenoy. A chemomechanical model for nuclear morphology and stresses during cell transendothelial migration. Biophys. J., 111(7):1541–1552, 4 October 2016.

[9] Melanie Salvermoser, Daniela Begandt, Ronen Alon, and Barbara Walzog. Nuclear deformation during neutrophil migration at sites of inflammation. Front. Immunol., 9:2680, 16 November 2018.

[10] Ning Wang, Shu Chien, and Martin A Schwartz. Mechanomedicine: Present state and future promise. Proc. Natl. Acad. Sci. U. S. A., 122(46):e2509566122, 18 November 2025.

[11] Yohalie Kalukula, Giuseppe Ciccone, Danahe Mohammed, Anthony Procès, Marie Versaevel, Amandine Deridoux, Lucie Ergot, Zoé Barbier, Maxime Mansy, Roxane Aucouturier, Rémi Tranzer, Mathieu Surin, Sylvain Gabriele, and Marine Luciano. Unlocking the therapeutic potential of cellular mechanobiology. Sci. Adv., 11(44):eaea6817, 31 October 2025.

[12] Alberto Elosegui-Artola, Ion Andreu, Amy E M Beedle, Ainhoa Lezamiz, Marina Uroz, Anita J Kosmalska, Roger Oria, Jenny Z Kechagia, Palma Rico-Lastres, Anabel-Lise Le Roux, Catherine M Shanahan, Xavier Trepat, Daniel Navajas, Sergi Garcia-Manyes, and Pere Roca-Cusachs. Force triggers YAP nuclear entry by regulating transport across nuclear pores. Cell, 171(6):1397–1410.e14, 30 November 2017.

[13] Ion Andreu, Ignasi Granero-Moya, Nimesh R Chahare, Kessem Clein, Marc Molina-Jordán, Amy E M Beedle, Alberto Elosegui-Artola, Juan F Abenza, Leone Rossetti, Xavier Trepat, Barak Raveh, and Pere Roca-Cusachs. Mechanical force application to the nucleus regulates nucleocytoplasmic transport. Nat. Cell Biol., 24(6):896– 905, 9 June 2022.

[14] Ting-Ching Wang, Samere Abolghasemzade, Brendan P McKee, Ishita Singh, Kavya Pendyala, Mohammad Mohajeri, Hailee Patel, Aakansha Shaji, Anna L Kersey, Kajol Harsh, Simran Kaur, Christina R Dollahon, Sasanka Chukkapalli, Pushkar P Lele, Daniel E Conway, Akhilesh K Gaharwar, Richard B Dickinson, and Tanmay P Lele. Matrix stiffness drives drop like nuclear deformation and lamin a/C tension-dependent YAP nuclear localization. Nat. Commun., 15(1):10151, 22 November 2024.

[15] Christian E Zimmerli, Matteo Allegretti, Vasileios Rantos, Sara K Goetz, Agnieszka Obarska-Kosinska, Ievgeniia Zagoriy, Aliaksandr Halavatyi, Gerhard Hummer, Julia Mahamid, Jan Kosinski, and Martin Beck. Nuclear pores dilate and constrict in cellulo. Science, 374(6573):eabd9776, 10 December 2021.

[16] Newsha Koushki, Ajinkya Ghagre, Luv Kishore Srivastava, Clayton Molter, and Allen J Ehrlicher. Nuclear compression regulates YAP spatiotemporal fluctuations in living cells. Proc. Natl. Acad. Sci. U. S. A., 120(28):e2301285120, 11 July 2023.

[17] Celine M Denais, Rachel M Gilbert, Philipp Isermann, Alexandra L McGregor, Mariska te Lindert, Bettina Weigelin, Patricia M Davidson, Peter Friedl, Katarina Wolf, and Jan Lammerding. Nuclear envelope rupture and repair during cancer cell migration. Science, 352(6283):353–358, 15 April 2016.

[18] Charlotte R Pfeifer, Michael P Tobin, Sangkyun Cho, Manasvita Vashisth, Lawrence J Dooling, Lizeth Lopez Vazquez, Emma G Ricci-De Lucca, Keiann T Simon, and Dennis E Discher. Gaussian curvature dilutes the nuclear lamina, favoring nuclear rupture, especially at high strain rate. Nucleus, 13(1):129–143, 16 December 2022.

[19] Einollah Sarikhani, Vrund Patel, Zhi Li, Dhivya Pushpa Meganathan, Keivan Rahmani, Leah Sadr, Ryan Hosseini, Diether Visda, Shivani Shukla, Hamed Naghsh-Nilchi, Adarsh Balaji, Gillian McMahon, Shaoming Chen, Johannes Schöneberg, Colleen A McHugh, Lingyan Shi, and Zeinab Jahed. Engineered nanotopographies induce transient openings in the nuclear membrane. Adv. Funct. Mater., 35(6):2410035, 5 February 2025.

[20] Emmet A Francis, Einollah Sarikhani, Vrund Patel, Dhivya Pushpa Meganathan, Leah Sadr, Anum Tahir, Zeinab Jahed, and Padmini Rangamani. Nanoscale curvature regulates YAP/TAZ nuclear localization through nuclear deformation and rupture. Adv. Sci. (Weinh.), page e2415029, 3 June 2025.

[21] Zeinab Jahed, Mohammad Soheilypour, Mohaddeseh Peyro, and Mohammad R K Mofrad. The LINC and NPC relationship - it’s complicated! *J*. Cell Sci., 129(17):3219–3229, 1 September 2016.

[22] Alexandre Méjat and Tom Misteli. LINC complexes in health and disease. Nucleus, 1(1):40–52, January 2010.

[23] Abigail Buchwalter. Intermediate, but not average: The unusual lives of the nuclear lamin proteins. Curr. Opin. Cell Biol., 84(102220):102220, 22 October 2023.

[24] Kris Noel Dahl, Samuel M Kahn, Katherine L Wilson, and Dennis E Discher. The nuclear envelope lamina network has elasticity and a compressibility limit suggestive of a molecular shock absorber. J. Cell Sci., 117(Pt 20):4779–4786, 15 September 2004.

[25] Luv Kishore Srivastava, Zhaoping Ju, Ajinkya Ghagre, and Allen J Ehrlicher. Spatial distribution of lamin a/C determines nuclear stiffness and stress-mediated deformation. J. Cell Sci., 134(10):jcs248559, 15 May 2021.

[26] Jan Lammerding, Loren G Fong, Julie Y Ji, Karen Reue, Colin L Stewart, Stephen G Young, and Richard T Lee. Lamins a and C but not lamin B1 regulate nuclear mechanics. J. Biol. Chem., 281(35):25768–25780, 1 September 2006.

[27] Joe Swift, Irena L Ivanovska, Amnon Buxboim, Takamasa Harada, P C Dave P Dingal, Joel Pinter, J David Pajerowski, Kyle R Spinler, Jae-Won Shin, Manorama Tewari, Florian Rehfeldt, David W Speicher, and Dennis E Discher. Nuclear lamin-A scales with tissue stiffness and enhances matrix-directed differentiation. Science, 341(6149):1240104, 30 August 2013.

[28] Xianrong Wong and Colin L Stewart. The laminopathies and the insights they provide into the structural and functional organization of the nucleus. Annu. Rev. Genomics Hum. Genet., 21(1):263–288, 31 August 2020.

[29] Howard J Worman. Nuclear lamins and laminopathies. J. Pathol., 226(2):316–325, January 2012.

[30] Lindsey Hanson, Wenting Zhao, Hsin-Ya Lou, Ziliang Carter Lin, Seok Woo Lee, Praveen Chowdary, Yi Cui, and Bianxiao Cui. Vertical nanopillars for in situ probing of nuclear mechanics in adherent cells. Nat. Nanotechnol., 10(6):554–562, June 2015.

[31] Xiao Li, Lasse H Klausen, Wei Zhang, Zeinab Jahed, Ching-Ting Tsai, Thomas L Li, and Bianxiao Cui. Nanoscale surface topography reduces focal adhesions and cell stiffness by enhancing integrin endocytosis. Nano Lett., 21(19):8518–8526, 13 October 2021.

[32] Catherine S Hansel, Spencer W Crowder, Samuel Cooper, Sahana Gopal, Maria João Pardelha da Cruz, Leonardo de Oliveira Martins, Debora Keller, Stephen Rothery, Michele Becce, Anthony E G Cass, Chris Bakal, Ciro Chiappini, and Molly M Stevens. Nanoneedle-mediated stimulation of cell mechanotransduction machinery. ACS Nano, 13(3):2913–2926, 26 March 2019.

[33] Hsin-Ya Lou, Wenting Zhao, Xiao Li, Liting Duan, Alexander Powers, Matthew Akamatsu, Francesca Santoro, Allister F McGuire, Yi Cui, David G Drubin, and Bianxiao Cui. Membrane curvature underlies actin reorga-nization in response to nanoscale surface topography. Proc. Natl. Acad. Sci. U. S. A., 116(46):23143–23151, 12 November 2019.

[34] Kexin Zhu, Xiangfu Guo, Aravind Chandrasekaran, Xinwen Miao, Padmini Rangamani, Wenting Zhao, and Yansong Miao. Membrane curvature catalyzes actin nucleation through nano-scale condensation of N-WASP-FBP17. bioRxivorg, page 2024.04.25.591054, 25 April 2024.

[35] Dong-Hwee Kim, Bo Li, Fangwei Si, Jude M Phillip, Denis Wirtz, and Sean X Sun. Volume regulation and shape bifurcation in the cell nucleus. J. Cell Sci., 128(18):3375–3385, 15 September 2015.

[36] Sreenath Balakrishnan, Suma S Mathad, Geetika Sharma, Shilpa R Raju, Uma B Reddy, Saumitra Das, and G K Ananthasuresh. A nondimensional model reveals alterations in nuclear mechanics upon hepatitis C virus replication. Biophys. J., 116(7):1328–1339, 2 April 2019.

[37] A Vaziri, H Lee, and M R Kaazempur Mofrad. Deformation of the cell nucleus under indentation: Mechanics and mechanisms. J. Mater. Res., 21(8):2126–2135, 1 August 2006.

[38] Ashkan Vaziri and Mohammad R Kaazempur Mofrad. Mechanics and deformation of the nucleus in micropipette aspiration experiment. J. Biomech., 40(9):2053–2062, 1 January 2007.

[39] Gur Fabrikant, Soumya Gupta, G V Shivashankar, and Michael M Kozlov. Model of T-cell nuclear deformation by the cortical actin layer. Biophys. J., 105(6):1316–1323, 17 September 2013.

[40] Francesca Ballatore, Anotida Madzvamuse, Cécile Jebane, Emmanuèle Helfer, and Rachele Allena. A geometric-surface PDE model for cell-nucleus translocation through confinement. bioRxiv, page 2025.12.18.695144, 20 December 2025.

[41] Dan Deviri, Dennis E Discher, and Sam A Safran. Rupture dynamics and chromatin herniation in deformed nuclei. Biophys. J., 113(5):1060–1071, 5 September 2017.

[42] Ji-Eun Park, Juhyeon Jo, Kun Xu, Sun-Ah Lee, Seong-Beom Han, Yigji Lee, Won-Ki Cho, Bo Li, Soo Hyun Kim, and Dong-Hwee Kim. Attenuated nuclear tension regulates progerin-induced mechanosensitive nuclear wrinkling and chromatin remodeling. Adv. Sci. (Weinh.), 12(31):e2502375, 1 August 2025.

[43] Ann E Cowan, Ion I Moraru, James C Schaff, Boris M Slepchenko, and Leslie M Loew. Spatial modeling of cell signaling networks. Methods Cell Biol., 110:195–221, 2012.

[44] Emmet A Francis, Justin G Laughlin, Jørgen S Dokken, Henrik N T Finsberg, Christopher T Lee, Marie E Rognes, and Padmini Rangamani. Spatial modeling algorithms for reactions and transport in biological cells. Nat. Comput. Sci., 5(1):76–89, January 2025.

[45] Alessandro Contri, André Massing, and Padmini Rangamani. A finite element framework for solving coupled multiphysics problems with moving boundaries in cell biophysics. Comput. Methods Appl. Mech. Eng., 459(119071):119071, 1 September 2026.

[46] Alessandro Contri, Emmet A Francis, André Massing, and Padmini Rangamani. Mechanochemical feedback between cell shape and intracellular mechanics revealed by a finite-element framework. bioRxiv, page 2026.07.03.736361, 6 July 2026.

[47] R W Ogden. Non-Linear Elastic Deformations. Dover Publications, New York, NY, 2013.

[48] Larry A Taber. Nonlinear theory of elasticity: Applications in biomechanics: Applications in biomechanics. World Scientific Publishing, Singapore, Singapore, 23 February 2004.

[49] Dan Deviri and Samuel A Safran. Balance of osmotic pressures determines the nuclear-to-cytoplasmic volume ratio of the cell. Proc. Natl. Acad. Sci. U. S. A., 119(21):e2118301119, 24 May 2022.

[50] Dong-Hwee Kim, Jungwon Hah, and Denis Wirtz. Mechanics of the cell nucleus. Adv. Exp. Med. Biol., 1092:41–55, 2018.

[51] Irena L Ivanovska, Michael P Tobin, Tianyi Bai, Lawrence J Dooling, and Dennis E Discher. Small lipid droplets are rigid enough to indent a nucleus, dilute the lamina, and cause rupture. J. Cell Biol., 222(8):e202208123, 7 August 2023.

[52] Shyam B Khatau, Christopher M Hale, P J Stewart-Hutchinson, Meet S Patel, Colin L Stewart, Peter C Searson, Didier Hodzic, and Denis Wirtz. A perinuclear actin cap regulates nuclear shape. Proc. Natl. Acad. Sci. U. S. A., 106(45):19017–19022, 10 November 2009.

[53] Jau-Ye Shiu, Lina Aires, Zhe Lin, and Viola Vogel. Nanopillar force measurements reveal actin-cap-mediated YAP mechanotransduction. Nat. Cell Biol., 20(3):262–271, 5 March 2018.

[54] Andrea Rothballer, Thomas U Schwartz, and Ulrike Kutay. LINCing complex functions at the nuclear envelope: what the molecular architecture of the LINC complex can reveal about its function: What the molecular architecture of the LINC complex can reveal about its function. Nucleus, 4(1):29–36, 1 January 2013.

[55] Emma Carley, Rachel M Stewart, Abigail Zieman, Iman Jalilian, Diane E King, Amanda Zubek, Samantha Lin, Valerie Horsley, and Megan C King. The LINC complex transmits integrin-dependent tension to the nuclear lamina and represses epidermal differentiation. Elife, 10(e58541):e58541, 29 March 2021.

[56] Jan Lammerding, Jorine Eeftens, Gregg G Gundersen, Megan C King, Tanmay P Lele, Song Li, Gant Luxton G W, Ohad Medalia, Matthieu Piel, Pere Roca-Cusachs, Jörg Renkawitz, Kyle Roux, Daniel A Starr, Amy R Strom, Hawa Racine Thiam, Sara A Wickström, and Howard J Worman. Nuclear mechanobiology: a brief history and five unresolved questions. Nucleus, 17(1):2709894, 24 December 2026.

[57] Atsushi Matsuda and Mohammad R K Mofrad. Role of pore dilation in molecular transport through the nuclear pore complex: Insights from polymer scaling theory. PLoS Comput. Biol., 21(4):e1012909, 7 April 2025.

[58] N Daigle, J Beaudouin, L Hartnell, G Imreh, E Hallberg, J Lippincott-Schwartz, and J Ellenberg. Nuclear pore complexes form immobile networks and have a very low turnover in live mammalian cells. J. Cell Biol., 154(1):71–84, 9 July 2001.

[59] J L Broers, B M Machiels, G J van Eys, H J Kuijpers, E M Manders, R van Driel, and F C Ramaekers. Dynamics of the nuclear lamina as monitored by GFP-tagged a-type lamins. J. Cell Sci., 112 ( Pt 20)(20):3463– 3475, 15 October 1999.

[60] Shuaiyu Liu, Fangyuan Xiong, Zhen Dou, Lingluo Chu, Yihan Yao, Ming Wang, Xuebiao Yao, Xing Liu, and Zhikai Wang. Phosphorylation of lamin a/C regulates the structural integrity of the nuclear envelope. J. Biol. Chem., 301(1):108033, 1 January 2025.

[61] Dan Deviri, Charlotte R Pfeifer, Lawrence J Dooling, Irena L Ivanovska, Dennis E Discher, and Samuel A Safran. Scaling laws indicate distinct nucleation mechanisms of holes in the nuclear lamina. Nat. Phys., 15(8):823–829, 6 August 2019.

[62] Amy J Prunuske and Katharine S Ullman. The nuclear envelope: form and reformation. Curr. Opin. Cell Biol., 18(1):108–116, February 2006.

[63] Natalie Y Chen, Ye Yang, Thomas A Weston, Jason N Belling, Patrick Heizer, Yiping Tu, Paul Kim, Lovelyn Edillo, Steven J Jonas, Paul S Weiss, Loren G Fong, and Stephen G Young. An absence of lamin B1 in migrating neurons causes nuclear membrane ruptures and cell death. Proc. Natl. Acad. Sci. U. S. A., 116(51):25870– 25879, 17 December 2019.

[64] Luv Kishore Srivastava and Allen J Ehrlicher. Sensing the squeeze: nuclear mechanotransduction in health and disease. Nucleus, 15(1):2374854, 31 December 2024.

[65] G I Bell. Models for the specific adhesion of cells to cells: A theoretical framework for adhesion mediated by reversible bonds between cell surface molecules. Science, 200(4342):618–627, 12 May 1978.

[66] Francisco Merino-Casallo, Maria Jose Gomez-Benito, Silvia Hervas-Raluy, and Jose Manuel Garcia-Aznar. Unravelling cell migration: defining movement from the cell surface. Cell Adh. Migr., 16(1):25–64, December 2022.

[67] Brendon M Baker and Christopher S Chen. Deconstructing the third dimension: how 3D culture microenvironments alter cellular cues. J. Cell Sci., 125(Pt 13):3015–3024, 1 July 2012.

[68] Anna Pawluchin and Milos Galic. Moving through a changing world: Single cell migration in 2D vs. 3D. Front. Cell Dev. Biol., 10:1080995, 20 December 2022.

[69] David H Johnson, Orianna H Kou, Nicoletta Bouzos, and Wade F Zeno. Protein-membrane interactions: sensing and generating curvature. Trends Biochem. Sci., 49(5):401–416, 1 May 2024.

[70] Carole Arnold, Ismail Tahmaz, Marie-Ly Chapon, Hasna Maayouf, Valeriy Luchnikov, Jean-Louis Milan, Fabricio Borghi, and Laurent Pieuchot. Bending the rules: curvature’s impact on cell biology. BMC Biol., 23(1):296, 6 October 2025.

[71] Stefan Stöberl, Johannes Flommersfeld, Maximilian M Kreft, Martin Benoit, Chase P Broedersz, and Joachim O Rädler. Nuclear deformation and dynamics of migrating cells in 3D confinement reveal adaptation of pulling and pushing forces. Sci. Adv., 10(34):eadm9195, 23 August 2024.

[72] Martin Alnæs, Jan Blechta, Johan Hake, August Johansson, Benjamin Kehlet, Anders Logg, Chris Richardson, Johannes Ring, Marie E Rognes, and Garth N Wells. The FEniCS project version 1.5. Archive of Numerical Software, 3, 2015.

[73] Sirio Dupont, Leonardo Morsut, Mariaceleste Aragona, Elena Enzo, Stefano Giulitti, Michelangelo Cordenonsi, Francesca Zanconato, Jimmy Le Digabel, Mattia Forcato, Silvio Bicciato, Nicola Elvassore, and Stefano Piccolo. Role of YAP/TAZ in mechanotransduction. Nature, 474(7350):179–183, 8 June 2011.

[74] Tito Panciera, Luca Azzolin, Michelangelo Cordenonsi, and Stefano Piccolo. Mechanobiology of YAP and TAZ in physiology and disease. Nat. Rev. Mol. Cell Biol., 18(12):758–770, 27 December 2017.

[75] Spencer C Wei, Laurent Fattet, Jeff H Tsai, Yurong Guo, Vincent H Pai, Hannah E Majeski, Albert C Chen, Robert L Sah, Susan S Taylor, Adam J Engler, and Jing Yang. Matrix stiffness drives epithelial-mesenchymal transition and tumour metastasis through a TWIST1-G3BP2 mechanotransduction pathway. Nat. Cell Biol., 17(5):678–688, May 2015.

[76] Ashutosh Agrawal and Tanmay P Lele. Mechanics of nuclear membranes. J. Cell Sci., 132(14):jcs229245, 15 July 2019.

[77] Mehdi Torbati, Tanmay P Lele, and Ashutosh Agrawal. Ultradonut topology of the nuclear envelope. Proc. Natl. Acad. Sci. U. S. A., 113(40):11094–11099, 4 October 2016.

[78] Shalaka Patil and Kundan Sengupta. Role of a- and B-type lamins in nuclear structure-function relationships. Biol. Cell, 113(7):295–310, 1 July 2021.

[79] Jerome Irianto, Joe Swift, Rui P Martins, Graham D McPhail, Martin M Knight, Dennis E Discher, and David A Lee. Osmotic challenge drives rapid and reversible chromatin condensation in chondrocytes. Biophys. J., 104(4):759–769, 19 February 2013.

[80] Baihui Wang, Rafael Kronenberg-Tenga, Valentina Rosti, Emanuele Di Patrizio Soldateschi, Qiang Luo, Ugo Maria Iannacchero, Louise Pinet, Matthias Eibauer, Rajaa Boujemaa-Paterski, Benjamin Schuler, Chiara Lanzuolo, and Ohad Medalia. The molecular basis of lamin-specific chromatin interactions. Nat. Struct. Mol. Biol., 32(10):1999–2011, 1 October 2025.

[81] Lukasz Suprewicz, Fitzroy J Byfield, Thomas T Dutta, and Paul A Janmey. Role of nuclear ATPases in nuclear mechanics and cell migration through confined spaces: Opposite effects of BRG1 and cohesin. Biophys. J., 14 February 2026.

[82] Fitzroy J Byfield, Behnaz Eftekhari, Kaeli Kaymak-Loveless, Kalpana Mandal, David Li, Rebecca G Wells, Wenjun Chen, Jasna Brujic, Giulia Bergamaschi, Gijs J L Wuite, Alison E Patteson, and Paul A Janmey. Metabolically intact nuclei are fluidized by the activity of the chromatin remodeling motor BRG1. Biophys. J., 124(3):494–507, 4 February 2025.

[83] Jing Yao, Ying Fan, Yuanke Li, and Leaf Huang. Strategies on the nuclear-targeted delivery of genes. J. Drug Target., 21(10):926–939, 22 December 2013.

[84] Howard J Worman. Nuclear lamins and laminopathies. J. Pathol., 226 (2):316–325, January 2012.

[85] Subarna Dutta, Jitendra K Das, Lakshmi Maganti, Maitree Bhattacharyya, Dhananjay Bhattacharyya, Sujoy Mukherjee, and Kaushik Sengupta. Skeletal muscle dystrophy mutant of lamin a alters the structure and dynamics of the ig fold domain. Sci. Rep., 8(1):13793, 14 September 2018.

[86] Elin Torvaldson, Vitaly Kochin, and John E Eriksson. Phosphorylation of lamins determine their structural properties and signaling functions. Nucleus, 6(3):166–171, 20 March 2015.

[87] Ashley J Earle, Tyler J Kirby, Gregory R Fedorchak, Philipp Isermann, Jineet Patel, Sushruta Iruvanti, Steven A Moore, Gisèle Bonne, Lori L Wallrath, and Jan Lammerding. Mutant lamins cause nuclear envelope rupture and DNA damage in skeletal muscle cells. Nat. Mater., 19(4):464–473, April 2020.

[88] Atsuki En, Hanumakumar Bogireddi, Briana Thomas, Alexis V Stutzman, Sachie Ikegami, Brigitte LaForest, Omar Almakki, Peter Pytel, Ivan P Moskowitz, and Kohta Ikegami. Pervasive nuclear envelope ruptures precede ECM signaling and disease onset without activating cGAS-STING in Lamin-cardiomyopathy mice. Cell Rep., 43(6):114284, 25 June 2024.

[89] Winnok H De Vos, Frederik Houben, Miriam Kamps, Ashraf Malhas, Fons Verheyen, Juliën Cox, Erik M M Manders, Valerie L R M Verstraeten, Maurice A M van Steensel, Carlo L M Marcelis, Arthur van den Wijngaard, David J Vaux, Frans C S Ramaekers, and Jos L V Broers. Repetitive disruptions of the nuclear envelope invoke temporary loss of cellular compartmentalization in laminopathies. Hum. Mol. Genet., 20(21):4175–4186, 1 November 2011.

[90] Emmet A Francis, Justin Laughlin, Henrik Finsberg, Jørgen Schartum Dokken, Yuan Gao, Marie E Rognes, Christopher Lee, Will Xu, and JJ. RangamaniLabUCSD/smart: SMART branch for mechanochemical model of nucleus (DOI 10.5281/zenodo.18654652), 2026.

[91] Emmet A Francis. RangamaniLabUCSD/smart-mechanotransduction: v1.0.0 (DOI 10.5281/ZENODO.18690946), 2026.

[92] Aanya Sawhney. 20260922 code and compiled data - francis, sawhney, et al. mechanochemical modeling of the nuclear lamina (DOI: 10.5281/ZENODO.22905725), 2026.

[93] San Diego Supercomputer Center. Triton shared computing cluster, 2022.

