## Supporting Information for "A predictive mechanochemical modeling framework for the deformation and remodeling of the nuclear lamina"

#### S1 Model Formulation

##### S1.1 Hyperelastic model of the nuclear envelope

In line with previous studies, we model the NE as a hyperelastic shell with finite thickness [1, 2, 3]. This approach lumps the double bilayer with the underlying nuclear lamina consisting largely of a network of lamin A/C. The stress-free configuration is a spherical shell with center at [0, 0, $z_{\text{nuc}} + R_0$], outer radius $R_0$, and thickness $\Delta T$,

$$X^2 + Y^2 + (Z - z_{\text{nuc}} - R_0)^2 = R_0^2 \quad \text{on } \Gamma_{\text{outer}}, \tag{S1}$$

$$X^2 + Y^2 + (Z - z_{\text{nuc}} - R_0)^2 = (R_0 - \Delta T)^2 \quad \text{on } \Gamma_{\text{inner}}, \tag{S2}$$

where $\mathbf{X} = [X, Y, Z]$ are the undeformed material coordinates (Figure S1A). The volume between $\Gamma_{\text{inner}}$ and $\Gamma_{\text{outer}}$ is $\Omega_{\text{NE}}$ (Figure S1B). $\mathbf{N}$, the unit normal of the reference geometry (Figure S1A), can be expressed analytically over any slice of the NE with constant radius,

$$\mathbf{N} = \frac{1}{\sqrt{X^2 + Y^2 + (Z - z_{\text{nuc}} - R_0)^2}} \left[X, Y, Z - z_{\text{nuc}} - R_0\right]. \tag{S3}$$

Given a displacement field $\mathbf{u}$ defined within $\Omega_{\text{NE}}$ resulting in new deformed coordinates $\mathbf{x} = \mathbf{X} + \mathbf{u}$, we define conventional displacement metrics including the deformation gradient tensor,

$$\mathbf{F} = \left(\frac{\partial \mathbf{x}}{\partial \mathbf{X}}\right)^T, \tag{S4}$$

and the right Cauchy-Green tensor,

$$\mathbf{C} = \mathbf{F}^T \cdot \mathbf{F}. \tag{S5}$$

These tensors are associated with the following strain invariants,

$$I_1 = \text{tr } \mathbf{C}, \tag{S6}$$

$$I_2 = \frac{1}{2}\left[(\text{tr } \mathbf{C})^2 - \text{tr}(\mathbf{C} \cdot \mathbf{C})\right], \tag{S7}$$

$$J = \det \mathbf{F}, \tag{S8}$$

where the final invariant is the Jacobian of transformation representing volume dilatation. The strain energy functional is given in terms of these invariants assuming an incompressible Mooney-Rivlin material as in [1],

$$W = E_1(I_1 - 3) + E_2(I_2 - 3) - p(J - 1), \tag{S9}$$

where $E_1$ and $E_2$ are elastic constants (values for these and other parameters given in Table S1) and $p$ is a pressure-like variable that acts as a Lagrange multiplier to enforce incompressibility within the shell. The first Piola-Kirchoff stress tensor ($\mathbf{t}$) and the Cauchy stress tensor ($\boldsymbol{\sigma}$) can then be computed by

$$\mathbf{t} = \frac{\partial W}{\partial \mathbf{F}^T}, \tag{S10}$$

$$\boldsymbol{\sigma} = \frac{1}{J}\mathbf{F} \cdot \frac{\partial W}{\partial \mathbf{F}^T}. \tag{S11}$$

We also consider the surface tension at the outer edge of the NE as a scalar output for the Cauchy stress tensor,

$$\gamma = \frac{\Delta \hat{z}}{2}(\sigma_{\gamma 1} + \sigma_{\gamma 2}) \tag{S12}$$

$$\text{where} \quad \sigma_{\gamma 1} = \frac{(\mathbf{e}_\Theta \cdot \mathbf{t}) \cdot \mathbf{e}_\Theta}{\alpha_\Theta} \tag{S13}$$

$$\text{and} \quad \sigma_{\gamma 2} = \frac{(\mathbf{e}_\Phi \cdot \mathbf{t}) \cdot \mathbf{e}_\Phi}{\alpha_\Phi}. \tag{S14}$$

The unit vectors in the $\Theta$ and $\Phi$ directions (Figure S1A), as well as the stretch in the associated planes are given by

$$\mathbf{e}_\Theta = \frac{1}{\sqrt{X^2 + Y^2}}[-Y, X, 0], \tag{S15}$$

$$\mathbf{e}_\Phi = \mathbf{e}_\Theta \times \mathbf{N}, \tag{S16}$$

$$\alpha_\Theta = |J\mathbf{e}_\Phi \cdot \mathbf{F}^{-1}|, \tag{S17}$$

$$\alpha_\Phi = |J\mathbf{e}_\Theta \cdot \mathbf{F}^{-1}|. \tag{S18}$$

Assuming mechanical equilibrium and no body forces, the following equation must be satisfied:

$$\nabla \cdot \mathbf{t} = 0 \quad \text{in } \Omega_{\text{NE}}, \tag{S19}$$

We assume the center of the nucleus is positioned directly above a central nanopillar, leading to a symmetry condition about the $x$ and $y$ axes,

$$u_y = 0 \quad \text{for } x = 0, \tag{S20}$$

$$u_x = 0 \quad \text{for } y = 0, \tag{S21}$$

such that we only need to simulate a quarter of the nuclear geometry. Any region of the NE that contacts the lower region of the substrate experiences an additional Dirichlet condition:

$$u_z = -h_{\text{NP}} - Z \quad \text{for } z \leq -h_{\text{NP}}, \tag{S22}$$

where coordinates are defined such that the bottom of the NE rests at $z = -h_{\text{NP}}$.

To model nanopillar-mediated indentation, we apply a repulsive potential acting between the PM adhered to the nanopillar surface and the outer NE,

$$\sigma_{\text{contact}} = \begin{cases} \sigma_{\text{contact},0} \exp\left(-\dfrac{Z + u_z - d_{\text{steric}}}{d_{\text{steric}}}\right) & \text{on } \Gamma_{\text{contact}} \\ 0 \quad \text{otherwise} \end{cases}, \tag{S23}$$

where $\Gamma_{\text{contact}}$ is the portion of the outer NE directly above a nanopillar contact (Figure S1B, Equation (S47)) and $d_{\text{steric}}$ is the characteristic distance associated with steric repulsion. This provides a Robin boundary condition for the $z$-displacement, which is supplemented by an additional Dirichlet boundary condition for the $z$-displacement at the cap that was found to aid in numerical stability (Equation (S50)).

The above Robin boundary condition is grouped with other stress-type boundary conditions including stress on the outer NE ($\Gamma_{\text{outer}}$) due to the perinuclear actin cap and stress at the interior nuclear surface ($\Gamma_{\text{inner}}$) due to an osmotic pressure gradient. To apply these conditions, we note that the surface traction at the inner and outer boundaries of the NE is given by

$$\sigma_{\text{NE}} = \frac{\mathbf{N} \cdot \mathbf{t}}{\alpha}, \tag{S24}$$

where $\alpha$ is the local stretch ratio of the nuclear surface, needed to convert the psuedotraction stress to a true traction over the deformed surface [4]. Accordingly,

$$\mathbf{N} \cdot \mathbf{t} = \begin{cases} -\alpha(\sigma_{\text{cap,tot}} \mathbf{e}_z + P_{\text{cyto}} \mathbf{n}) & \text{on } \Gamma_{\text{outer}} \backslash \Gamma_{\text{contact}} \\ \alpha P_{\text{nuc}} \mathbf{n} & \text{on } \Gamma_{\text{inner}} \\ \alpha \left[(\sigma_{\text{contact}} - \sigma_{\text{cap,tot}}) \mathbf{e}_z - P_{\text{cyto}} \mathbf{n}\right] & \text{on } \Gamma_{\text{contact}} \end{cases}, \tag{S25}$$

where $\mathbf{e}_z$ is the unit vector in the $z$ direction and $\mathbf{n}$ is the unit normal associated with the deformed surface (Figure S1B). $\alpha$ and $\mathbf{n}$ are computed using Nanson's formula, which relates the area and unit normal ($\mathbf{N}$, Equation (S3)) in the reference configuration to those in the deformed configuration,

$$\alpha = |J \mathbf{N} \cdot \mathbf{F}^{-1}|, \tag{S26}$$

$$\mathbf{n} = \frac{J \mathbf{N} \cdot \mathbf{F}^{-1}}{\alpha}. \tag{S27}$$

$\sigma_{\text{cap}}$ varies over the geometry, reaching its maximum value at the top of the nucleus and always directed in the $-z$-direction,

$$\sigma_{\text{cap,tot}} = \sigma_{\text{cap}} \exp\left(\frac{Z - (z_{\text{nuc}} + 2R_0)}{Z_0}\right), \tag{S28}$$

where $\sigma_{\text{cap}}$ sets the force magnitude and $Z_0$ determines the spatial extent of the cap stress.

We also consider the osmotic stress gradient acting across the nuclear envelope. To initialize the model, we increase the pressure gradient ($P_{\text{nuc}} - P_{\text{cyto}}$) from 0 Pa to $\Delta P_{\text{max}}$, after which the pressure is determined by the scaling relationship between nuclear and cytosolic volumes and respective pressures as in [5],

$$P_{\text{nuc}} = P_{\text{nuc,other}} + \frac{E_{\text{solvent,nuc}}}{\text{vol}_{\text{nuc}} - \text{vol}_{\text{shell}} - \text{vol}_{\text{excl,nuc}}} \tag{S29}$$

$$P_{\text{cyto}} = P_{\text{cyto,other}} + \frac{E_{\text{solvent,cyto}}}{\text{vol}_{\text{cyto}} - \text{vol}_{\text{excl,cyto}}}. \tag{S30}$$

The volume of the nucleus, $\text{vol}_{\text{nuc}}$, is solved for in our model, and the volume of the NE-lamin composite, $\text{vol}_{\text{shell}}$ remains constant over time due to the incompressibility constraint. Assuming conserved total cell volume, we can solve for the cytosolic volume at each time step as well, $\text{vol}_{\text{cyto}} = \text{vol}_{\text{tot}} - \text{vol}_{\text{nuc}}$, and the excluded volumes

($\text{vol}_{\text{excl,nuc}}$ and $\text{vol}_{\text{excl,cyto}}$) are given by estimates in [5] and assumed constant over time (Table S1). The total cell volume is set according to the average relative nuclear volume measured in our experiments (0.363), such that $\text{vol}_{\text{nuc},0} = 0.363\text{vol}_{\text{tot}}$ (Table S1). $P_{\text{nuc,other}}$ and $P_{\text{cyto,other}}$ represent pressure-like stresses due to chromatin, microtubules, and other factors. We kept $\Delta P_{\text{other}} = 0$ Pa through this study, as these stresses were previously estimated to be much smaller than osmotic stresses or cytoskeletal stresses [5]. $E_{\text{solvent,nuc}}$ and $E_{\text{solvent,nuc}}$ are energies associated with confined solute molecules in each compartment and are assumed constant over time [5]. We constrained these energies to match the assumed pressure difference at rest, $\Delta P_{\max}$:

$$E_{\text{solvent,nuc}} = (\text{vol}_{\text{nuc},0} - \text{vol}_{\text{shell}} - \text{vol}_{\text{excl,nuc}})(P_0 + \Delta P_{\max}) \tag{S31}$$

$$E_{\text{solvent,cyto}} = (\text{vol}_{\text{cyto},0} - \text{vol}_{\text{excl,cyto}})P_0 \tag{S32}$$

$P_0$, the osmotic pressure on each side in the absence of NE tension or other forces, was previously estimated at 8000 Pa. We tested values between 1000 and 10000 Pa before fixing $P_0$ to 2400 Pa. This was chosen to result in up to 10% volume loss in our simulations, suggested by previous experiments measuring a 10% increase in nuclear volume following latrunculin B treatment [2] or confined cell migration [6] (Figure S2).

To solve for the initial configuration of the nucleus with applied pressure gradient $\Delta P_{\max}$ prior to applying any nanopillar or cap stress, it is necessary to include a Dirichlet boundary condition in the $z$-direction to obtain a unique solution (otherwise $z$ displacement can only be determined up to a constant). In this case, we apply a symmetry condition about the middle of the nucleus:

$$u_z = 0 \quad \text{for } Z = z_{\text{nuc}} + R_0. \tag{S33}$$

**Table S1:** Parameters governing nuclear mechanics.

| | Value | Description | Sources/Notes |
|---|---|---|---|
| $R_0$ | 4.1 µm | Stress-free nuclear radius | 5.0 µm post-inflation [7]* |
| $\Delta T$ | 0.2 µm | Stress-free NE thickness | 0.13 µm post-inflation [8]** |
| $z_{\text{nuc}}$ | 1.04 µm | Initial location of the NE bottom | Ensures substrate contact post-inflation |
| $Z_0$ | 2.05 µm | Characteristic decay length for cap stress | Restricts to mainly upper half of NE |
| $\Delta P_{\max}$ | 820 Pa | Pressure difference across the NE | [1, 2] |
| $P_0$ | 2400 Pa | Baseline osmotic pressure | [5] |
| $E_1$ | 5000 Pa | First Mooney-Rivlin elastic constant | [1] |
| $E_2$ | 1000 Pa | Second Mooney-Rivlin elastic constant | [1] |
| $t_{0,\text{cap}}$ | 1000 s | Time scale for perinuclear actin cap formation | |
| $\sigma_{\max}$ | 0-800 Pa | Maximum perinuclear actomyosin force | Varied for testing |
| $\phi_{\text{inner}}$ | 100 Pa | Nucleoplasm bulk modulus | Constrained by volume loss [9, 2] |
| $\phi_0$ | $1 \times 10^8$ Pa | Bulk modulus (incompressibility constraint) | Set by numerical testing |
| $\sigma_{\text{contact},0}$ | 1000 Pa | Contact repulsion strength | Set by numerical testing |
| $d_{\text{steric}}$ | 0.2 µm | Steric repulsion radius | Set by numerical testing |
| $h_{\text{edge}}$ | 0.15 µm | Mesh resolution on flat substrate | Set by numerical testing |
| | 0.1 µm | Mesh resolution on 200 nm nanopillars | Set by numerical testing |
| | 0.15 µm | Mesh resolution on 500 nm nanopillars | Set by numerical testing |
| $D_{\text{roof}}$ | $10h_{\text{edge}}$ | Upper region cutoff (Equation (S51)) | Set by numerical testing |
| $E_{1,\text{nuc}}$ | 5000 Pa | Elastic modulus for nucleoplasm deformation | Set by numerical testing |
| $\phi_{\text{nuc}}$ | $1 \times 10^5$ Pa | Bulk modulus for nucleoplasm deformation | Set by numerical testing |
| $\Delta\hat{z}$ | 50 nm | Double bilayer plus perinuclear space thickness | [10] |

* Post-inflation radius matches nuclear size from [7]
** Post-inflation thickness is at upper end of double membrane plus laminar thickness [8]

#### S1.2 Biochemical reaction and transport

We consider the reaction-transport of several molecular species with the NE (lamin A/C and NPCs) or within the nucleoplasm (YAP/TAZ and phosphorylated lamin A/C). Dephosphorylated lamin A/C is defined on the inner surface ($\Gamma_{\text{inner}}$), whereas NPC species are defined on the outer NE surface ($\Gamma_{\text{outer}}$). However, because we assume that all interactions between lamins, NPCs, YAP/TAZ, actin, and myosin occur at the outer surface of the NE, we track the projected lamin concentration on $\Gamma_{\text{outer}}$ as detailed below. We do not explicitly include the cytosolic space in our model, but we do consider the cytosolic concentrations of F-actin ($[F]$), activated myosin ($[M]_{\text{A}}$), and free cytosolic YAP/TAZ ($[Y]_{\text{free}}$) at the NE interface with the cytosol. We assume cytosolic YAP/TAZ and activated myosin are well-mixed, as YAP/TAZ diffuses rapidly in the cytosol [11, 12] and myosin activation is determined by Rho-associated protein kinase in our previous models, which also diffuses rapidly in the cytosol [13]. Myosin is fixed according to values from our previous simulations [14], and cytosolic YAP/TAZ species are computed at each time point via Equations (S86) and (S87). F-actin concentration is expressed as a function of distance between the NE and the PM ($d_{\text{NE-PM}}$) and of curvature of the closest point on the PM ($H_{\text{PM}}$) to capture the trends modeled in our previous work [14]:

$$[F] = [F]_0 + [F]_1 \exp\left(-\frac{d_{\text{NE-PM}}}{d_F}\right)\exp\left(\frac{H_{\text{PM}}}{H_0}\right). \tag{S34}$$

$[F]_0$ (globally averaged F-actin), $[F]_1$ (curvature-dependent F-actin assembly), $d_F$, and $H_0$ were constrained according to our previous model; values and definitions for these parameters and all others are provided in Table S2. Global F-actin concentration, $[F]_0$, varies according to the nanopillar radius to capture the curvature-dependent inhibition of focal adhesion formation shown in previous experiments [15, 16] and included in our previous model [14]. The PM-NE distance and curvature of the PM were computed assuming an ideal cylindrical geometry of the PM around the nanopillar and only considering the closest nanopillar to a given point on the NE as fully detailed in Equation (S90).

Given these cytosolic variables, the reaction-transport equation governing dephosphorylated lamin in the NE ($[L]$) is

$$\frac{\partial [L]}{\partial t} + \nabla_{\text{S,inner}} \cdot (\mathbf{v}_{\text{NE,inner}}[L]) = D_L \nabla^2_{\text{S,inner}}[L] - k_{\text{rl}}[L] + k_{\text{fl}}\frac{p_{\text{cyto}}[F]^{2.6}}{C_L + p_{\text{cyto}}[F]^{2.6}}[L]_{\text{phos}} \quad \text{on} \quad \Gamma_{\text{outer}}. \tag{S35}$$

Note that while the surface differential operator ($\nabla_{\text{S,inner}}$) and the advection velocity ($\mathbf{v}_{\text{NE,inner}}$) are defined according to the motion of the inner surface $\Gamma_{\text{inner}}$, the concentration itself is computed on the outer surface $\Gamma_{\text{outer}}$, allowing all reactions to occur locally over this interface. $\nabla_{\text{S,inner}}$ and $\mathbf{v}_{\text{NE,inner}}$ are defined in Section S2.3. Lamin diffuses with coefficient $D_L$ (fixed to zero based on previous experiments [17]) and phosphorylation and dephosphorylation are described using relations from our previous work [18] (see definitions and values of $k_{\text{rl}}$, $k_{\text{fl}}$, $p_{\text{cyto}}$, and $C_{\text{L}}$ in Table S2). In contrast to our previous model, we consider phosphorylated lamin concentration ($[L]_{\text{phos}}$) within the nuclear volume, as it is assumed disassembled and effectively solubilized [19, 20],

$$\frac{\partial [L]_{\text{phos}}}{\partial t} + \nabla \cdot (\mathbf{v}_{\text{nuc}}[L]_{\text{phos}}) = D_{L,\text{vol}} \nabla^2 [L]_{\text{phos}} \quad \text{in} \quad \Omega_{\text{full}}, \tag{S36}$$

$$D_{L,\text{vol}} \nabla [L]_{\text{phos}} \cdot \mathbf{n} = \frac{1}{N_{\text{conv}}}\left(k_{\text{rl}}[L] - k_{\text{fl}}\frac{p_{\text{cyto}}[F]^{2.6}}{C_L + p_{\text{cyto}}[F]^{2.6}}[L]_{\text{phos}}\right) \quad \text{on} \quad \Gamma_{\text{outer}}, \tag{S37}$$

where $\mathbf{v}_{\text{nuc}}$ is the extrapolated velocity field within the nucleus (see Section S2) and $D_{\text{L,vol}}$ is the diffusion coefficient of phosphorylated lamin A/C in the nucleoplasm. $\Omega_{\text{full}}$ includes both volumetric domains ($\Omega_{\text{full}} = \Omega_{\text{nuc}} \cup \Omega_{\text{NE}}$), which are combined for the case of volume species in the biochemical model.

Dilated ($[NPC]_{\text{dilated}}$) and constricted ($[NPC]$) NPCs are also included in our framework, advected with the outer surface of the NE (velocity $\mathbf{v}_{\text{NE,outer}}$) and assumed to be nondiffusive [21],

$$\frac{\partial [NPC]}{\partial t} + \nabla_S \cdot (\mathbf{v}_{\text{NE,outer}}[NPC]) = -k_{\text{fNPC}}[F][L][M]_A[NPC] + k_{\text{rNPC}}[NPC]_{\text{dilated}} \text{ on } \Gamma_{\text{outer}}, \tag{S38}$$

$$\frac{\partial [NPC]_{\text{dilated}}}{\partial t} + \nabla_S \cdot (\mathbf{v}_{\text{NE,outer}}[NPC]_{\text{dilated}}) = k_{\text{fNPC}}[F][L][M]_A[NPC] - k_{\text{rNPC}}[NPC]_{\text{dilated}} \text{ on } \Gamma_{\text{outer}}, \tag{S39}$$

where $k_{\text{fNPC}}$ and $k_{\text{rNPC}}$ are the rates of NPC dilation and constriction, respectively. $\nabla_S$ is the surface gradient operator as defined in Equation (S58).

The resulting transport of YAP/TAZ across the NE depends on nuclear stretch, such that the maximum rate of transport is a function of outer NE stretch $\alpha$ (Equation (S26)),

$$k_{\text{in,tot}} = \phi_{\text{scale}} \left( k_{\text{in,b}} + k_{\text{in}} [NPC]_{\text{dilated}} \right) \exp \left( \frac{\alpha - \alpha_{\text{init}}}{\alpha_0} \right), \tag{S40}$$

where $\alpha_{\text{init}}$ is the average NE stretch at $t = 0$ (after increasing the nuclear pressure), $k_{\text{in,b}}$ and $k_{\text{in}}$ are the baseline and NPC-dilation dependent import rates, and $\alpha_0$ determines the stretch sensitivity. $\phi_{\text{scale}}$ is a flux scaling factor included as a calibration parameter for this revised model. This results in the following equation and boundary condition for nuclear YAP/TAZ ($[Y]_{\text{nuc}}$):

$$\frac{\partial [Y]_{\text{nuc}}}{\partial t} + \nabla \cdot (\mathbf{v}_{\text{nuc}} [Y]_{\text{nuc}}) = D_Y \nabla^2 [Y]_{\text{nuc}} \quad \text{in } \Omega_{\text{nuc,tot}}, \tag{S41}$$

$$D_Y \nabla [Y]_{\text{nuc}} \cdot \mathbf{n} = \frac{1}{N_{\text{conv}}} \left( k_{\text{in,tot}} [Y]_{\text{free}} - k_{\text{out}} [Y]_{\text{nuc}} \right) \quad \text{on } \Gamma_{\text{outer}}, \tag{S42}$$

where $D_Y$ is the nuclear diffusion coefficient of YAP/TAZ and $k_{\text{out}}$ is the rate of YAP/TAZ efflux from the nucleus.

#### S1.3 Mechanochemical coupling

We introduced time-dependence into our mechanical system by writing the cap stress as a function of time (Eq 1). Solving for associated nuclear deformations at over a given time step then determines the velocities $\mathbf{v}_{\text{NE,outer}}$, $\mathbf{v}_{\text{NE,inner}}$, $\mathbf{v}_{\text{nuc}}$ (via extrapolation as described in Section S2.2) and the NE stretch $\alpha$, all of which appear in the above biochemical reaction-transport equations. We also consider coupling from the biochemical model to the mechanical model via lamin-dependent changes in NE stiffness. Based on measured correlations between lamin A/C abundance and nuclear bulk modulus [22], we scale both elastic moduli ($E_1$ and $E_2$ in Equation (S9)) by factor $E_{\text{scale}}$,

$$E_{\text{scale}} = (1 - \phi_L) + \phi_L \frac{[L]}{[L]_{\text{ref}}}, \tag{S43}$$

where $\phi_L$ is a fraction between 0 and 1 that determines the relative contribution of lamin A/C to NE stiffness.

As a main output relevant to mechanochemical coupling, we compute the force per lamin subunit. Although our model does not include the microscopic details of load sharing among the connected network of lamin filaments [23], this approach allows us to predict the local force felt by each dimer in a spatially averaged sense,

$$\mathbf{F}_L = \frac{\sigma_{\text{NE}}}{[L]}, \tag{S44}$$

where $\sigma_{\text{NE}}$ is the stress at the midplane of the NE in our simulations and $[L]$ is the projected lamin density.

**Table S2:** Reaction-transport parameters.

| | Value | Description | Source(s) |
|---|---|---|---|
| $d_F$ | 0.5 μm | Characteristic decay length for F-actin | * |
| $H_0$ | 2.0 μm | Sensitivity of actin polymerization to PM curvature | [15, 14] |
| $[F]_0$ | 140 μM | Baseline actin polymerization (flat substrate) | [15, 14] |
| | 100 μM | Baseline actin polymerization ($r_{\text{NP}} = 0.2$ μm) | [15, 14] |
| | 120 μM | Baseline actin polymerization ($r_{\text{NP}} = 0.5$ μm) | [15, 14] |
| $[F]_1$ | 20 μM | PM curvature-dependent actin polymerization | [14, 24] |
| $D_L$ | 0.0 μm$^2$ s$^{-1}$ | NE lamin diffusion coefficient | [17] |
| $D_{\text{L,vol}}$ | 4.0 μm$^2$ s$^{-1}$ | Nucleoplasmic lamin diffusion coefficient | [25]** |
| $k_{\text{rl}}$ | 0.001 s$^{-1}$ | Lamin phosphorylation rate | [26, 18] |

Continued on next page

| | Value | Description | Source(s) |
|---|---|---|---|
| $k_{\text{fl}}$ | 415.5 $\mu\text{m}^{-2}\mu\text{M}^{-1}\text{s}^{-1}$ | Maximum lamin dephosphorylation rate | [18, 27]*** |
| $p_{\text{cyto}}$ | $9 \times 10^{-6}$ kPa $\mu\text{M}^{-2.6}$ | Cytoplasmic stiffness coefficient | [28, 18] |
| $C_L$ | 100 kPa | Cytoplasmic stiffness sensitivity (lamin A dephos.) | [27, 18] |
| $k_{\text{fNPC}}$ | $2.8 \times 10^{-7}$ $\mu\text{m}^2$ $\text{s}^{-1}$ $\mu\text{M}^{-1}$ | NPC dilation rate | [18] |
| $k_{\text{rNPC}}$ | 8.7 $\text{s}^{-1}$ | NPC constriction rate | [18] |
| $k_{\text{in,b}}$ | 1.0 $\mu\text{m}^{-2}$ $\text{s}^{-1}$ $\mu\text{M}^{-1}$ | Baseline YAP/TAZ import rate | [18] |
| $k_{\text{in}}$ | 10.0 $\mu$M $\text{s}^{-1}$ | Lamin-dependent YAP/TAZ import through NPCs | [18] |
| $\alpha_0$ | 5.0 | NPC stretch sensitivity | [14] |
| $D_Y$ | 4.0 $\mu\text{m}^2$ $\text{s}^{-1}$ | YAP/TAZ diffusion rate in nucleus | [11] |
| $k_{\text{CN}}$ | 0.56 $\text{s}^{-1}$ | Baseline YAP/TAZ activation rate | [18] |
| $k_{\text{CY}}$ | $7.6 \times 10^{-4}$ $\mu\text{M}^{-2}$ $\text{s}^{-1}$ | Actomyosin-dependent YAP/TAZ activation | [18] |
| $k_{\text{NC}}$ | 0.14 $\text{s}^{-1}$ | YAP/TAZ sequestration rate | [18] |
| $\text{vol}_{\text{tot}}$ | 418.7 $\mu\text{m}^3$ | Total volume for U2OS (quarter cell) | |
| $N_{\text{NPC}}$ | 594.7 | Total number of NPCs | [18, 14] |
| $N_{\text{L}}$ | $3.202 \times 10^5$ | Total number of lamin A/C subunits (wild type) | [18, 14] |
| | $1.601 \times 10^5$ | Total number of lamin A/C subunits (low lamin) | [18, 14] |
| $N_{\text{Y}}$ | $5.339 \times 10^5$ | Total number of YAP/TAZ molecules | [18, 14] |
| $\phi_{\text{scale}}$ | 0.75 | Scale factor for YAP/TAZ influx | † |
| $\phi_{\text{L}}$ | 0.5 | Relative contribution of lamin A/C to NE stiffness | [22] |
| $[L]_{\text{ref}}$ | 2980 $\mu\text{m}^{-2}$ | Lamin stiffness normalization (matches WT density) | |
| $N_{\text{conv}}$ | 602.2 # $\mu\text{m}^{-3}$ $\mu\text{M}^{-1}$ | Conversion factor | |
| $k_{\text{rupture},0}$ | $6.151 \times 10^{-18}$ $\text{s}^{-1}$ | Lamina rupture rate at zero force | Table S3 |
| $\hat{\delta}$ | 65.0 nm | Lumped molecular displacement parameter | Table S3 |
| $k_B T$ | $4.1 \times 10^{-21}$ J | Thermal energy scale (room temperature) | |
| Initial conditions (determined by mechanical model): | | | |
| $\text{SA}_{\text{NE,inner},0}$ | 107.5 $\mu\text{m}^2$ | Initial surface area of inner NE | |
| $\text{SA}_{\text{NE,outer},0}$ | 118.4 $\mu\text{m}^2$ | Initial surface area of outer NE | |
| $\text{vol}_{\text{nuc},0}$ | 152.0 $\mu\text{m}^3$ | Initial volume of nucleus | |

$$[L]_{\text{phos,init}} = \frac{N_{\text{L}} k_{\text{rl}}}{k_{\text{rl}} \text{vol}_{\text{nuc},0} N_{\text{conv}} + k_{\text{fl}} \text{SA}_{\text{NE,inner},0}} \quad \text{(SS)}^{\dagger\dagger}$$

$$[L]_{\text{init}} = \frac{N_{\text{L}} - [L]_{\text{phos,init}} \text{vol}_{\text{nuc},0} N_{\text{conv}}}{\text{SA}_{\text{NE,inner},0}} \quad \text{(SS)}^{\dagger\dagger}$$

$$[NPC]_{\text{dilated,init}} = \frac{N_{\text{NPC}}}{SA_{\text{NE,outer},0}} \frac{k_{\text{fNPC}} [L]_{\text{init}} [F][M]_{\text{A}}}{k_{\text{fNPC}} [L]_{\text{init}} [F][M]_{\text{A}} + k_{\text{rNPC}}} \quad \text{(SS)}^{\dagger\dagger}$$

$$[NPC]_{\text{init}} = \frac{N_{\text{NPC}}}{SA_{\text{NE,outer},0}} - [NPC]_{\text{dilated,init}} \quad \text{(SS)}^{\dagger\dagger}$$

$$[Y]_{\text{nuc,init}} = \frac{N_{\text{Y}}}{N_{\text{conv}}} \frac{\frac{[Y]_{\text{free}}}{[Y]_{\text{cyto}}} \phi_{\text{scale}} \left(k_{\text{inb}} + k_{\text{in}} [NPC]_{\text{dilated,init}}\right)}{k_{\text{out}} (\text{vol}_{\text{tot}} - \text{vol}_{\text{nuc},0}) + \frac{[Y]_{\text{free}}}{[Y]_{\text{cyto}}} \phi_{\text{scale}} \left(k_{\text{inb}} + k_{\text{in}} [NPC]_{\text{dilated,init}}\right) \text{vol}_{\text{nuc},0}} \quad \text{(SS)}^{\dagger\dagger\dagger}$$

* Set to approximately match length scale from our previous simulations [14]
** Intermediate value used between two mobile populations of lamin A in the nucleoplasm in [25].
*** Converted from $k_{\text{fl}}$ in previous studies assuming nuclear volume-to-surface-area ratio of 1.5 $\mu$m
† Set to match values of YAP/TAZ N/C measured in our experiments.
†† Expressions given for predicted steady-state values in well-mixed limit.
††† Well-mixed steady-state expression uses $\frac{[Y]_{\text{free}}}{[Y]_{\text{cyto}}}$ from Equation (S87).

### S2 Numerical Methods

#### S2.1 Numerical approximation of hyperelastic nuclear deformation

To solve the hyperelastic equations via FEniCS, we recast them into their associated weak form:

$$\int \nabla\mathbf{v}\cdot\mathbf{t}d\Omega + \int q\left(\phi_0(J-1)+p\right)d\Omega + \int_{\Gamma_{\text{outer}}}[\alpha\sigma_{\text{cap,tot}}\mathbf{v}\cdot\mathbf{e}_z + JP_{\text{cyto}}\mathbf{v}\cdot(\mathbf{N}\cdot\mathbf{F}^{-1})]d\Gamma \\ - \int_{\Gamma_{\text{contact}}}\alpha\sigma_{\text{contact}}\mathbf{v}\cdot\mathbf{e}_z d\Gamma - \int_{\Gamma_{\text{inner}}} JP_{\text{nuc}}\mathbf{v}\cdot(\mathbf{N}\cdot\mathbf{F}^{-1})d\Gamma = 0, \tag{S45}$$

where $\mathbf{u},\mathbf{v}$ are trial and test functions for displacement and $p,q$ are trial and test functions for pressure. The pressure is only specified up to an additive constant, and its boundary condition is given as

$$\mathbf{n}\cdot\nabla p = 0 \quad \text{on} \quad \Gamma_{\text{inner}} \text{ and } \Gamma_{\text{outer}}. \tag{S46}$$

This mixed pressure-displacement form is consistent with a compressible formulation in which the bulk modulus is $\phi_0$. We choose $\phi_0$ to be sufficiently large to approximate incompressibility with sufficient accuracy (less than 0.01% deviation in NE volume). With the correct choice of function spaces (linear for pressure, quadratic for displacements), this approach mitigates numerical instabilities when solving incompressible problems via the finite element method [29, 30].

The domain $\Gamma_{\text{contact}}$ evolves over time and/or iteration number during simulations, according to the proximity of a material point to nanopillars. For implementation, it is advantageous to rewrite integrals of any function over this region ($\mathcal{F}$) to avoid discrete updates to the domain at each step,

$$\int_{\Gamma_{\text{contact}}}\mathcal{F}d\Gamma = \int_{\Gamma_{\text{outer}}}\mathcal{F}\mathcal{H}_{\text{NP}}(x,y)d\Gamma, \tag{S47}$$

$$\text{where}\quad \mathcal{H}_{\text{NP}}(x,y) = \sum_{n_x=-N_{\text{NP}}}^{N_{\text{NP}}}\left(\sum_{n_y=-N_{\text{NP}}}^{N_{\text{NP}}}\begin{cases}1 & \text{if } \sqrt{(x-n_x p_{\text{NP}})^2+(y-n_y p_{\text{NP}})^2}\le r_{\text{NP}}\\ 0 & \text{otherwise}\end{cases}\right),$$

where $p_{\text{NP}}$ and $r_{\text{NP}}$ are the pitch and radius of the nanopillars on a given substrate and $N_{\text{NP}} = \text{round}\left(\frac{r_{\max}}{p_{\text{NP}}}\right)$ for $r_{\max}$ setting a maximum length cutoff. Provided the nanopillars do not overlap ($p_{\text{NP}} \ge 2r_{\text{NP}}$), $\mathcal{H}_{\text{NP}}(x,y)$ is equal to one for a region of the NE directly above a nanopillar and is zero otherwise. To avoid sharp cutoffs in our simulation, we define a smooth approximation of $\mathcal{H}_{\text{NP}}$:

$$\mathcal{H}_{\text{NP,smooth}}(x,y) = \sum_{n_x=-N_{\text{NP}}}^{N_{\text{NP}}}\left(\sum_{n_y=-N_{\text{NP}}}^{N_{\text{NP}}}\exp\left(-\frac{((x-n_x p_{\text{NP}})^2+(y-n_y p_{\text{NP}})^2)^2}{(r_{\text{NP}}+0.05)^4}\right)\right), \tag{S48}$$

where super-Gaussian functions are used to approximate the nanopillar domain locations (Figure S1D). Therefore, the fourth term in the above weak form is rewritten as (note there is no dependence on actin or myosin in this term)

$$\int_{\Gamma_{\text{contact}}} J\sigma_{\text{contact}}\mathbf{v}\cdot(\mathbf{N}\cdot\mathbf{F}^{-1})d\Gamma \approx \int_{\Gamma_{\text{outer}}}\mathcal{H}_{\text{NP,smooth}}(x,y)J\sigma_{\text{contact}}\mathbf{v}\cdot(\mathbf{N}\cdot\mathbf{F}^{-1})d\Gamma. \tag{S49}$$

We found that this Robin condition on z-displacement was not sufficient to ensure convergence. Therefore, we supplemented the condition with an additional Dirichlet condition over the uppermost region of the NE at each time step,

$$u_z = \frac{\int_{\Gamma_{\text{upper},2}} \alpha u_{z,\text{prev}} d\Gamma}{\int_{\Gamma_{\text{upper},2}} \alpha d\Gamma} \quad \text{on} \quad \Gamma_{\text{upper},1}, \tag{S50}$$

where $\Gamma_{\text{upper},1}$ contains the surface elements of the outer NE directly connected to the uppermost point of the reference geometry and

$$\Gamma_{\text{upper},2} = \Gamma_{\text{upper}} \backslash \Gamma_{\text{upper},1}, \tag{S51}$$

for $\Gamma_{\text{upper}}$ the region of the upper NE over which $X^2 + Y^2 \leq D^2_{\text{roof}}$ (Figure S1C). $u_{z,\text{prev}}$ is the $z$ displacement at the previous time point.

#### S2.2 Numerical approximation of reaction-transport

To readily facilitate mechanochemical coupling in our equations, we express all of the reaction-transport equations in terms of Lagrangian (material) coordinates. The deformation field $\mathbf{u}$ at the NE is given by the hyperelastic equations above. We extend the displacment field to the nucleoplasm by solving for $\mathbf{u}_{\text{nuc}}$, the displacement associated with a neo-Hookean material within the nucleus. Keeping notation consistent with the equations above, the nucleoplasmic domain $\Omega_{\text{nuc}}$ is the region in which $X^2 + Y^2 + (Z - z_{\text{nuc}} - R_0) \leq R_0 - \Delta T$ (Figure S1B). Defining deformation gradient tensor $\mathbf{F}_{\text{nuc}} = \left(\frac{\partial}{\partial \mathbf{X}}(\mathbf{X} + \mathbf{u}_{\text{nuc}})\right)^T$, right Cauchy-Green tensor $\mathbf{C}_{\text{nuc}} = \mathbf{F}^T_{\text{nuc}} \cdot \mathbf{F}_{\text{nuc}}$, and their invariants ($I_{1,\text{nuc}} = \text{tr}\ \mathbf{C}_{\text{nuc}}, J_{\text{nuc}} = \det\ \mathbf{F}_{\text{nuc}}$), the associated strain energy function is

$$W_{\text{nuc}} = E_{1,\text{nuc}}(I_{1,\text{nuc}} - 3) + \phi_{\text{nuc}}(J_{\text{nuc}} - 1)^2, \tag{S52}$$

where $E_{1,\text{nuc}}$ is the shear modulus and $\phi_{\text{nuc}}$ is the bulk modulus, both chosen according to initial numerical testing (Table S1). Dirichlet boundary conditions are specified by the deformation field at the NE,

$$\mathbf{u}_{\text{nuc}} = \mathbf{u} \quad \text{on} \quad \Gamma_{\text{inner}}. \tag{S53}$$

The associated weak form is

$$\int \nabla \mathbf{v}_{\text{nuc}} \cdot \frac{\partial W_{\text{nuc}}}{\partial \mathbf{F}^T} d\Omega = 0, \tag{S54}$$

where $\mathbf{v}_{\text{nuc}}$ is a test function. As such, we have a global Lagrangian mapping from the reference mesh to the deformed geometry,

$$d\mathbf{x} = d\mathbf{X} \cdot \mathbf{F}^T, \tag{S55}$$

where $\mathbf{F}$ is now defined to extend over the entire nuclear volume; that is,

$$\mathbf{F} = \begin{cases} \left(\frac{\partial}{\partial \mathbf{X}}(\mathbf{X} + \mathbf{u})\right)^T & \text{in } \Omega_{\text{NE}} \\ \left(\frac{\partial}{\partial \mathbf{X}}(\mathbf{X} + \mathbf{u}_{\text{nuc}})\right)^T & \text{in } \Omega_{\text{nuc}} \end{cases}. \tag{S56}$$

We define differential operators in terms of this transformation. Writing $\nabla := \frac{\partial}{\partial \mathbf{x}}$ and $\bar{\nabla} := \frac{\partial}{\partial \mathbf{X}}$,

$$\nabla = \mathbf{F}^{-T} \cdot \bar{\nabla}. \tag{S57}$$

For surface gradients, we must account for changes in the direction of the normal associated with deformations, leading to the relation

$$\nabla_S = \nabla - \mathbf{n}(\mathbf{n} \cdot \nabla) = \mathbf{F}^{-T} \cdot \bar{\nabla} - \left(\frac{J}{\alpha}\right)^2 (\mathbf{N} \cdot \mathbf{F}^{-1}) \left((\mathbf{N} \cdot \mathbf{F}^{-1}) \cdot (\mathbf{F}^{-T} \cdot \bar{\nabla})\right). \tag{S58}$$

Similarly, we can define integrals in terms of Lagrangian coordinates. In particular, the $L^2(O)$-inner product over physical space (given as an integral over the reference domain $\Omega_{\text{full}}$) is

$$\langle a, b\rangle_{\Omega_{\text{full}}} = \int_{\Omega_{\text{full}}} a \cdot b \, J d\Omega. \tag{S59}$$

The $L^2(O)$-inner product written as an integral over reference surface $\Gamma_{\text{outer}}$ is

$$\langle a, b\rangle_{\Gamma_{\text{outer}}} = \int_{\Gamma_{\text{outer}}} a \cdot b \alpha d\Gamma. \tag{S60}$$

We then consider the weak form for a volume species within the nucleoplasm $u$ (either nuclear YAP/TAZ or phosphorylated lamin here),

$$\langle \partial_t u, \phi\rangle_{\Omega_{\text{nuc}}} + \langle D_u \nabla u, \nabla\phi\rangle_{\Omega_{\text{nuc}}} - \langle u\mathbf{v}_{\text{nuc}}, \nabla\phi\rangle_{\Omega_{\text{nuc}}} - \langle f_u, \phi\rangle_{\Omega_{\text{nuc}}} - \langle R_u, \phi\rangle_{\Gamma_{\text{outer}}} = 0, \tag{S61}$$

and for a surface species in the NE, $v$ (dilated or constricted NPCs here),

$$\langle \partial_t v, \psi\rangle_{\Gamma_{\text{outer}}} + \langle D_v \nabla_S v, \nabla_S\psi\rangle_{\Gamma_{\text{outer}}} - \langle v\mathbf{v}_{\text{NE,outer}}, \nabla_S\psi\rangle_{\Gamma_{\text{outer}}} - \langle g_v, \psi\rangle_{\Gamma_{\text{outer}}} = 0, \tag{S62}$$

where $\phi$ and $\psi$ are test functions associated with $u$ and $v$, respectively, $f_u$ is the volumetric reaction rate of $u$, $R_u$ is the surface reaction rate for $u$, and $g_v$ is the surface reaction rate for $v$. These are expressed in a fully Lagrangian fashion by using the above definitions of the inner products (Equations (S59) and (S60)) and differential operators (Equations (S57) and (S58)).

In practice, we solve the system using Lie-Trotter operator splitting, first applying the advection operator. Due to our choice of material coordinates, the effect of advection is to compensate for local changes in volume and surface area over each element (similar to the scheme in [31]),

$$u_A(t) = \frac{J(t-\Delta t)}{J(t)} u(t-\Delta t), \tag{S63}$$

$$v_A(t) = \frac{\alpha(t-\Delta t)}{\alpha(t)} v(t-\Delta t). \tag{S64}$$

We then solve the implicit Euler time-discretized version of the reaction-diffusion system over the fixed domain at time $t$,

$$\frac{1}{\Delta t}\langle u(t) - u_A(t), \phi\rangle_{\Omega_{\text{nuc}}} + \langle D_u \cdot \nabla u(t), \nabla\phi\rangle_{\Omega_{\text{nuc}}} - \langle f_u(t), \phi\rangle_{\Omega_{\text{nuc}}} - \langle R_u(t), \phi\rangle_{\Gamma_{\text{NE}}} = 0, \tag{S65}$$

$$\frac{1}{\Delta t}\langle v(t) - v_A(t), \psi\rangle_{\Gamma_{\text{outer}}} + \langle D_v \nabla_S v(t), \nabla_S\psi\rangle_{\Gamma_{\text{NE}}} - \langle g_v(t), \psi\rangle_{\Gamma_{\text{NE}}} = 0, \tag{S66}$$

where $u_A$ and $v_A$ are used in place of the previous time point values in the time discretization, in line with conventional Lie-Trotter splitting [32, 33].

The reaction terms for each surface and volume species are

$$f_{L_{\text{phos}}} = 0, \tag{S67}$$

$$R_{L_{\text{phos}}} = \frac{1}{N_{\text{conv}}}\left(k_{\text{rl}}[L] - k_{\text{fl}}\frac{p_{\text{cyto}}[F]^{2.6}}{C_L + p_{\text{cyto}}[F]^{2.6}}[L]_{\text{phos}}\right), \tag{S68}$$

$$f_{Y_{\text{nuc}}} = 0, \tag{S69}$$

$$R_{Y_{\text{nuc}}} = \frac{1}{N_{\text{conv}}}\left(k_{\text{in,tot}}[Y]_{\text{free}} - k_{\text{out}}[Y]_{\text{nuc}}\right), \tag{S70}$$

$$g_{NPC_{\text{A}}} = k_{\text{fNPC}}[F][L][M]_A[NPC] - k_{\text{rNPC}}[NPC]_{\text{dilated}}, \tag{S71}$$

$$g_{NPC} = -k_{\text{fNPC}}[F][L][M]_A[NPC] + k_{\text{rNPC}}[NPC]_{\text{dilated}}. \tag{S72}$$

#### S2.3 Mapping species from the inner to outer NE

For lamins present at the inner NE surface, we define an additional mapping. First, we write the natural mapping from the coordinates at the outer NE reference coordinates, $\mathbf{X}_{\text{outer}}$, to the inner NE reference coordinates $\mathbf{X}_{\text{inner}}$:

$$\mathbf{X}_{\text{inner}} = \mathbf{X}_{\text{center}} + (\mathbf{X}_{\text{outer}} - \mathbf{X}_{\text{center}}) \left( \frac{R_0 - \Delta T}{R_0} \right), \tag{S73}$$

where $\mathbf{X}_{\text{center}} = [0, 0, z_{\text{nuc}} + R_0]$ (see Equations (S1) and (S2)). Then, we can express the displacement of the deformed inner surface with respect to $\mathbf{X}_{\text{outer}}$:

$$\mathbf{u}_{\text{inner}} := \mathbf{x}_{\text{inner}} - \mathbf{X}_{\text{outer}} = \mathbf{u}[\mathbf{X}_{\text{inner}}] + \mathbf{X}_{\text{inner}} - \mathbf{X}_{\text{outer}}, \tag{S74}$$

where square brackets ($[\cdot]$) are used to indicate function evaluation at the indicated coordinates. The associated deformation gradient tensor $\mathbf{F}_{\text{alt}}$ is then defined as

$$\mathbf{F}_{\text{alt}} := \frac{\partial}{\partial \mathbf{X}_{\text{outer}}} (\mathbf{X}_{\text{outer}} + \mathbf{u}_{\text{inner}}) = \frac{\partial \mathbf{X}_{\text{inner}}}{\partial \mathbf{X}_{\text{outer}}} \mathbf{F}[\mathbf{X}_{\text{inner}}] = \left( \frac{R_0 - \Delta T}{R_0} \right) \mathbf{F}[\mathbf{X}_{\text{inner}}], \tag{S75}$$

The associated mapping is then

$$d\mathbf{x}_{\text{inner}} = d\mathbf{X}_{\text{outer}} \cdot \mathbf{F}_{\text{alt}}^T. \tag{S76}$$

Using the following relations (analogous to Equations (S26) and (S27)),

$$\alpha_{\text{inner}} = |\det(\mathbf{F}_{\text{alt}}) \mathbf{N} \cdot \mathbf{F}_{\text{alt}}^{-1}|, \tag{S77}$$

$$\mathbf{n}_{\text{inner}} = \frac{\det(\mathbf{F}_{\text{alt}}) \mathbf{N} \cdot \mathbf{F}_{\text{alt}}^{-1}}{\alpha_{\text{inner}}}, \tag{S78}$$

the resulting definition of the surface gradient is

$$\nabla_{\text{S,inner}} = \nabla - \mathbf{n}_{\text{inner}} (\mathbf{n}_{\text{inner}} \cdot \nabla) = \mathbf{F}_{\text{alt}}^{-T} \cdot \bar{\nabla} - \left( \frac{\det \mathbf{F}_{\text{alt}}}{\alpha_{\text{inner}}} \right)^2 (\mathbf{N} \cdot \mathbf{F}_{\text{alt}}^{-1}) \left( (\mathbf{N} \cdot \mathbf{F}_{\text{alt}}^{-1}) \cdot (\mathbf{F}_{\text{alt}}^{-T} \cdot \bar{\nabla}) \right), \tag{S79}$$

and the $L^2(O)$-inner product written as an integral over the reference surface $\Gamma_{\text{outer}}$ is

$$\langle a, b \rangle_{\Gamma_{\text{inner}}} = \int_{\Gamma_{\text{outer}}} a \cdot b \alpha_{\text{inner}} d\Gamma. \tag{S80}$$

Given these definitions, we give the weak form for dephosphorylated lamins,

$$\langle \partial_t [L], \chi \rangle_{\Gamma_{\text{inner}}} + \langle D_L \nabla_{\text{S,inner}} [L], \nabla_{\text{S,inner}} \chi \rangle_{\Gamma_{\text{inner}}} - \langle [L] \mathbf{v}_{\text{NE,inner}}, \nabla_{\text{S,inner}} \chi \rangle_{\Gamma_{\text{inner}}} - \langle g_L, \chi \rangle_{\Gamma_{\text{inner}}} = 0, \tag{S81}$$

where

$$g_L = -k_{\text{rl}} [L] + k_{\text{fl}} \frac{p_{\text{cyto}} [F]^{2.6}}{C_L + p_{\text{cyto}} [F]^{2.6}} [L]_{\text{phos}}, \tag{S82}$$

and $\chi$ is a test function. In keeping with the above splitting of diffusion and advection, we solve this in two steps:

$$[L]_A(t) = \frac{\alpha_{\text{inner}}(t - \Delta t)}{\alpha_{\text{inner}}(t)} [L](t - \Delta t). \tag{S83}$$

$$\frac{1}{\Delta t} \langle [L](t) - [L]_A(t), \chi \rangle_{\Gamma_{\text{inner}}} + \langle D_L \nabla_{\text{S,inner}} [L](t), \nabla_{\text{S,inner}} \chi \rangle_{\Gamma_{\text{inner}}} - \langle g_L(t), \chi \rangle_{\Gamma_{\text{inner}}} = 0, \tag{S84}$$

#### S2.4 Calculation of cytosolic variables

We assume that cytosolic YAP/TAZ is well-mixed, resulting in a single concentration of free YAP/TAZ at each time. We furthermore assume that the sequestration/de-sequestration of YAP/TAZ in the cytosol is at pseudo-steady state; that is:

$$(k_{\mathrm{CN}} + k_{\mathrm{CY}}[F]_0[M]_A)[Y]_{\mathrm{seq}} - k_{\mathrm{NC}}[Y]_{\mathrm{free}} = 0, \tag{S85}$$

where $k_{\mathrm{CN}}$ and $k_{\mathrm{CY}}$ are the cytoskeleton-independent and -dependent de-sequestration rates, respectively. Note that the global actin concentration $[F]_0$ is used here, as the concentration adopted in the majority of the cytosol. Assuming a conserved number of YAP/TAZ molecules over time, $N_Y$ and a conserved total cell volume $\mathrm{vol}_{\mathrm{tot}}$, we can compute the total cytosolic concentration as:

$$[Y]_{\mathrm{cyto}} = \frac{N_Y - N_{\mathrm{conv}} \int_{\Omega_{\mathrm{nuc}}} [Y]_{\mathrm{nuc}} d\Omega}{N_{\mathrm{conv}}(\mathrm{vol}_{\mathrm{tot}} - \int_{\Omega_{\mathrm{nuc}}} d\Omega)}, \tag{S86}$$

where $N_{\mathrm{conv}}$ is a conversion factor, 602.2 (molecules/$\mu$m$^3$)/$\mu$M. Observing that $[Y]_{\mathrm{cyto}} = [Y]_{\mathrm{free}} + [Y]_{\mathrm{seq}}$, we can then write $[Y]_{\mathrm{free}}$ at any time point:

$$[Y]_{\mathrm{free}} = [Y]_{\mathrm{cyto}} \frac{k_{\mathrm{CN}} + k_{\mathrm{CY}}[F]_0[M]_A}{k_{\mathrm{NC}} + k_{\mathrm{CN}} + k_{\mathrm{CY}}[F]_0[M]_A}. \tag{S87}$$

Cytosolic F-actin is computed from Equation (S34), which requires estimates of the distance between the PM and the NE and the local curvature of the PM. We assume a steric radius of 0.05 $\mu$m between the nanopillar and PM and consider only the closest nanopillar (centered at $(x_{\mathrm{NP}}, y_{\mathrm{NP}})$) to a given point on the NE. Assuming a cylindrical PM geometry, the distance to the PM at the side and the top of the nanopillar are then given by

$$d_{\mathrm{top}} = \begin{cases} z + d_{\mathrm{steric}} & \text{if } r_{\mathrm{local}} \leq r_{\mathrm{NP}} + 0.05 \\ \sqrt{(r_{\mathrm{local}} - r_{\mathrm{NP}})^2 + (z + d_{\mathrm{steric}})^2} & \text{otherwise} \end{cases}, \tag{S88}$$

$$d_{\mathrm{side}} = \begin{cases} \sqrt{(r_{\mathrm{local}} - r_{\mathrm{NP}})^2 + (z + d_{\mathrm{steric}})^2} & \text{if } z > -d_{\mathrm{steric}} \\ r_{\mathrm{local}} - r_{\mathrm{NP}} & \text{otherwise} \end{cases}, \tag{S89}$$

where $r_{\mathrm{local}} = \sqrt{(x - x_{\mathrm{NP}})^2 + (y - y_{\mathrm{NP}})^2}$ and $d_{\mathrm{steric}}$ is a gap due to steric repulsion between the PM and NE. According to the assumed geometry, PM curvature at the top is identically zero and curvature at the side is $\frac{1}{2(r_{\mathrm{NP}}+0.05)}$. Given these definitions, we solve for F-actin using Equation (S34) as follows:

$$[F] = \begin{cases} [F]_0 + [F]_1 \exp\left(-\frac{z + d_{\mathrm{steric}}}{d_F}\right) & \text{if } h_{\mathrm{NP}} = 0 \\ [F]_0 + [F]_1 \max\left[\exp\left(-\frac{d_{\mathrm{top}}}{d_F}\right), \exp\left(-\frac{d_{\mathrm{side}}}{d_F}\right) \exp\left(\frac{1}{2(r_{\mathrm{NP}}+0.05)H_0}\right)\right] & \text{otherwise} \end{cases}, \tag{S90}$$

where $h_{\mathrm{NP}} = 0$ corresponds to a flat substrate. The use of the maximum in the second case guarantees a smooth transition in the F-actin concentration from the side to the top region of the NE near the nanopillar.

#### S2.5 Software implementation

In brief, solving the coupled mechanochemical system involves the following steps:

1. Compute resting NE displacement ($\mathbf{u}$) after applying osmotic pressure gradient $\Delta P_{\mathrm{max}}$ with $z$-axis symmetry condition Equation (S33), no cap stress, and no nanopillar contact stresses.
2. Initialize reaction-transport system in SMART over reference mesh with initial deformation $\mathbf{u}$.
3. Update nuclear deformation according to stiffness governed by initial lamin content (Equation (S43)).

4. Compute $[F]$ (Equation (S90)), $\alpha_{\text{init}}$ (Equation (S26)), and $[Y]_{\text{free}}$ (Equation (S87)) over deformed coordinates.
5. Update cap stress to $\sigma_{\text{cap}}(t+\Delta t)$ and solve for new nuclear displacement (Equation (S45)). Time step is restricted such that the maximum increment in applied stress is 2 kPa.
6. Update volume estimate using Anderson acceleration with $m = 3$, then update pressures $P_{\text{nuc}}$ and $P_{\text{cyto}}$ according to Equations (S29) and (S30) and solve for updated displacements. Repeat this process until the change in volume is less than 0.1%.
7. Update $[F]$ (Equation (S34)) and $\alpha$ (Equation (S26)).
8. Account for changes in local volume due to advection (Equations (S63), (S64) and (S83)).
9. Solve for updated concentrations according to reaction-diffusion (Equations (S65), (S66) and (S84)).
10. Update $[Y]_{\text{free}}$ (Equation (S87)) and NE stiffness (Equation (S43)).
11. Repeat steps 5-10 until $t > 10000$ s.

##### S2.6 Estimation of NER likelihood

To determine the likelihood of rupture, we used Bell's law, calculating an effective rupture rate for an attachment between lamin dimers that compose the filaments,

$$k_{\text{rupture}}(t) = k_{\text{rupture},0} \exp\left(\frac{\delta_L \frac{F_L(t)}{\phi_{\text{layer}}}}{k_B T}\right), \tag{S91}$$

where $\delta_L$ is the molecular displacement associated with rupture of a single lamin dimer, $k_B T$ is the thermal energy scale, and $k_{\text{rupture},0}$ is the rupture rate when no force is applied. $F_L$ is defined as the magnitude of the force defined in Equation (S44), based on evaluating the stress acting on a single representative cross section of the NE-lamin composite shell. We account for the fact that not all lamins composing the lamina pass through this cross section by dividing by a geometric scaling factor, $\phi_{\text{layer}}$, which is expected to scale as the thickness of a single dimer divided by the total thickness of the lamina (about 0.1). Practically speaking, we consider a lumped parameter $\hat{\delta} = \frac{\delta}{\phi_{\text{layer}}}$ below. Calculations of the per-molecule force in future iterations can include additional details of the relationship between the bulk mechanics and lamin microstructure, as in [34].

The likelihood of rupture for a single lamin subunit over a time interval $[t, t+\Delta t]$ is then

$$P_{\text{rupture}} = 1 - \exp\left(-\int_t^{t+\Delta t} k_{\text{rupture}} d\tau\right). \tag{S92}$$

Treating rupture at each dimer-dimer interface as independent events, the likelihood of at least one rupture event occurring over a patch with area $A_{\text{patch}}$ and average lamin density $[L]$ is

$$P_{\text{rupture,patch}} = 1 - \left[\exp\left(-\int_t^{t+\Delta t} k_{\text{rupture}} d\tau\right)\right]^{A_{\text{patch}}[L]}. \tag{S93}$$

The cumulative rupture probability over the entire nuclear envelope at time $t$ is therefore given by

$$P_{\text{rupture,tot}}(t) = 1 - \prod_{i=1}^{N_{\text{el}}} \left(\left[\exp\left(-\int_0^t k_{\text{rupture,i}} d\tau\right)\right]^{A_{\text{patch,i}}[L]_i}\right), \tag{S94}$$

where the product is evaluated over all $N_{\text{el}}$ faces of the NE surface mesh, with each element $i$ associated with area $N_{\text{el,i}}$, lamin density $[L]_i$, and rupture rate $k_{\text{rupture,i}}$. The latter two quantities were evaluated at the centroid of the element under consideration.

This leaves two new parameters, $k_{\text{rupture},0}$ and $\hat{\delta}_L$. We varied $\hat{\delta}_L$ from 5-75 nm according to characteristic lamin deformations measured in[23] and the expected value of $\phi_{\text{layer}}$ above ($\approx 0.1$). At each value of $\hat{\delta}$, we set $k_{\text{rupture},0}$

such that the rupture probability for baseline (100% lamin expression) cells on 3 μm pitch, 200 nm radius nanopillars was 80%, in line with measurements on similar nanopillar substrates in previous work (radius 250 nm, spacing 3.5 microns) [35]. The associated value for $k_{\text{rupture},0}$ was determined iteratively using the secant method, with the results summarized in Table S3.

**Table S3:** Determination of $k_{\text{rupture}}$ for different choices of $\hat{\delta}$ on a nanopillar substrate with $r_{\text{NP}} = 200$ nm, $p_{\text{NP}} = 3.0$ μm, and $h_{\text{NP}} = 2.5$ μm.

| $\hat{\delta}$ (nm) | $k_{\text{rupture},0}$ (1/s) | Rupture likelihood |
|---|---|---|
| 5.0 | $2.212 \times 10^{-10}$ | 0.8000 |
| 10.0 | $7.646 \times 10^{-11}$ | 0.7998 |
| 15.0 | $2.559 \times 10^{-11}$ | 0.7997 |
| 20.0 | $8.152 \times 10^{-12}$ | 0.7996 |
| 25.0 | $2.417 \times 10^{-12}$ | 0.7995 |
| 30.0 | $6.509 \times 10^{-13}$ | 0.7994 |
| 35.0 | $1.568 \times 10^{-13}$ | 0.7996 |
| 40.0 | $3.383 \times 10^{-14}$ | 0.7999 |
| 45.0 | $6.540 \times 10^{-15}$ | 0.7943 |
| 50.0 | $1.234 \times 10^{-15}$ | 0.8024 |
| 55.0 | $2.154 \times 10^{-16}$ | 0.7998 |
| 60.0 | $3.678 \times 10^{-17}$ | 0.7998 |
| 65.0 | $6.151 \times 10^{-18}$ | 0.7997 |
| 70.0 | $1.014 \times 10^{-18}$ | 0.7997 |
| 75.0 | $1.651 \times 10^{-19}$ | 0.7997 |

We then tested this set of values for $\hat{\delta}$ and $k_{\text{rupture},0}$ for cells on a flat substrate. At low values of $\hat{\delta}$, rupture likelihood was still very high on these surfaces; we chose the value of $\hat{\delta}$ that resulted in only a 1% rupture likelihood on flat surfaces ($\hat{\delta} = 65$ nm, as shown in Figure S11A) The constrained value for the rupture rate constant was very small ($k_{\text{rupture}} = 6.151 \times 10^{-18}$ s$^{-1}$), indicating that lamin rupture should almost never occur in the absence of applied force.

### S3 Comparison with our previous model

To inform the relationship between cytoskeletal assembly, nucleoskeletal assembly, and YAP transport in our mechanochemical model, we ran simulations on whole cell geometries using our previous formulation [14]. We constructed cell meshes directly informed by the cell and nuclear volumes measured in experiments (Figure S7A). Nuclear volumes were chosen as slightly smaller than average measurements to ensure that the nucleus fit cleanly within the cell boundary without any mesh intersections. Furthermore, the cell-substrate contact areas were chosen to match experimentally measured projected cell areas on each substrate (Figure S7A). To better represent the variability in cell-substrate contact in these simulations, we constructed meshes from the average contact area ± 2 × the standard error of the mean (approximately the 95% confidence interval for each distribution).

Given these geometries, we ran simulations using our previous model equations with two important modifications:

- We assumed a much higher maximum density of bound integrins ($\rho_{\text{I,max}}$ in equation 1 from [14] was set to 10× its previous value), which was needed to recapitulate the nuclear localization of YAP/TAZ despite smaller cell-substrate contact areas than previously tested. This choice is consistent with a higher ligand density on our substrates, as this can lead to restricted cell spreading [36].
- We assumed the plasma membrane did not form robust contacts with the lower surface on tall (2.5 μm height) nanopillars, as shown in Figure S7. Assuming conserved total cell volume, it is difficult for the cell to fully spread downward between pillars; indeed, previous images show that cells often sit on top of larger nanopillar arrays, especially on lower-pitch arrangements [37]. However, some small contacts may still form (e.g., via filopodial extensions [38, 39]), so we include an adhesive signal along the lower cell surface that scales with the macroscopic gap between the lower substrate and the plasma membrane,

$$\rho_{\text{I,max}} = \rho_{\text{I,max},0} \exp\left(-\frac{d_{\text{gap}}}{1\ \mu\text{m}}\right) \tag{S95}$$

Altogether, this treatment resulted in strong agreement with our experimental measurements of YAP nuclear-to-cytosolic ratio (N/C) (Figure S7C).

### S4 Experimental Methods

#### S4.1 Nanopillar substrate fabrication and functionalization

Quartz nanopillar arrays were fabricated using projection photolithography and reactive ion etching (RIE) as previously described [40, 35, 14]. Briefly, we patterned 4-inch fused quartz wafers (Wafer Pro) using a Heidelberg MLA system and a chromium hard mask. The pattern was transferred onto the quartz substrate via REI using Ar/CHF3 plasma (Oxford Plasmalab 80), followed by wet etching to remove the mask and residual defects. The resulting nanopillars used for the experiments in Fig 5 had a height of $3.24 \pm 0.25$ µm, diameter of $0.90 \pm 0.016$ µm, and a center-to-center pitch of $3.49 \pm 0.06$ µm, as quantified via scanning electron microscopy as in previous work [41]. The fabricated nanopillars used in all other experiments (Figs 2-4, Figures S6 and S7) had a height of $2.98 \pm .16$ µm ($N$=100), a diameter of $0.703 \pm 0.046$ µm ($N$=38), and center-to-center pitches of $3.09 \pm 0.05$ µm and $6.05 \pm 0.13$ µm ($N$=30). The dimensions of these nanopillar arrays were also measured via SEM, as shown in Figure S4.

Prior to seeding cells, nanopillar chips were sterilized and then washed with 70% ethanol and dionized (DI) water, or sonicated in isopropyl alcohol (IPA). After being air-dried, the chips underwent UV-ozone treatment for 10-20 min. To promote cell adhesion, the chips were functionalized with a sequential coating of poly-L-lysine (PLL, Sigma-Aldrich, USA), glutaraldehyde (Thermo Scientific Chemicals, Thermo Fisher Scientific, USA), and gelatin (Sigma-Aldrich, USA). For the experiments in Fig 5, chips were incubated with 100 µg/mL PLL for 30 min at room temperature (RT), 0.5% glutaraldehyde (Thermo Scientific Chemicals, Thermo Fisher Scientific, USA) for 30 min (RT), and 0.1 wt% gelatin (Sigma-Aldrich, USA) for 30 min at 37°C. For all other experiments (Figs 2-4, Figures S6 and S7), 1% glutaraldehyde was used for 5 min (RT), and 1 wt% gelatin was applied and stored overnight at 4°C, then crosslinked at 37°C for 30 minutes before seeding.

#### S4.2 Cell culture and seeding

U2OS cells (ATCC, USA) were maintained in McCoy's 5A (modified) Medium (ATCC, USA) supplemented with 10% fetal bovine serum (FBS, Sigma-Aldrich, USA) and 1% penicillin-streptomycin (P/S, Sigma-Aldrich, USA) at 37°C and 5% $CO_2$. For experiments, cells were detached using TrypLE Express Enzyme (1X) (Gibco, USA) and resuspended to a density of $5 \times 10^5$ cells/mL. A 100 µL droplet of the cell suspension (∼50,000 cells) was dispensed onto the functionalized nanopillar chips in a 24-well plate. Cells were allowed to adhere for 10 min before the addition of 1 mL of growth media.

#### S4.3 Lamin A/C silencing

To induce lamin A/C depletion, U2OS cells were transfected with lamin A/C siRNA (sc-35776, Santa Cruz Biotechnology, USA) using the siRNA Transfection Reagent (sc-29528) and Transfection Medium (sc-36868) according to the manufacturer's protocol. The siRNA stock was reconstituted in RNase-free water to a concentration of 10 µM prior to transfection. Knockdown efficiency was subsequently confirmed via immunofluorescence staining using lamin A/C antibodies.

#### S4.4 Immunofluorescence Staining

Cells were fixed with 4% paraformaldehyde (PFA, Electron Microscopy Sciences, USA) for 10 min at room temperature, washed with PBS to remove excess PFA, and then permeabilized with 1% Triton X-100 (Sigma-Aldrich, USA) for 10 min. Samples were then blocked with 2% w/v Bovine Serum Albumin (BSA, Thermo Scientific, USA) for 1 h at RT, or with 1% w/v BSA overnight at 4°C. For primary antibody labeling in lamin experiments, samples were incubated with mouse anti-Lamin A/C (1:400, Biolegend; Cat: 600002) in 1% BSA overnight at 4°C. For primary antibody labeling in YAP experiments, samples were incubated with rabbit anti-YAP (1:400, Invitrogen, Thermo Fisher Scientific, USA; Cat: PA1-46189). For Ku-80 immunolabeling, samples were incubated with rabbit anti-Ku-80 (1:400, Cell Signaling Technology, USA; Cat: 4412S) concurrently with anti-lamin A/C in lamin experiments. In YAP experiments, samples were incubated with mouse anti-Ku-80 (1:400, Thermo Fisher Scientific, USA; Cat: MA5-15873) concurrently with anti-YAP. Following PBS washes, secondary antibodies conjugated to anti-mouse

Alexa Fluor 647 (Thermo Fisher Scientific, USA; Cat: A-31571) and anti-rabbit Alexa Fluor 488 (Cell Signaling, USA; Cat: 4412) were applied at a 1:400 dilution in 1% w/v BSA for 1 h at RT. Following antibody incubation, samples were washed with PBS. Nuclei were stained with 4',6-diamidino-2-phenylindole (DAPI, Thermo Scientific, USA; dilution 300 nM) for 5 min.

#### S4.5 Microscopy

Fluorescence images were acquired using an Echo Revolution microscope equipped with a 60x PLAN Fluorite Water Dipping objective (NA = 1.00). Confocal images were acquired using an Eclipse Ti2-E microscope (Nikon, Japan) equipped with a CSU-X1 spinning disk confocal unit (Yokogawa Electric Corporation, Japan), a 60x Plan Apo λ oil-immersion objective (NA = 1.40), and a 100x SR HP APO TIRF oil-immersion objective (NA = 1.49). Glass-bottomed dishes with #1.5 cover glass (Cellvis, USA; Cat: D35-20-1.5-N) were used and filled with 1 mL PBS. We placed NP chips in the dishes with the pillars facing downward before imaging with a step size of 200 nm.

#### S4.6 Image Analysis

Image analysis was performed using ImageJ 1.53 (NIH, USA). For quantifying indentation and lamin intensity profiles from confocal images, images were pre-processed by subtracting the average background intensity. Each image was straightened, and an orthogonal projection was generated capturing NPs along the major axis of the cell. A curve was drawn manually along the lower NE, and an ImageJ Macro was run to extract the spatial location and gray value for each point along the curve. Then, a MATLAB code was run to find peaks and troughs in spatial coordinates, quantifying NP-induced indentations. Once NP locations were found, MATLAB was used to extract the gray values along the lower NE, and the central NP was used as the origin point. If there were an even number of total NPs, producing two central NPs, the rightmost central NP was chosen as the origin. For quantifying YAP N/C from confocal images, images were pre-processed by subtracting the average background intensity. Image channels were split, and the DAPI channel was thresholded to generate a nuclear mask, while the YAP channel was thresholded to generate a cytosolic mask, both using Otsu's method. An ImageJ Macro was used to iterate through each slice, apply the masks, and quantify the integrated nuclear and cytosolic YAP intensities for each slice. The integrated intensities were summed across all slices and divided by the nuclear vs. cytoplasmic volume, as determined by the voxel count, to determine average nuclear YAP and average cytosolic YAP, which were divided to obtain the YAP N/C ratio. To quantify the flattening ratio, the effective maximum diameter of the nucleus was calculated from the maximum area projection and the nuclear height. MATLAB code and ImageJ macros can be found in Data Availability.

For quantifying NER in low Lamin A/C cells, images were pre-processed prior to segmentation using a rolling ball background subtraction (radius: 50 pixels, sliding parabolic enabled) and a Gaussian blur ($\sigma = 2.0$) to reduce noise. Nuclear boundaries were segmented by thresholding the DAPI channel (Otsu's method). To define the whole-cell boundary, the transmitted light channel was used to outline the cell edge. The mean fluorescence intensity (MFI) of lamin A/C was measured within the defined nuclear boundary. For Ku-80, the cytoplasmic-to-nuclear ratio (C/N) was determined via line profile analysis. A linear region of interest (ROI) was drawn across the nucleus extending into the cytoplasm. Intensity profiles were generated to capture both the nuclear signal and the perinuclear cytoplasmic plateau. The C/N was calculated by dividing the mean cytoplasmic intensity by the mean nuclear intensity after normalizing for background signal.

#### S4.7 Rupture Classification

NER was classified based on the Ku-80 C/N. To objectively determine the threshold separating ruptured from non-ruptured cells, a Gaussian Mixture Model (GMM) was applied to the population distribution of Ku-80 C/N. The GMM analysis identified two distinct clusters corresponding to non-ruptured and ruptured phenotypes. The intersection point of the fitted Gaussian distributions provided a data-driven threshold of 0.16. Cells with a Ku-80 C/N exceeding this value were classified as ruptured. For experiments on populations of nonruptured cells, ruptured cells were manually excluded from the dataset through visual identification during confocal imaging.

#### S4.8 Statistical Analysis

All experimental results were taken from at least three independent experiments. Statistical comparisons shown in Fig 2, Fig 4, and Figures S6 and S7 were conducted using the 'scipy' and 'statsmodels' modules in Python. As detailed in each associated caption, comparisons between only two experimental groups were made using an unpaired Student's $t$-test, whereas comparisons between 3 or more groups were made by first conducting one-way or two-way analysis of variance (ANOVA), followed by Tukey's posthoc test for multiple comparisons. Multiple comparisons were only assessed when ANOVA revealed a significant difference between groups. Pearson's correlation coefficient and the associated $p$-value were computed for the cases shown in Fig 3 using 'scipy'. The statistical comparison shown in Fig 5 was performed using an unpaired Student's $t$-test in GraphPad Prism software.

### S5 Supplementary Figures

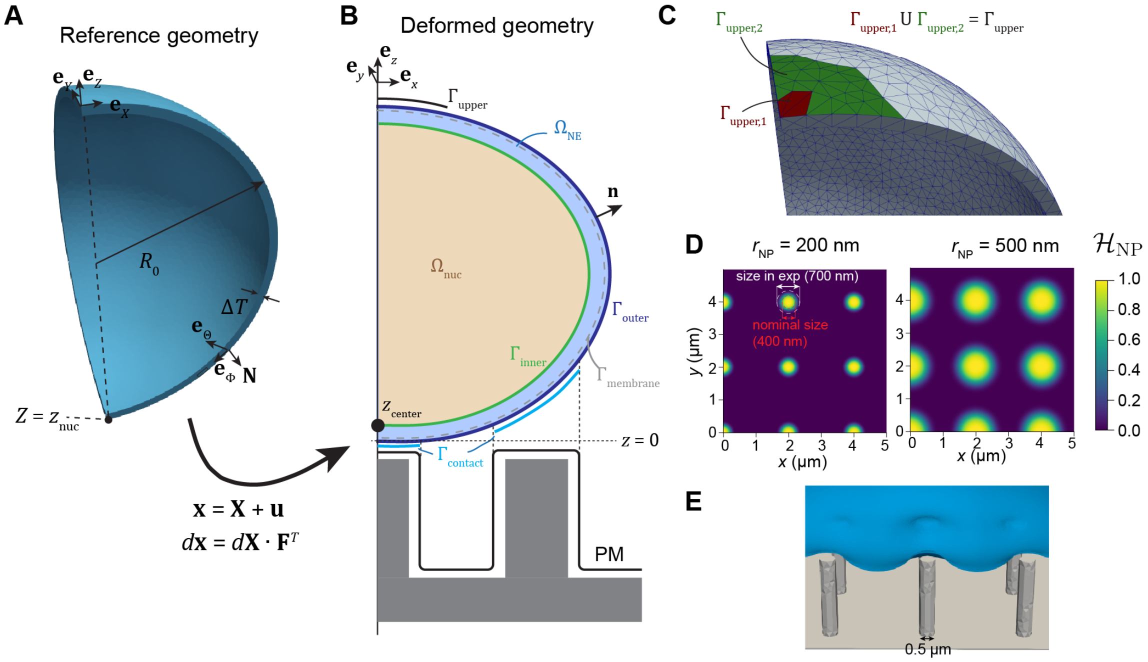


**Figure S1: Coordinate and domain definitions for mechanical model.** A) Reference geometry with unit vectors denoting Cartesian coordinates and spherical coordinates. B) Schematic showing volumetric domains ($\Omega_{nuc}$ and $\Omega_{NE}$) and surface domains ($\Gamma_{inner}$ and $\Gamma_{outer}$) of the nucleus. The outer NE includes subdomains $\Gamma_{upper}$ and $\Gamma_{contact}$ as indicated. $\Gamma_{membrane}$ denotes the midplane of the nuclear membranes and perinuclear space, assumed to occupy a thickness of 50 nm in the resting configuration [10]. PM surface included for illustrative purposes. C) Depiction of upper NE subdomains included in Equation (S50) and Equation (S51). D) Smooth approximation of $\Gamma_{contact}$ given by Equation (S48) for $r_{NP}$ = 200 or 500 nm, $p_{NP}$ = 2 μm. Outlines show the prescribed radius versus the size measured in experiments. E) Example NE morphology draped over a 200 nm radius nanopillar (with depiction including steric radius of 50 nm).

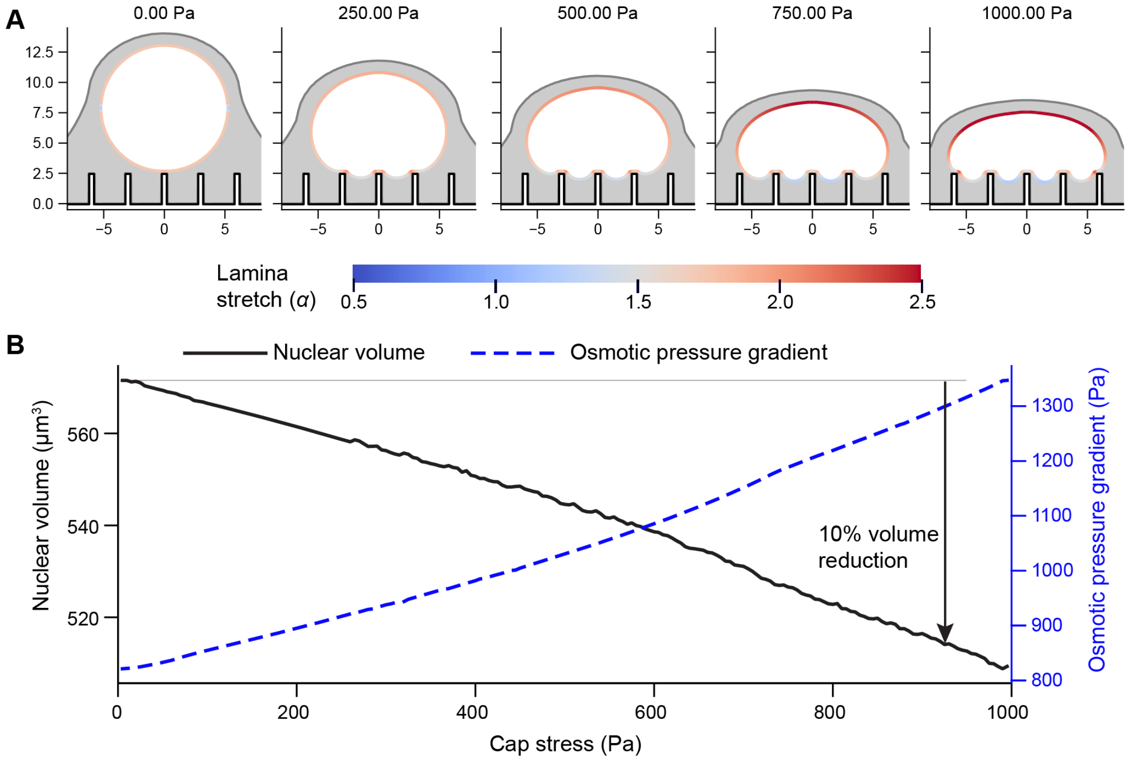


**Figure S2: Evolution of pressure and volume during nuclear flattening on 3 micron pitch nanopillars** A) Nuclear deformation over time on a 3 μm pitch nanopillar substrate, with the color map indicating nuclear lamina stretch. B) Decreases in the nuclear volume are associated with changes in the osmotic pressure gradient across the NE, dictated by Equations (S29) and (S30).

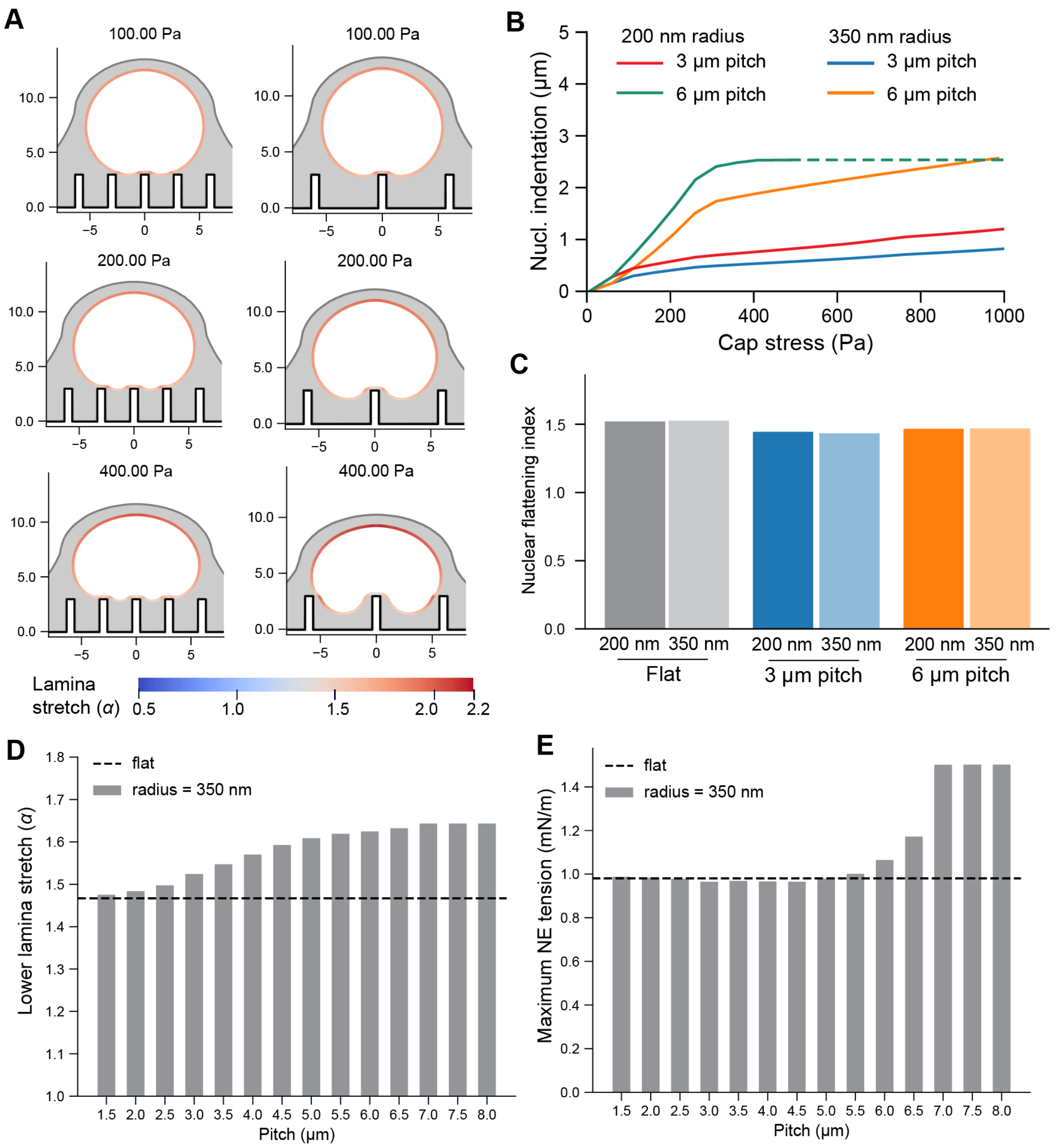


**Figure S3: Nuclear deformation on larger radius nanopillars.** A) Predicted nuclear geometries for deformation on 3 μm (left) and 6 μm (right) pitch substrates with $r_{\mathrm{NP}} = 350$ nm and $h_{\mathrm{NP}} = 3.0$ μm. Color depicts stretch ($\alpha$) of the laminar surface at the specified cap stress. B-C) Comparison of NE indentation (B) and the nuclear flattening index (C) on the 200 nm radius nanopillars testing throughout versus the 350 nm radius pillars here. D-E)

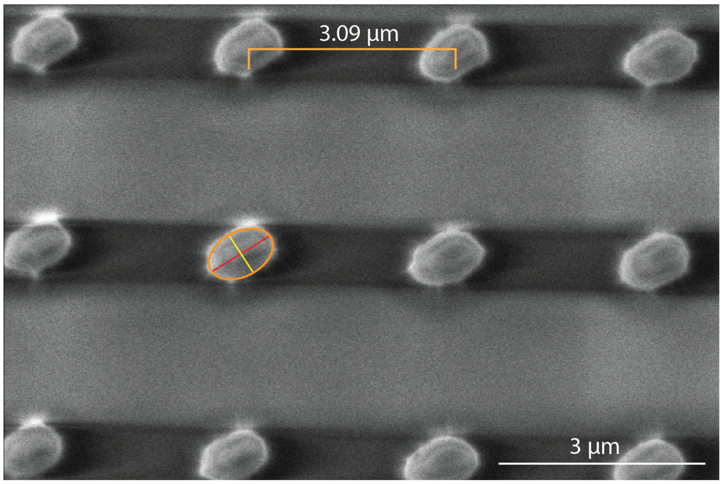


**Figure S4:** Representative SEM image of the nanopillar array. Nanopillar effective diameter was calculated as the mean of the major and minor axes of the elliptical nanopillar profile (red and yellow lines, respectively), as illustrated. The nanopillars had an effective diameter of 0.703 ± 0.046 µm and a center-to-center pitch of 3.09 ± 0.05 µm. Scale bar, 3 µm.

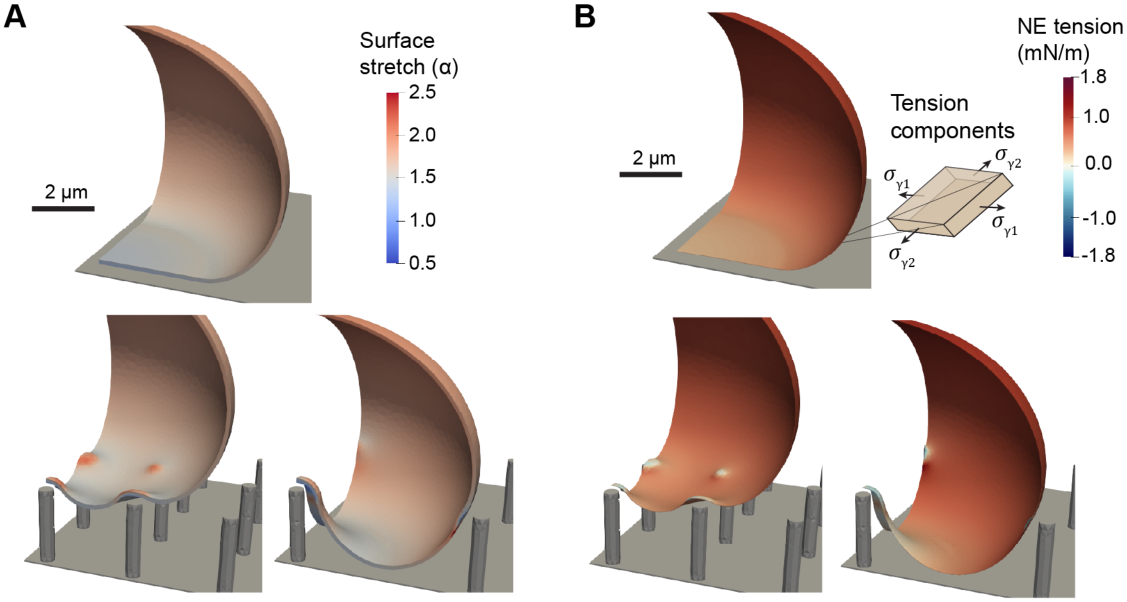


**Figure S5: NE stretch and tension are maximized on moderate to high-pitch nanopillar substrates.** A) The stretch ratio of the nuclear surface ($\alpha$) is shown for different nanopillar pitches with a cap stress of 400 Pa. B) The prediction NE tension is shown for different nanopillar pitches with a cap stress of 400 Pa.

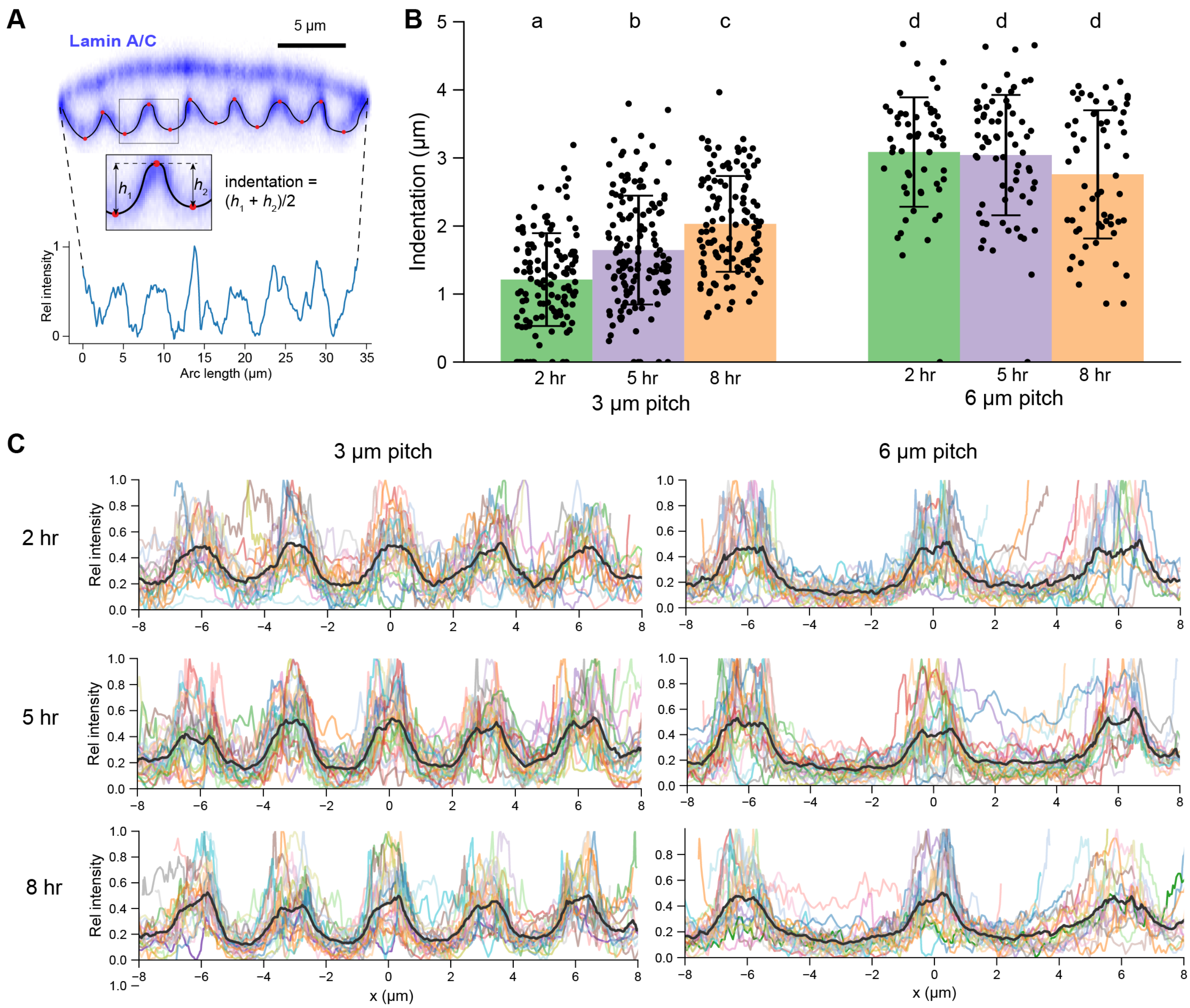


**Figure S6: NE indentation and lamin localization at different timepoints.** A) Sample lamin cross section from confocal microscopy used to quantify indentation and lamin density. The black line indicates the path over which the lamin density was quantified from the image and the red points indicate the peaks and troughs used to measure indentation. B) NE indentation as a function of time and pitch. On 3 μm pitch substrates, we measured $N = 24$ cells, 124 indentations at 2 hr, $N = 27$ cells, 141 indentations at 5 hr, and $N = 24$ cells, 124 indentations at 8 hr. On 6 μm pitch substrates, we measured $N = 24$ cells, 56 indentations at 2 hr, $N = 27$ cells, 68 indentations at 5 hr, and $N = 24$ cells, 60 indentations at 8 hr. Statistical significance was determined by using 2-way ANOVA followed by Tukey's post-hoc test, letters above each group denote statistical significance, where groups not sharing a letter are statistically significantly different. C) Average (black lines) and cell-by-cell (colored lines) lamin contours as a function of time point and pitch (24 profiles in each 2 hr and 8 hr case, 27 profiles in each 5 hr case). The $y$-axis provides the rescaled lamin intensity, where the scale is set from 0 to 1 on a cell-by-cell basis.

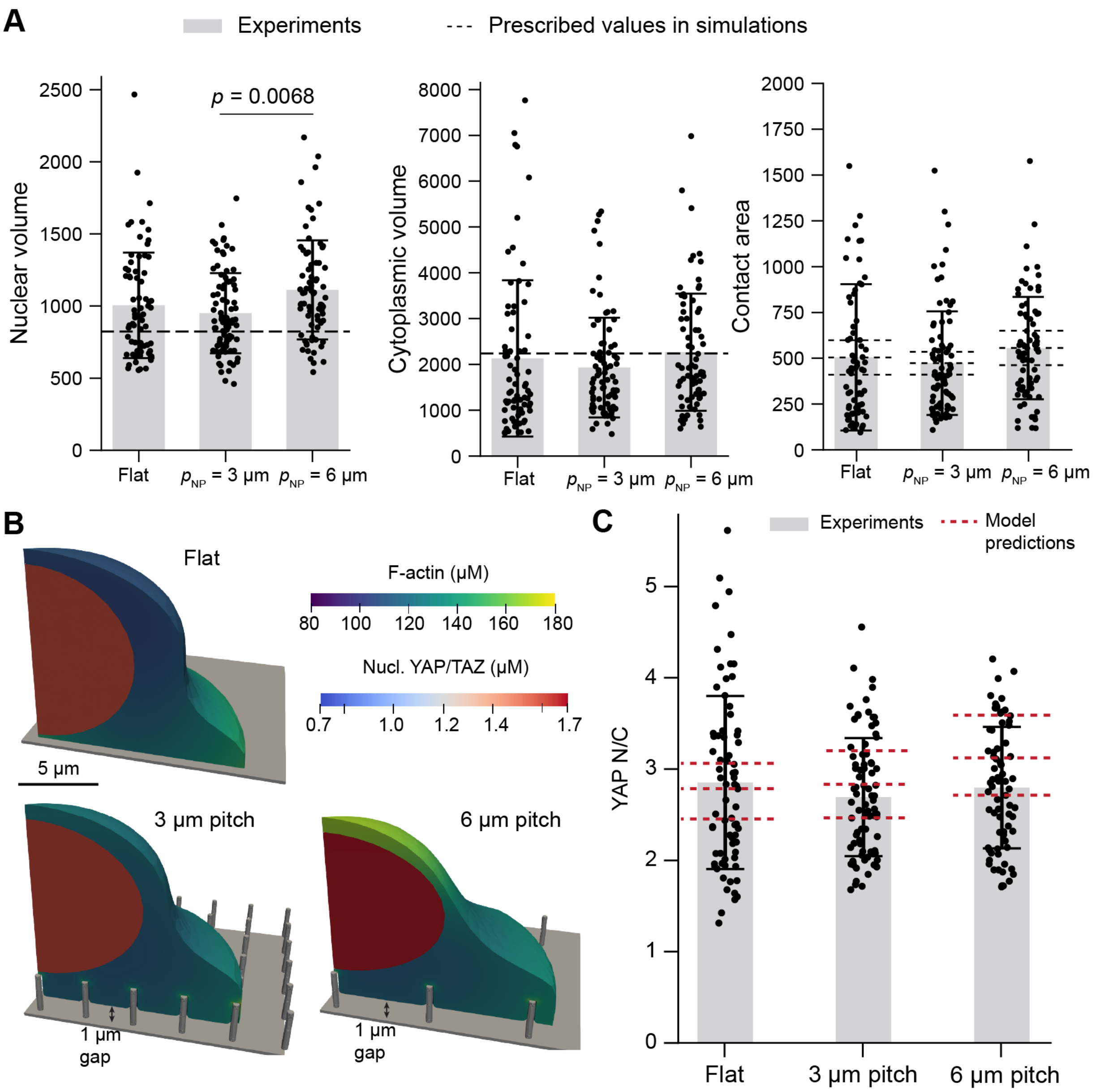


**Figure S7: Experimentally informed geometries show similar levels of nuclear YAP/TAZ on flat and nanopillar substrates.** A) Nuclear volume, cellular volume, and cell contact areas were chosen to match values measured experimentally (Flat $N = 72$ cells, 3 µm pitch $N = 84$ cells, 6 µm pitch $N = 75$ cells). A smaller, but realistic, nuclear volume was chosen to ensure the nucleus fit within the cell boundaries. Three contact areas were chosen for each condition to match the average $\pm\ 2 \times$ SEM (approximately the 95% confidence interval) of the projected cell areas in experiments. B) Steady-state predictions for F-actin and nuclear YAP/TAZ in cells with the average contact area for each condition. C) Comparison between experimental measurements of YAP N/C (Flat $N = 72$ cells, 3 µm pitch $N = 84$ cells, 6 µm pitch $N = 75$ cells)and model predictions for YAP/TAZ N/C.

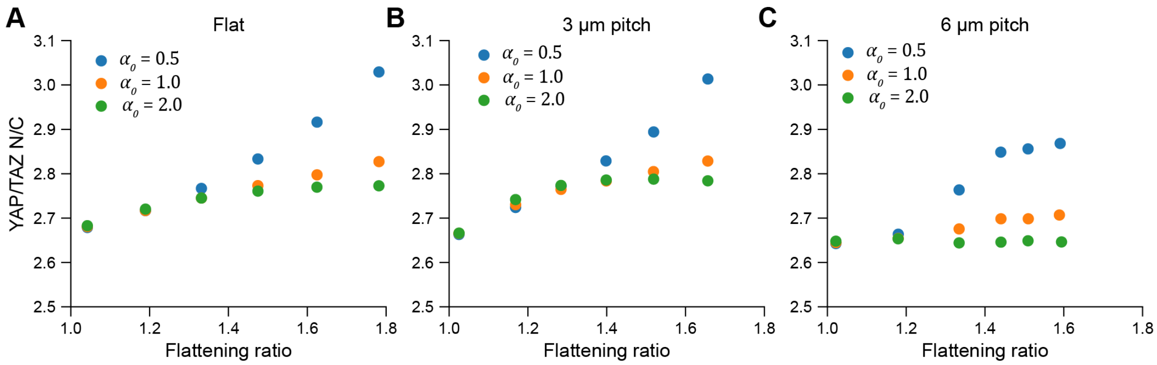


**Figure S8: Correlation between YAP/TAZ and nuclear flattening for different values of stretch sensitivity.** A-C) The correlation between nuclear flattening and YAP/TAZ N/C is shown for 3 different choices of the stretch sensitivity parameter $\alpha_0$. As shown, most cases result in a positive correlation, but the slope is determined by $\alpha_0$. Out of the tested options, $\alpha_0 = 0.5$ provides the best match to experiments.

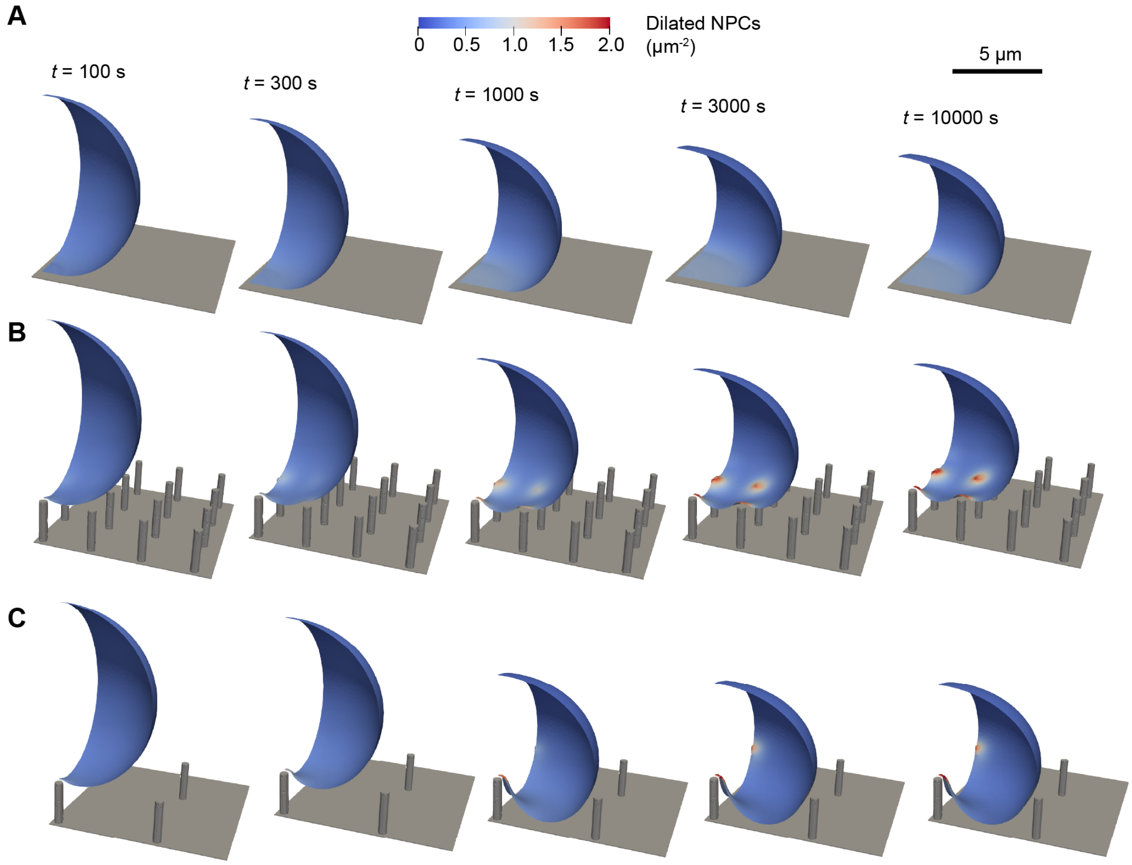


**Figure S9: Simulated NPC dilation occurs locally around nanopillars during nuclear flattening** Select time-points show the local density of dilated NPCs within the nucleus in cells on a flat substrate (A), 3 µm pitch nanopillar substrate (B), or 6 µm pitch nanopillar substrate (C).

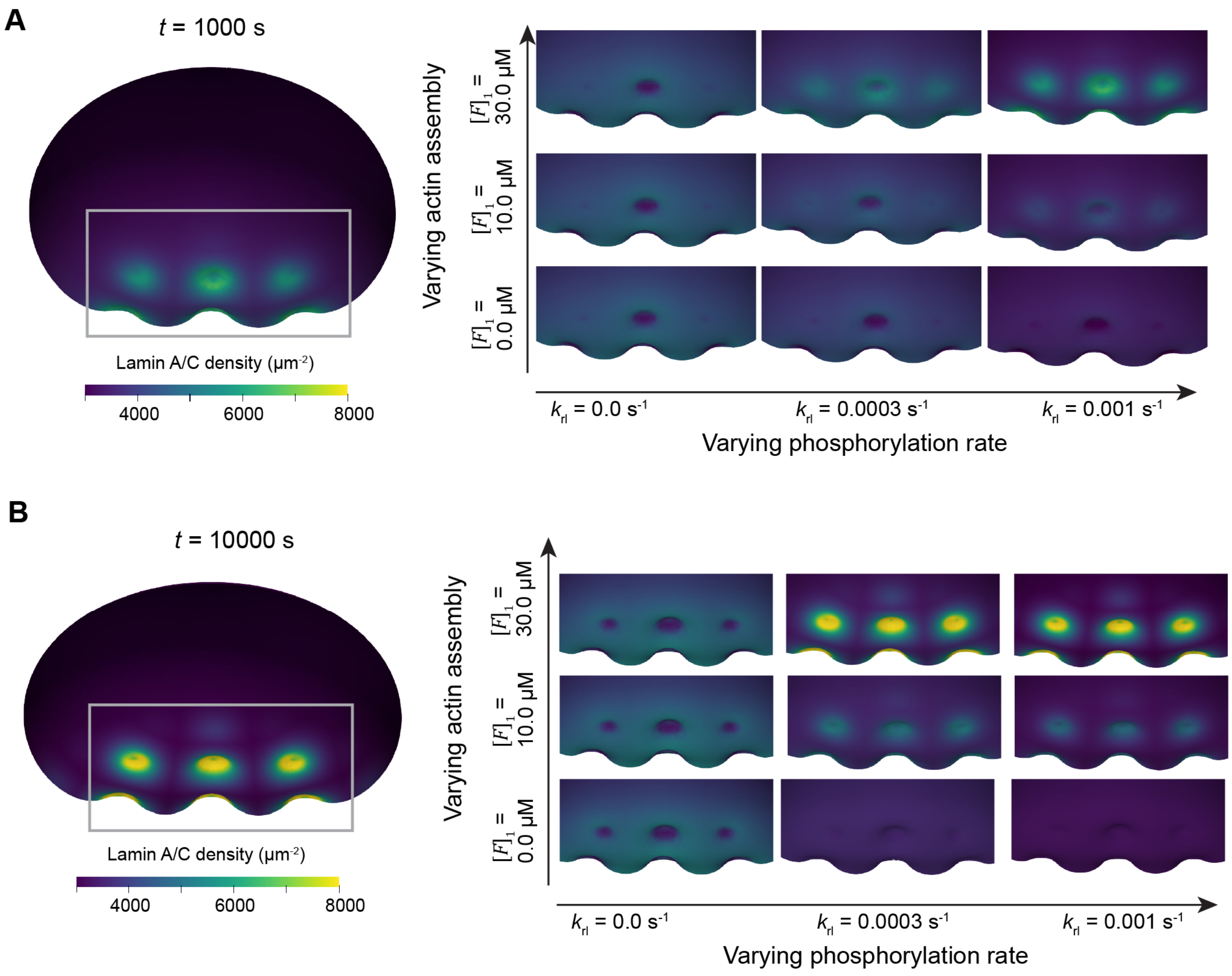


**Figure S10: Lamin localization around nanopillars is sensitive to the rate of lamin phosphorylation and curvature-dependent actin assembly.** Lamin localization is displayed at an intermediate time ($t = 1000$ s, panel A) and the final time ($t = 10000$ s, panel B) for different values of the lamin A/C phosphorylation rate ($k_{\mathrm{fl}}$) and the curvature-dependent actin assembly rate ($[F]_1$).

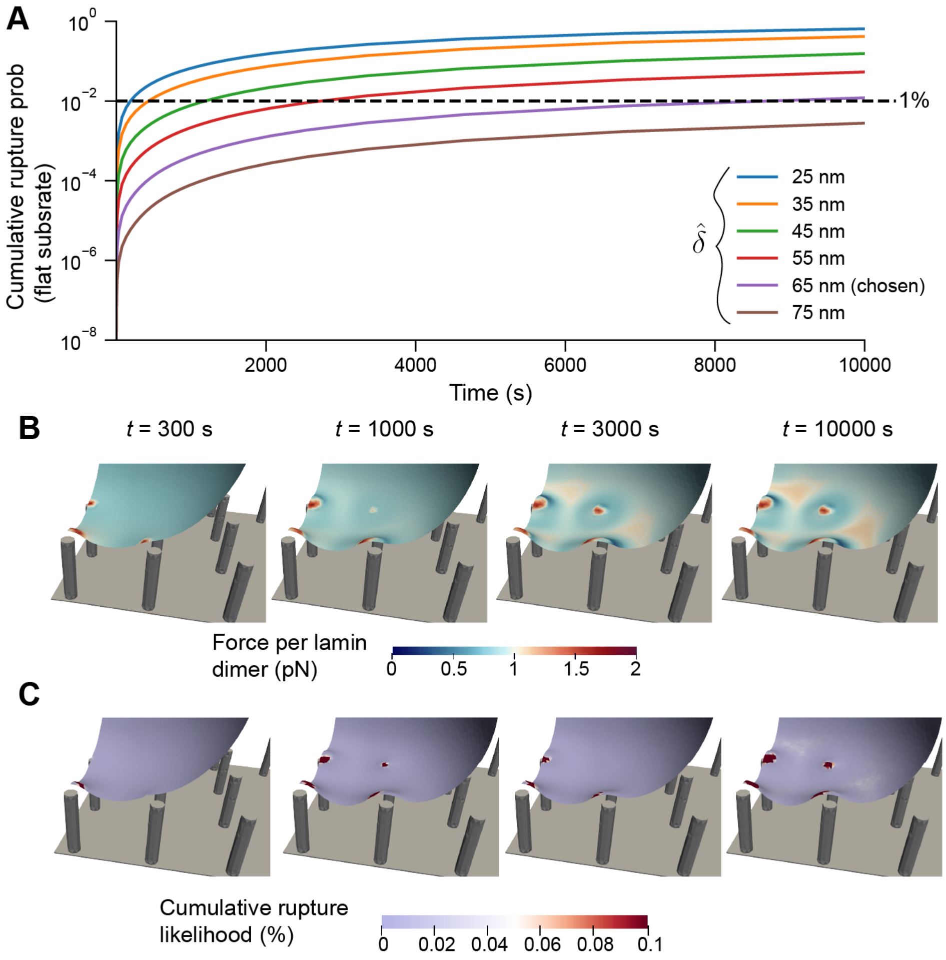


**Figure S11: Process of parameter calibration for rupture likelihood calculations.** A) The value of $\hat{\delta}$ determines rupture likelihood on a flat substrate. $k_{\text{rupture},0}$ was determined separately for each value of $\hat{\delta}$ to ensure an 80% rupture likelihood on 3 µm pitch substrates in each case. For higher values of $\hat{\delta}$, this results in a much lower rupture likelihood on flat substrates, in agreement with experiments. The chosen $\hat{\delta}$ corresponds to a rupture likelihood of about 1% on flat substrates. B-C) Force per lamin dimer (B) and the resulting cumulative rupture likelihood over each element (C) for a nucleus on a 3 micron pitch substrate. Calculations in B-C used $\hat{\delta} = 65$ nm and $k_{\text{rupture},0} = 6.151 \times 10^{-18}$ s$^{-1}$.

**Movie 1:** Lamina stretch ($\alpha$) during nuclear compression with $\sigma_{\text{cap}} = 0 - 1000$ Pa on flat versus nanopillar substrates. This movie shows a flat substrate, followed by substrates with $r_{\text{NP}} = 200$ nm, $h_{\text{NP}} = 2.5$ μm, and $p_{\text{NP}} = 3$ μm and 6 μm.

**Movie 2:** NE tension ($\gamma$) during nuclear compression with $\sigma_{\text{cap}} = 0 - 1000$ Pa on flat versus nanopillar substrates. This movie shows a flat substrate, followed by substrates with $r_{\text{NP}} = 200$ nm, $h_{\text{NP}} = 2.5$ μm, and $p_{\text{NP}} = 3$ μm and 6 μm.

**Movie 3:** NPC dilation on the outer surface of the NE during simulations with $\sigma_{\max} = 100$ Pa versus $\sigma_{\max} = 400$ Pa on flat and nanopillar ($r_{\text{NP}} = 200$ nm, $h_{\text{NP}} = 2.5$ μm, and $p_{\text{NP}} = 3$ μm or 6 μm) substrates.

**Movie 4:** Lamin density over time during nuclear compression by 400 Pa cap stress on 3 μm or 6 μm pitch nanopillar substrate.

**Movie 5:** Lamin density over time during nuclear compression by 400 Pa cap stress on 3 μm or 6 μm pitch nanopillar substrate.

**Movie 6:** Rupture likelihood in low-lamin (70%) versus normal lamin (100%) cells on substrates with $r_{\text{NP}} = 0.45$ μm, $p_{\text{NP}} = 3.5$ μm, $h_{\text{NP}} = 3.25$ μm.

### References


[1] Sreenath Balakrishnan, Suma S Mathad, Geetika Sharma, Shilpa R Raju, Uma B Reddy, Saumitra Das, and G K Ananthasuresh. A nondimensional model reveals alterations in nuclear mechanics upon hepatitis C virus replication. *Biophys. J.*, 116(7):1328–1339, 2 April 2019.

[2] Dong-Hwee Kim, Bo Li, Fangwei Si, Jude M Phillip, Denis Wirtz, and Sean X Sun. Volume regulation and shape bifurcation in the cell nucleus. *J. Cell Sci.*, 128(18):3375–3385, 15 September 2015.

[3] Xuan Cao, Emad Moeendarbary, Philipp Isermann, Patricia M Davidson, Xiao Wang, Michelle B Chen, Anya K Burkart, Jan Lammerding, Roger D Kamm, and Vivek B Shenoy. A chemomechanical model for nuclear morphology and stresses during cell transendothelial migration. *Biophys. J.*, 111(7):1541–1552, 4 October 2016.

[4] Larry A Taber. *Nonlinear theory of elasticity: Applications in biomechanics: Applications in biomechanics.* World Scientific Publishing, Singapore, Singapore, 23 February 2004.

[5] Dan Deviri and Samuel A Safran. Balance of osmotic pressures determines the nuclear-to-cytoplasmic volume ratio of the cell. *Proc. Natl. Acad. Sci. U. S. A.*, 119(21):e2118301119, 24 May 2022.

[6] Stefan Stöberl, Johannes Flommersfeld, Maximilian M Kreft, Martin Benoit, Chase P Broedersz, and Joachim O Rädler. Nuclear deformation and dynamics of migrating cells in 3D confinement reveal adaptation of pulling and pushing forces. *Sci. Adv.*, 10(34):eadm9195, 23 August 2024.

[7] Britta Koch, Samuel Sanchez, Christine K Schmidt, Anka Swiersy, Stephen P Jackson, and Oliver G Schmidt. Confinement and deformation of single cells and their nuclei inside size-adapted microtubes. *Adv. Healthc. Mater.*, 3(11):1753–1758, November 2014.

[8] Chloe M Funkhouser, Rastko Sknepnek, Takeshi Shimi, Anne E Goldman, Robert D Goldman, and Monica Olvera de la Cruz. Mechanical model of blebbing in nuclear lamin meshworks. *Proc. Natl. Acad. Sci. U. S. A.*, 110(9):3248–3253, 26 February 2013.

[9] Newsha Koushki, Ajinkya Ghagre, Luv Kishore Srivastava, Clayton Molter, and Allen J Ehrlicher. Nuclear compression regulates YAP spatiotemporal fluctuations in living cells. *Proc. Natl. Acad. Sci. U. S. A.*, 120(28):e2301285120, 11 July 2023.

[10] Amy J Prunuske and Katharine S Ullman. The nuclear envelope: form and reformation. *Curr. Opin. Cell Biol.*, 18(1):108–116, February 2006.

[11] Anand P Singh, Rémi Galland, Megan L Finch-Edmondson, Gianluca Grenci, Jean-Baptiste Sibarita, Vincent Studer, Virgile Viasnoff, and Timothy E Saunders. 3D protein dynamics in the cell nucleus. *Biophys. J.*, 112(1):133–142, 10 January 2017.

[12] Nil Ege, Anna M Dowbaj, Ming Jiang, Michael Howell, Steven Hooper, Charles Foster, Robert P Jenkins, and Erik Sahai. Quantitative analysis reveals that actin and src-family kinases regulate nuclear YAP1 and its export. *Cell Syst.*, 6(6):692–708.e13, 27 June 2018.

[13] Linda Truebestein, Daniel J Elsner, Elisabeth Fuchs, and Thomas A Leonard. A molecular ruler regulates cytoskeletal remodelling by the rho kinases. *Nat. Commun.*, 6(1):10029, 1 December 2015.

[14] Emmet A Francis, Einollah Sarikhani, Vrund Patel, Dhivya Pushpa Meganathan, Leah Sadr, Anum Tahir, Zeinab Jahed, and Padmini Rangamani. Nanoscale curvature regulates YAP/TAZ nuclear localization through nuclear deformation and rupture. *Adv. Sci. (Weinh.)*, page e2415029, 3 June 2025.

[15] Xiao Li, Lasse H Klausen, Wei Zhang, Zeinab Jahed, Ching-Ting Tsai, Thomas L Li, and Bianxiao Cui. Nanoscale surface topography reduces focal adhesions and cell stiffness by enhancing integrin endocytosis. *Nano Lett.*, 21(19):8518–8526, 13 October 2021.

[16] Wei Zhang, Chih-Hao Lu, Melissa L Nakamoto, Ching-Ting Tsai, Anish R Roy, Christina E Lee, Yang Yang, Zeinab Jahed, Xiao Li, and Bianxiao Cui. Curved adhesions mediate cell attachment to soft matrix fibres in three dimensions. *Nat. Cell Biol.*, 25(10):1453–1464, October 2023.

[17] J L Broers, B M Machiels, G J van Eys, H J Kuijpers, E M Manders, R van Driel, and F C Ramaekers. Dynamics of the nuclear lamina as monitored by GFP-tagged a-type lamins. *J. Cell Sci.*, 112 ( Pt 20)(20):3463–3475, 15 October 1999.

[18] Kiersten Elizabeth Scott, Stephanie I Fraley, and Padmini Rangamani. A spatial model of YAP/TAZ signaling reveals how stiffness, dimensionality, and shape contribute to emergent outcomes. *Proc. Natl. Acad. Sci. U. S. A.*, 118(20):e2021571118, 18 May 2021.

[19] Abigail Buchwalter. Intermediate, but not average: The unusual lives of the nuclear lamin proteins. *Curr. Opin. Cell Biol.*, 84(102220):102220, 22 October 2023.

[20] Shuaiyu Liu, Fangyuan Xiong, Zhen Dou, Lingluo Chu, Yihan Yao, Ming Wang, Xuebiao Yao, Xing Liu, and Zhikai Wang. Phosphorylation of lamin a/C regulates the structural integrity of the nuclear envelope. *J. Biol. Chem.*, 301(1):108033, 1 January 2025.

[21] N Daigle, J Beaudouin, L Hartnell, G Imreh, E Hallberg, J Lippincott-Schwartz, and J Ellenberg. Nuclear pore complexes form immobile networks and have a very low turnover in live mammalian cells. *J. Cell Biol.*, 154(1):71–84, 9 July 2001.

[22] Luv Kishore Srivastava, Zhaoping Ju, Ajinkya Ghagre, and Allen J Ehrlicher. Spatial distribution of lamin a/C determines nuclear stiffness and stress-mediated deformation. *J. Cell Sci.*, 134(10):jcs248559, 15 May 2021.

[23] K Tanuj Sapra, Zhao Qin, Anna Dubrovsky-Gaupp, Ueli Aebi, Daniel J Müller, Markus J Buehler, and Ohad Medalia. Nonlinear mechanics of lamin filaments and the meshwork topology build an emergent nuclear lamina. *Nat. Commun.*, 11(1):6205, 4 December 2020.

[24] Kexin Zhu, Xiangfu Guo, Aravind Chandrasekaran, Xinwen Miao, Padmini Rangamani, Wenting Zhao, and Yansong Miao. Membrane curvature initiates Cdc42-FBP17-N-WASP clustering and actin nucleation. *EMBO J.*, 45(3):953–986, 3 February 2026.

[25] Takeshi Shimi, Katrin Pfleghaar, Shin-Ichiro Kojima, Chan-Gi Pack, Irina Solovei, Anne E Goldman, Stephen A Adam, Dale K Shumaker, Masataka Kinjo, Thomas Cremer, and Robert D Goldman. The a- and B-type nuclear lamin networks: microdomains involved in chromatin organization and transcription. *Genes Dev.*, 22(24):3409–3421, 15 December 2008.

[26] Amnon Buxboim, Joe Swift, Jerome Irianto, Kyle R Spinler, P C Dave P Dingal, Avathamsa Athirasala, Yun-Ruei C Kao, Sangkyun Cho, Takamasa Harada, Jae-Won Shin, and Dennis E Discher. Matrix elasticity regulates lamin-a,C phosphorylation and turnover with feedback to actomyosin. *Curr. Biol.*, 24(16):1909–1917, 18 August 2014.

[27] Joe Swift, Irena L Ivanovska, Amnon Buxboim, Takamasa Harada, P C Dave P Dingal, Joel Pinter, J David Pajerowski, Kyle R Spinler, Jae-Won Shin, Manorama Tewari, Florian Rehfeldt, David W Speicher, and Dennis E Discher. Nuclear lamin-A scales with tissue stiffness and enhances matrix-directed differentiation. *Science*, 341(6149):1240104, 30 August 2013.

[28] M L Gardel, J H Shin, F C MacKintosh, L Mahadevan, P Matsudaira, and D A Weitz. Elastic behavior of cross-linked and bundled actin networks. *Science*, 304(5675):1301–1305, 28 May 2004.

[29] Ted Belytschko, Kam Liu, Brian Moran, and Khalil Elkhodary. *Nonlinear finite elements for continua and structures*. John Wiley & Sons, Nashville, TN, 2 edition, 27 December 2013.

[30] Thomas J R Hughes. *The Finite Element Method*. Dover Publications, 16 August 2000.

[31] Marc Herant and Micah Dembo. Cytopede: a three-dimensional tool for modeling cell motility on a flat surface. *J. Comput. Biol.*, 17(12):1639–1677, December 2010.

[32] D Lanser and J G Verwer. Analysis of operator splitting for advection–diffusion–reaction problems from air pollution modelling. *J. Comput. Appl. Math.*, 111(1-2):201–216, 15 November 1999.

[33] Ersin Bahar and Gürhan Gürarslan. Numerical solution of advection-diffusion equation using operator splitting method. *Int. J. Eng. Appl. Sci.*, 9(4):76–88, 27 December 2017.

[34] Nima Mostafazadeh and Zhangli Peng. Microstructure-based nuclear lamina constitutive model. *Cytoskeleton (Hoboken)*, 81(8):297–309, 1 August 2024.

[35] Einollah Sarikhani, Vrund Patel, Zhi Li, Dhivya Pushpa Meganathan, Keivan Rahmani, Leah Sadr, Ryan Hosseini, Diether Visda, Shivani Shukla, Hamed Naghsh-Nilchi, Adarsh Balaji, Gillian McMahon, Shaoming Chen, Johannes Schöneberg, Colleen A McHugh, Lingyan Shi, and Zeinab Jahed. Engineered nanotopographies induce transient openings in the nuclear membrane. *Adv. Funct. Mater.*, 35(6):2410035, 5 February 2025.

[36] Adam Engler, Lucie Bacakova, Cynthia Newman, Alina Hategan, Maureen Griffin, and Dennis Discher. Substrate compliance versus ligand density in cell on gel responses. *Biophys. J.*, 86(1 Pt 1):617–628, January 2004.

[37] Lindsey Hanson, Ziliang Carter Lin, Chong Xie, Yi Cui, and Bianxiao Cui. Characterization of the cell-nanopillar interface by transmission electron microscopy. *Nano Lett.*, 12(11):5815–5820, 14 November 2012.

[38] Jörg Albuschies and Viola Vogel. The role of filopodia in the recognition of nanotopographies. *Sci. Rep.*, 3(1):1658, 15 April 2013.

[39] Gil-Sung Kim, Dong-Joo Kim, Jung-Hwan Hyung, Myung Kyu Lee, and Sang-Kwon Lee. Dependence of filopodia morphology and the separation efficiency of primary CD4+ T-lymphocytes on nanopillars. *Anal. Chem.*, 86(11):5330–5337, 3 June 2014.

[40] Zhi Li, Einollah Sarikhani, Sirasit Prayotamornkul, Dhivya Pushpa Meganathan, Zeinab Jahed, and Lingyan Shi. Multimodal imaging unveils the impact of nanotopography on cellular metabolic activities. *Chem. Biomed. Imaging*, 2(12):825–834, 23 December 2024.

[41] Einollah Sarikhani, Dhivya Pushpa Meganathan, Anne-Kathrine Kure Larsen, Keivan Rahmani, Ching-Ting Tsai, Chih-Hao Lu, Abel Marquez-Serrano, Leah Sadr, Xiao Li, Mingdong Dong, Francesca Santoro, Bianxiao Cui, Lasse Hyldgaard Klausen, and Zeinab Jahed. Engineering the cellular microenvironment: Integrating three-dimensional nontopographical and two-dimensional biochemical cues for precise control of cellular behavior. *ACS Nano*, 18(29):19064–19076, 23 July 2024.
